# Pre-metazoan origin of neuropeptide signalling

**DOI:** 10.1101/2021.11.19.469228

**Authors:** Yañez-Guerra Luis Alfonso, Thiel Daniel, Jékely Gáspár

## Abstract

Neuropeptides are a diverse class of signalling molecules in metazoans. They occur in all animals with a nervous system and also in neuron-less placozoans. However, their origin has remained unclear because no neuropeptide shows deep homology across lineages and none have been found in sponges. Here, we identify two neuropeptide precursors, phoenixin and nesfatin, with broad evolutionary conservation. By database searches, sequence alignments and gene-structure comparisons we show that both precursors are present in bilaterians, cnidarians, ctenophores and sponges. We also found phoenixin and a secreted nesfatin precursor homolog in the choanoflagellate *Salpingoeca rosetta*. Phoenixin in particular, is highly conserved, including its cleavage sites, suggesting that prohormone processing occurs also in choanoflagellates. In addition, based on phyletic patterns and negative pharmacological assays we question the originally proposed GPR-173 (SREB3) as a phoenixin receptor. Our findings indicate that signalling by secreted neuropeptide homologs has pre-metazoan origins and thus evolved before neurons.

## Introduction

Neuropeptides are one of the largest families of neuronal signalling molecules. They are derived from precursor molecules (proneuropeptides) that must undergo processing to release active mature peptides (Veenstra 2000). Neuropeptides play pivotal roles in the regulation of different biological processes such as feeding, cognition and reproduction, and they have been extensively studied in bilaterians (Jékely 2013; Mirabeau and Joly 2013; Elphick et al. 2018; Thiel et al. 2021). Neuropeptide-like molecules have also been described in cnidarians, ctenophores and in the neuron-less placozoan *Trichoplax adhaerens* (Jékely 2013; Senatore et al. 2017; Varoqueaux et al. 2018; Koch and Grimmelikhuijzen 2020; Burkhardt and Jékely 2021; Sachkova et al. 2021). So far, none of the neuropeptides described in these species have shown enough similarity to be considered homologues of bilaterian short neuropeptides. Only potential ctenophore homologues of bilaterian large cysteine-rich hormones including trunk-like proteins and prothoracicotropic hormone (PTTH) have been identified (de Oliveira et al. 2019). In addition, sponges have cystine-knot-family growth factors (CKGF), related to bilaterian glycoprotein hormones (Roch and Sherwood 2014). Despite the lack of deep conservation in neuropeptides, the machinery involved in precursor processing had evolved before metazoans. Homologues of the enzymes peptidyl-glycine α-amidating monooxygenase (PAM), an enzyme important for the amidation of neuropeptides in Bilateria, and prohormone convertases, important for the cleavage of proneuropeptides exist in the green algae *Chlamydomonas reinhardtii* and other ciliated protists (Kumar et al. 2016; Kumar et al. 2017; Luxmi et al. 2018; Luxmi et al. 2019). In *C. reindhardtii*, several amidated peptides have been identified and some have signalling functions during gamete chemotaxis (Luxmi et al. 2019). These results suggest that neuropeptide signalling in metazoans has a deep evolutionary ancestry in single-celled eukaryotes. However, the *C. reindhardtii* peptides show no similarity to any metazoans neuropeptides and it remains unclear when animal neuropeptides evolved.

Here we report two neuropeptide precursor sequences of pre-metazoan origin – the phoenixin and nesfatin precursors – with orthologs in all major metazoan branches as well as choanoflagellates. Phoenixin (PNX) was first identified by screening the human genome database (Yosten et al. 2013). The mature PNX peptide derives from a precursor named small integral membrane protein 20 (SMIM20). SMIM20 is highly conserved across vertebrates and contains a signal peptide, an amidation motif (Gly) and dibasic cleavage sites (Yosten et al. 2013). In mammals, SMIM20 undergoes post-translational processing to produce two C-terminally amidated peptides, phoenixin-14 (PNX-14) and phoenixin-20 (PNX-20) with PNX-14 being the most abundant peptide in rodent tissues (Lyu et al. 2013; Yosten et al. 2013). Some vertebrate phoenixin precursors lack the amidation signature in the C-terminal PNX region, including the *Xenopus*, *Silurana*, zebrafish and fugu precursors (Yosten et al. 2013). Recently, neuropeptidomic searches in arthropod transcriptomes revealed phoenixin precursors with conserved cleavage sites to produce the mature PNX peptide PNX14 and PNX-20 (Nguyen et al. 2018). PNX is involved in many different physiological functions in mammals, such as cardio-modulation, memory, anxiety, food intake and reproduction (Clarke and Dhillo 2019; Schalla and Stengel 2019; Billert et al. 2020; Haddock et al. 2020; Ma et al. 2020; Schalla et al. 2020; Friedrich and Stengel 2021; Yao et al. 2021). The mechanism of signalling of PNX, an orphan ligand, remains unclear. However, the GPR173, also known as Super-conserved Receptor Expressed on Brain 3 (SREB3) has been proposed as a potential receptor of PNX, based on a “Deductive Reasoning Strategy” (Stein et al. 2016; Yosten et al. 2021).

Nesfatin-1 (**N**ucleobindin-2-**E**ncoded **S**atiety and **FAT**-**I**nfluencing protei**N**-1) is a neuropeptide identified in 2006 as an 82 amino acid peptide located in the N-terminal region of the protein nucleobindin-2 (NUCB2) (Oh-I et al. 2006). NUCB precursors contain a signal peptide, dibasic cleavage sites, leucine-zipper motifs and EF-hand motifs. It has been shown that these precursors can act as calcium and DNA binding proteins in addition to neuropeptide precursors (Miura et al. 1992; Miura et al. 1994; Kmiecik et al. 2021). NUCB precursors encode three different potential peptides, known as nesfatin-1, 2 and 3. So far, only nesfatin-1 was shown to have a physiological function (Oh-I et al. 2006; Schalla and Stengel 2018). Different functions of nesfatin-1 have been reported in vertebrates, such as glucose metabolism and the regulation of reproduction, stress and anxiety responses (Schalla and Stengel 2018; Friedrich and Stengel 2021). Its role as a satiety-inducing factor has been widely reported in mammals and fish (Ayada et al. 2015; Sundarrajan et al. 2016; Rupp et al. 2021). NUCB precursors encoding the nesfatin-1 peptide region are also present in invertebrates, including ophiuroid echinoderms and the fruit fly *Drosophila melanogaster* (Otte et al. 1999; Zandawala et al. 2017). So far, no receptor has been identified for the nesfatin-1 peptide (Rupp et al. 2021).

## Results and discussion

### Phoenixin and nesfatin-1 precursors were already present in the last common ancestor of Metazoa and choanoflagellates

In an initial bioinformatic survey, we noticed that nesfatin and phoenixin have a broad phyletic distribution, suggesting that these may be the oldest neuropeptides found so far. To analyse this in more detail, we searched the transcriptome and genome sequences of 45 metazoan species (Supplementary Table 1), two choanoflagellates, one filasterean, and *Tunicaraptor unikontum*, a predatory flagellate belonging to a newly identified animal-related lineage (Tikhonenkov, et al. 2020). By sequence-similarity searches, multiple sequence alignments and gene-structure analyses, we identified homologs of the precursors of phoenixin (SMIM20) and nesfatin-1 (NUCB) across major groups of metazoans including Porifera, Ctenophora, Cnidaria, and in most of the bilaterian species analyzed. Furthermore, a PNX precursor was identified in the choanoflagellate *Salpingoeca rosetta*, and a NUCB homolog was detected in *S. rosetta* and *T. unikontum*. Placozoans lack a PNX precursor and the NUCB precursor in this phylum does not contain a nesfatin-1 peptide region. These findings reveal that these neuropeptide precursors originated in pre-metazoan times.

### The phoenixin precursor is highly conserved in metazoans and choanoflagellates

A multiple-sequence alignment reveals a high degree of conservation of the PNX neuropeptide precursor (Figure 1A). There are widely conserved residues across the length of the PNX precursor, with the C-terminal region – corresponding to the mammalian PNX-14 peptide (Lyu et al. 2013; Yosten et al. 2013) – being the most conserved across all tested species (with IQPGGMKVWSDPFD as the consensus sequence). The regions that contain the dibasic and monobasic cleavage sites for proteolytic processing (Veenstra 2000; Hook et al. 2008) are also well conserved across metazoans and in *S. rosetta* (Figure 1A). The predicted neuropeptides show variation in the C-terminal amidation site across metazoans, similar to the variability in C-terminal amidation that was already described within vertebrates (Yosten et al. 2013). In ecdysozoans, *Tribolium castaneum* has a PNX sequence with an amidation signature while other protostomes do not. Within Ctenophora, the *Mnemiopsis leidyi* peptide is predicted to be amidated, while the *Pukia falcata* PNX lacks the amidation motif. The *Oscarella carmella* and *Amphimedon queenslandica* sponge phoenixin precursors have the amidation site but the choanoflagellate *S. rosetta* precursor lacks it. Overall, an amidation site is missing for the majority of species and many show acidic residue (D,E) instead (Figure 1A). In most other neuropeptide families, the C-terminal amidation of homologous peptides is conserved across species (Mirabeau and Joly 2013; Yañez-Guerra et al. 2020). However, there are some examples where this is not the case, such as galanin, which is amidated in most vertebrates but not in humans (Sobrido-Cameán et al. 2019). This unamidated version of human galanin is, nevertheless, functional (Bersani et al. 1991). Thus, it is possible that the amidation of phoenixin, just like in the case of galanin, is important only in certain species.

**Figure 1:**
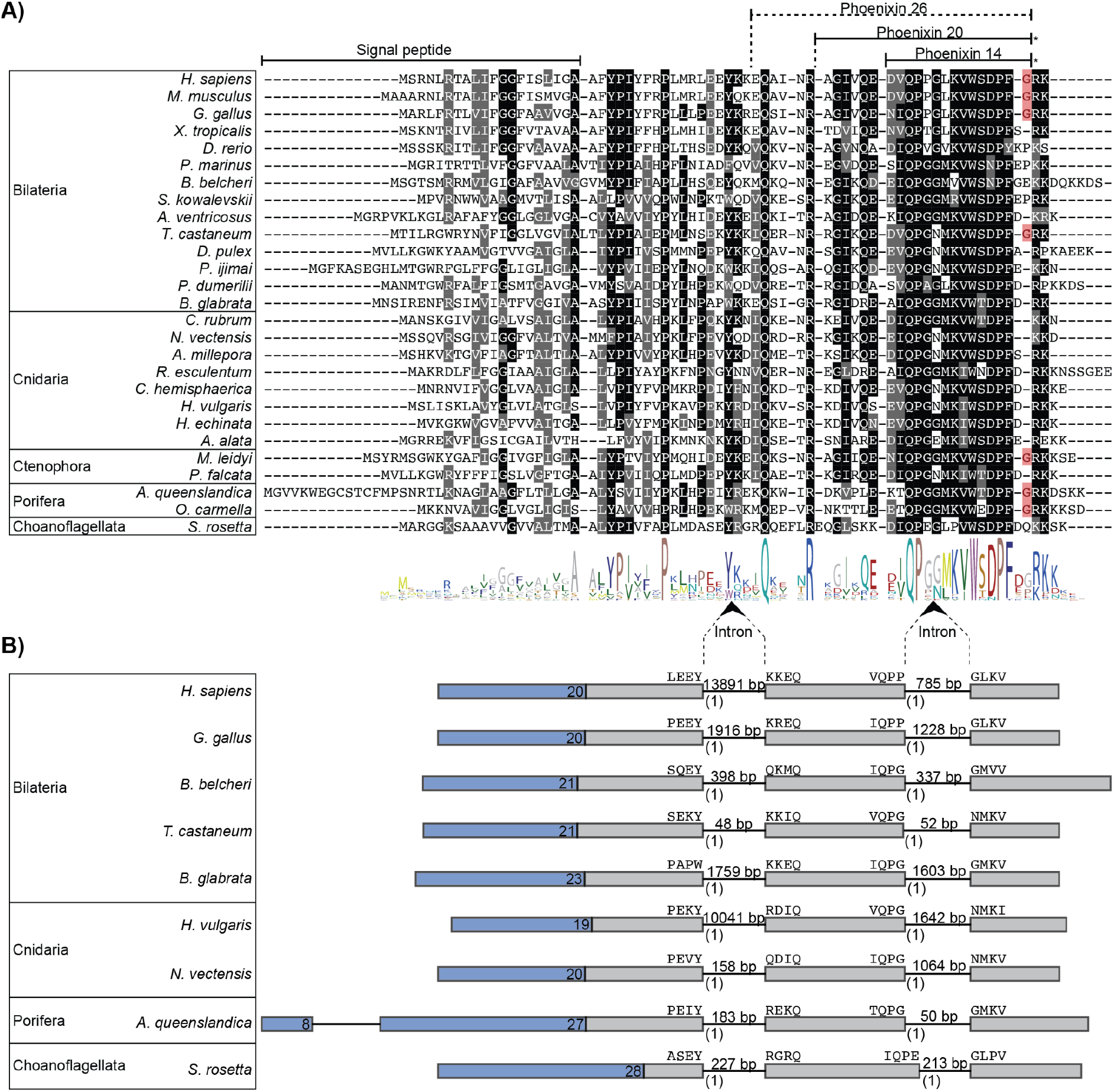
Sequence alignment and genomic structure of phoenixin precursors. **(A)** Alignment of the SMIM20 precursors encoding the PNX peptides. Predicted signal peptides and mature peptides are indicated with lines. Residues that are conserved in more than 50% of the sequences are shown in black and conservative substitutions are shown in grey. Amidation sites are highlighted in red. **(B)** The genomic exon-intron structure of PNX peptide precursors. Signal peptides are in blue with their length indicated. Amino acids encoded at the exon-intron junctions are shown above the exon boxes. Introns are shown as lines, and their length in base pairs is indicated above. Intron phase is shown below introns in parentheses.

To further test the homology of PNX neuropeptides, we carried out a gene structure analysis of the *Homo sapiens* (Mammalia), *Gallus gallus* (Sauropsida), *Branchiostoma belcheri* (Cephalochordata), *T. castaneum* (Ecdysozoa), *Biomphalaria glabrata* (Lophotrocozoa), *Nematostella vectensis* (Cnidaria), *Hydra vulgaris* (Cnidaria), *A. queenslandica* (Porifera) and *Salpingoeca rosetta* (Choanoflagellata) PNX precursor genes. This revealed a high level of conservation of these precursors at the gene-structure level within and beyond metazoans. Most of these proteins are encoded in 3 exons, divided by two phase-1 introns at homologous positions (Figure 1B). In the sponge *A. queenslandica*, there is an additional intron in the region encoding the signal peptide (Figure 1B). Of note, the signal peptide of these proteins could only be predicted by SignalP3.0 (Bendtsen et al. 2004), and not by SignalP4.1 SignalP5, and SignalP6. These newer versions of the software failed to identify the signal peptide even in the human SMIM20 protein or in the phoenixin precursor in *N. norvegicus* (Nguyen et al. 2018).

### The nesfatin-1 precursor, but not the peptide, shows conservation beyond metazoans

We identified homologous NUCB precursors in bilaterian, cnidarian, placozoan, ctenophore and poriferan species, as well as in choanoflagellates and *Tunicaraptor*. However, the N-terminal region that codes the mature neuropeptide nesfatin-1 is less conserved in sponges (Figure 2 A) and not identifiable in *T. adhaerens*, choanoflagellatas or *Tunicaraptor* (Figure 2 B, Supplementary Figures 2 and 3). These sequences only show conservation in the more C-terminal part of the precursor, with strongest conservation in the part corresponding to the N-terminus of the human nesfatin-3 peptide, which contains an EF-hand domain (Supplementary Figure 2). This partial conservation suggests a pre-metazoan origin of the NUCB precursor protein that only evolved the nesfatin-1 peptide in the ancestral metazoan lineage with a potential secondary loss in placozoans. In contrast to the phoenixin-1 precursor gene, the gene structure of the NUCB gene is not widely conserved, and differs already between deuterostome and protostome species in the C-terminal region (Figure 2B). The primary resemblance across all the bilaterian species is that the region encoding the nesfatin-1 peptide is divided by two phase-0 introns. This feature is not found in any non-bilaterian species, except in ctenophores in which a phase-0 intron is present in the region that matches the first phase-0 intron of bilaterians. Thus, the homology of NUCB precursors from bilaterians, non-bilaterian metazoans, choanoflagellates and *Tunicaraptor* was primarily established through sequence-similarity, reciprocal blast and alignment of the entire precursors. The *T. adhaerens*, choanoflagellate and *Tunicaraptor* sequences show homology only in their C-terminal part and not in the nesfatin-1 region (Supplementary Figures 2 and 3).

**Figure 2:**
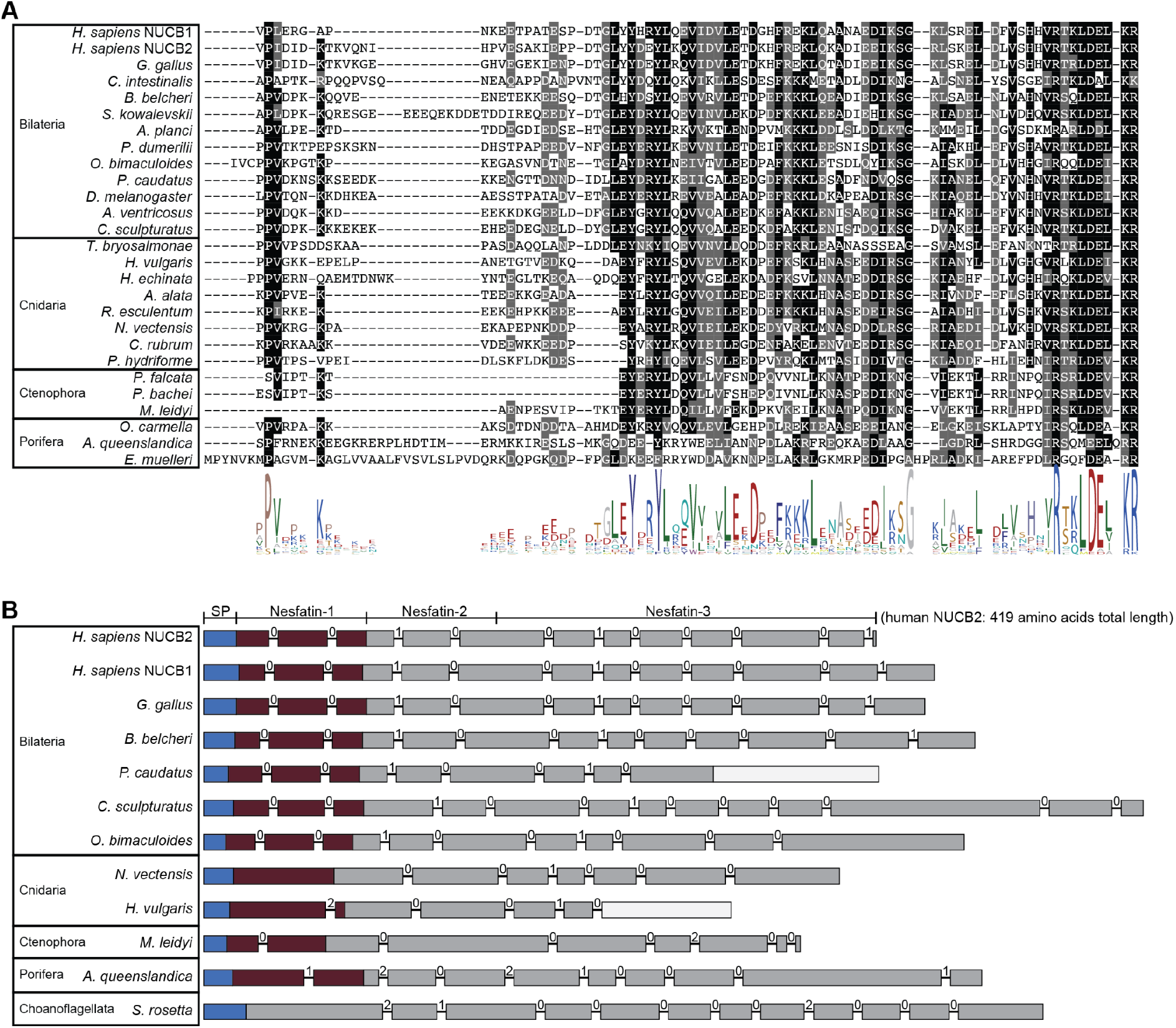
Sequence alignment of Nesfatin-1 and genomic structure of NUCB precursors. **(A)** Alignment of the N-terminal NUCB precursor region encoding the nesfatin-1 peptides. The conserved residues are highlighted, with conservation in more than 50% of sequences shown in black and conservative substitutions shown in grey. **(B)** The genomic exon-intron structure of NUCB precursors. Signal peptides are shown in blue. The nesfatin-1 peptide region is indicated in dark red. Introns are shown as lines, with the phase of the introns shown above the lines. An empty/white box indicates a missing part in the mRNA-genome alignment.

### GPR173 is unlikely to be the phoenixin receptor

The SREB (Super-conserved Receptors Expressed on Brain) family of receptors was named after their expression in the central nervous system with an exceptionally high level of conservation between vertebrate species (Matsumoto et al. 2000; Breton et al. 2021). There are at least three SREB receptors in vertebrates SREB1 (GPR27), SREB2 (GPR85) and SREB3 (GPR173) (Matsumoto et al. 2000). These receptors are orphans, as no ligand-receptor assay has so far identified a potent ligand for them. Based on a “Deductive reasoning strategy” SREB3 has been proposed as a potential receptor for the neuropeptide PNX (Stein et al. 2016). Some *in vivo* experiments further indicate, although indirectly, a ligand-receptor relationship between PNX and SREB3/GPR173. In female rats, exogenously administered PNX induces a pre-ovulatory-like secretion of luteinising hormone (LH). When GPR173 expression was reduced by siRNA treatment, this effect of PNX on LH secretion was significantly reduced (Stein et al. 2016). Furthermore, the siRNA knock-down of GPR173 doubled the length of the oestrous cycle in female rats (Stein et al. 2016) similar to the knock-down of PNX that increased the oestrous cycle by >50% (Yosten et al. 2013).

To further explore whether SREBs are PNX receptors, we searched for homologues of SREB receptors across animals. An initial cluster-based analysis showed that the SREB receptors form a tight cluster, indicating high levels of conservation (Supplementary Figure 4), as previously shown (Matsumoto et al. 2000). The only connection of SREBs to any other GPCR cluster is to monoaminergic receptors (at an e-value of 1e-27). Therefore, we used the SREB cluster with monoamine receptors as outgroups, to carry out a phylogenetic analysis.

SREB receptors have been described as vertebrate-specific, as they have not been identified in non-vertebrate chordates or in invertebrates in previous studies (Matsumoto et al. 2000; Breton et al. 2021). By searching an expanded group of species, we found that the SREB receptors are present in one copy in several invertebrates, including cephalochordates, ambulacrarians, ecdysozoans, and lophotrochozoans. This demonstrates that SREB receptors are of urbilaterian origin, and the three copies of SREBs in vertebrates likely originated during vertebrate evolution (Supplementary Figure 5).

We could not identify SREB receptors in any of the non-bilaterian species, contrasting with the much broader phyletic distribution of the PNX precursors. This indicates that PNX peptides in these organisms must signal via other types of receptors. The phyletic mismatch between PNX and SREB also casts doubt on the suggested ligand-receptor relationship between human PNX and SREB3. Most other GPRC families show tight co-occurrence with their peptide ligands across taxa (Jékely 2013; Mirabeau and Joly 2013)

To directly test if the human PNX-14 peptide is able to activate any of the three human SREB receptors, next we carried out calcium mobilisation assays. We used two different promiscuous chimeric G-proteins, Gqi9 and Gqs5 separately, to test for coupling to different G-alpha subunits. We could not detect any activation of the three SREB receptors by the PNX peptide, even at very high peptide concentrations (up to 1e-4 M; Supplementary Figure 6). In the same assay, we could get reliable activation of two other GPCRs by their cognate peptide ligand. It has to be noted that these types of deorphanisation assays may not work for all ligand-receptor pairs (Foster et al. 2019; Hauser et al. 2020). Nevertheless, the non-matching evolutionary pattern between PNX and SREB (Figure 3) together with the negative receptor activation assay (Supplementary Figure 6) suggest that this receptor-ligand pairing may not be correct and should be re-evaluated.

**Figure 3.**
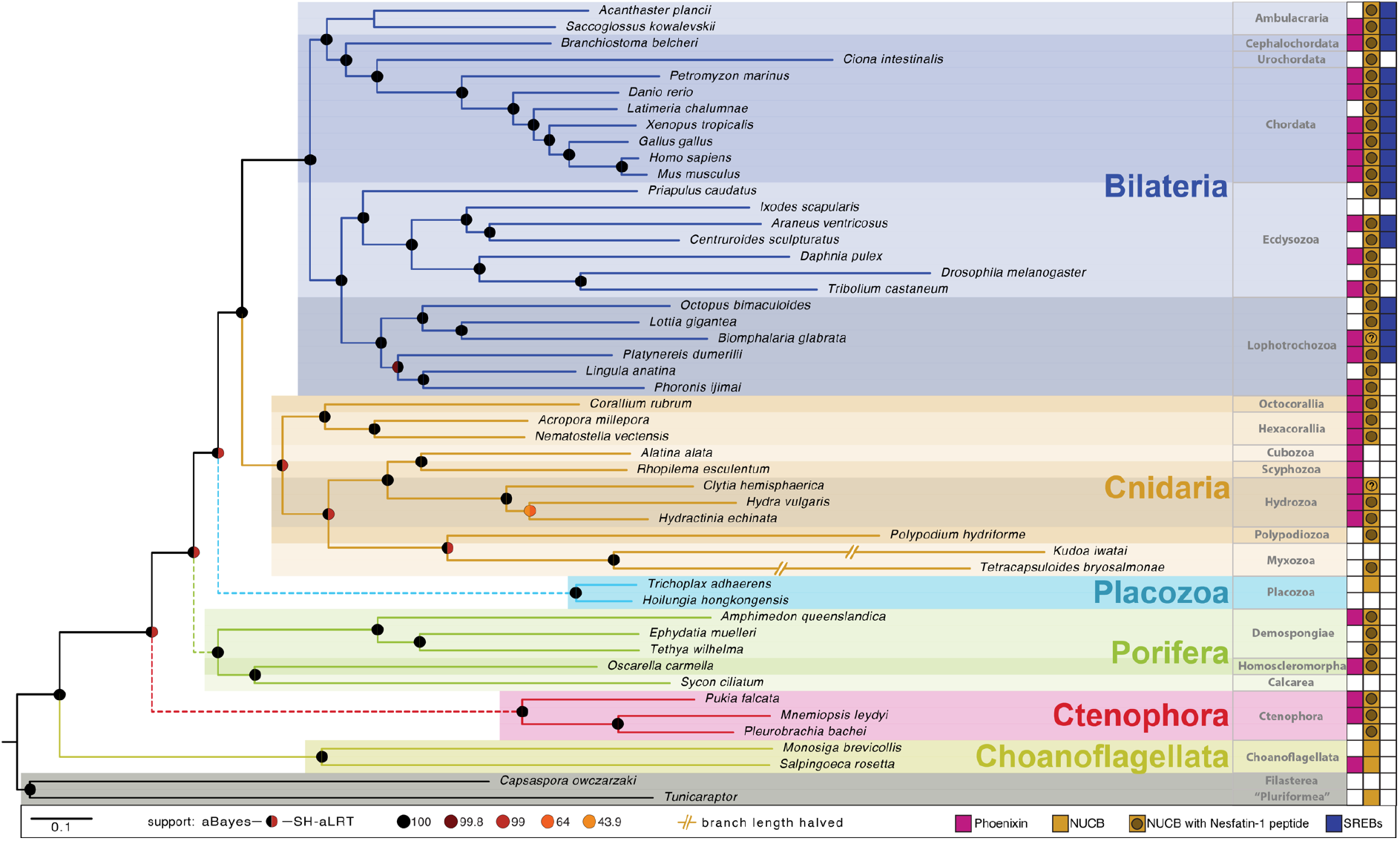
Presence and absence of PNX, nesfatin and SREB receptors in the investigated species. Phylogenomic tree of the investigated species, annotated with the presence/absence of phoenixin, NUCB and its nesfatin-1 peptide, and SREB receptors. The PNX neuropeptide precursor is conserved across metazoans and in the choanoflagellate *S. rosetta*, while GPR173 (proposed as a potential receptor for this peptide) is only present in Bilateria. The NUCB precursor gene is conserved across metazoans and present in choanoflagellates and *Tunicaraptor*, while the nesfatin-1 peptide is only encoded in metazoans. A question mark indicates that the NUCB sequence was only partially recovered with the N-terminal part that encodes the nesfatin-1 peptide missing in the transcriptome. An unfilled box indicates that the corresponding gene was not identified. Dashed lines in main bilaterian branches indicate generally contradicting results in different phylogenomic analyses.

## Conclusions

Our bioinformatic survey identified phoenixin and nesfatin as ancient neuropeptides with pre-metazoan origin. To our knowledge these are the first neuropeptides to be identified in sponges and with a broad conservation across animals, including sponges, ctenophores, cnidarians and bilaterians. The presence of a phoenixin peptide in choanoflagellates demonstrates that some animal neuropeptides have pre-metazoan origin and predate nervous systems. Many other neuronal molecules, including neurosecretory components (Göhde et al. 2021), postsynaptic proteins (Burkhardt et al. 2014), and voltage-gated channel subunits (Moran and Zakon 2014) have a similar history (Burkhardt and Jékely 2021).

What could be the functions of NUCB and phoenixin in choanoflagellates and non-neuronal sponges? The precursor sequences indicate that both are secreted proteins and PNX is processed to release a mature phoenixin peptide in both choanoflagellates and sponges. An interesting possibility is that these proteins regulate feeding. In mammals, nesfatin-1 produces anorexigenic effects, while PNX promotes feeding and drinking behaviour (Maejima et al. 2009; Stengel et al. 2012; Schalla et al. 2017).

We also speculate that PNX and nesfatin-1 may have functionally co-evolved since metazoan origins. This is suggested by their ancestral origins, similar distributions and their complementary roles in some physiological functions in vertebrates. Nesfatin-1 and PNX also broadly coexpress in the rat hypothalamus, with over 70% of the PNX-expressing neurons co-expressing nesfatin-1 (Pałasz et al. 2015). Besides their antagonistic effects on feeding, the two peptides also have opposing roles in the regulation of anxiety and fear-like behaviour. PNX has an anxiolytic effect in mice (Jiang et al. 2015) and likely also humans (Hofmann et al. 2017), while nesfatin-1 increases anxiety (Merali et al. 2008); (Hofmann et al. 2015). In addition, PNX administration leads to increased nesfatin-1-immunoreactivity in rats (Friedrich et al. 2019), indicating a functional interplay.

Overall, our findings suggest that secretion and intercellular signalling by peptides in animal evolution evolved before neurons and synapses, in agreement with the ‘chemical brain’ theory for the origin of nervous systems (Jékely). In future, it will be interesting to test if the two peptides coexpress and have antagonistic functions also in invertebrate nervous systems. Equally exciting will be to explore the function of these precursors in sponges and choanoflagellates. Will the peptides make these organisms anxious or hungry?

## Supplementary Figures

**Supplementary Figure 1.**
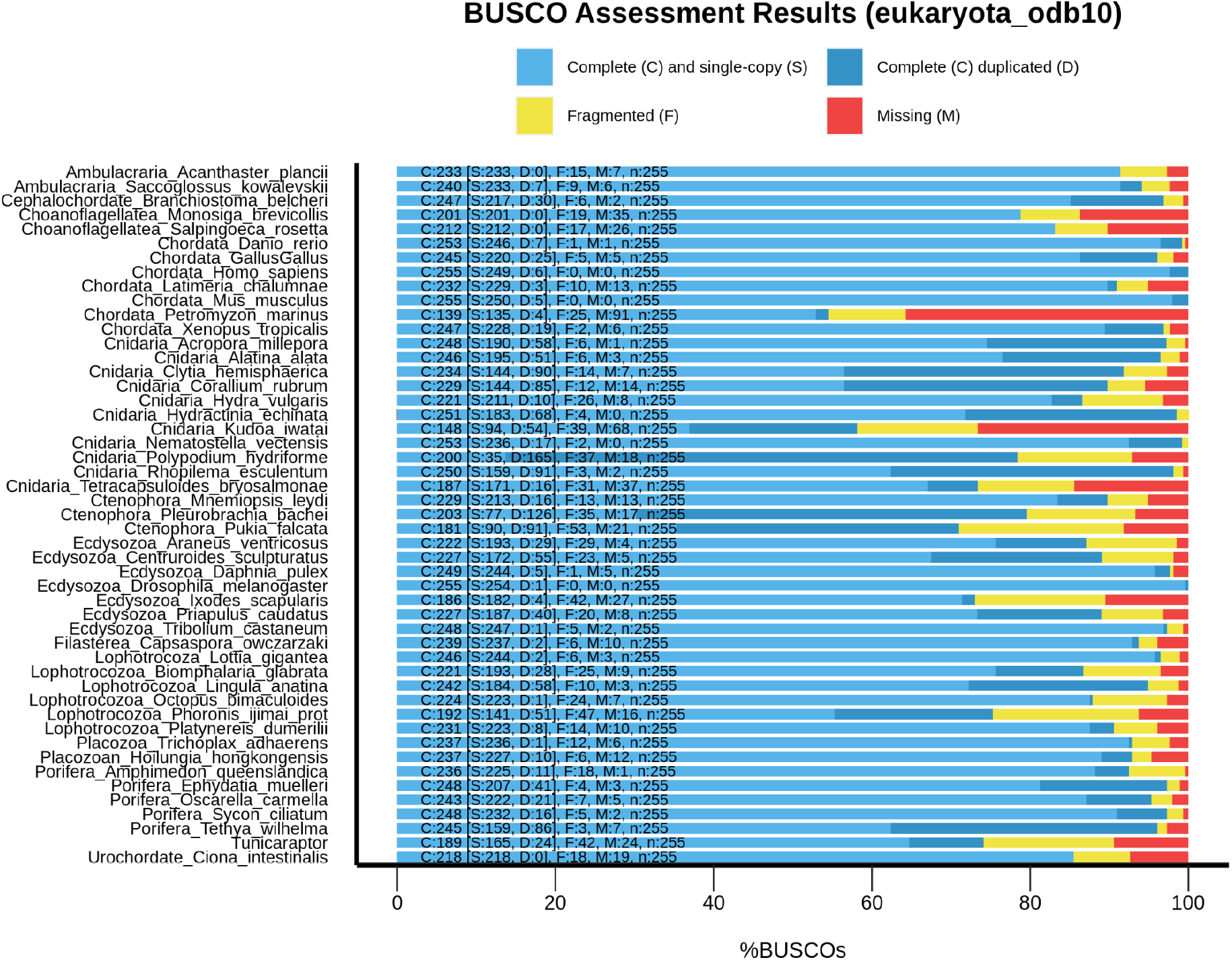
BUSCO completeness analysis. Completeness of the transcriptomes used in the study.

**Supplementary Figure 2.**
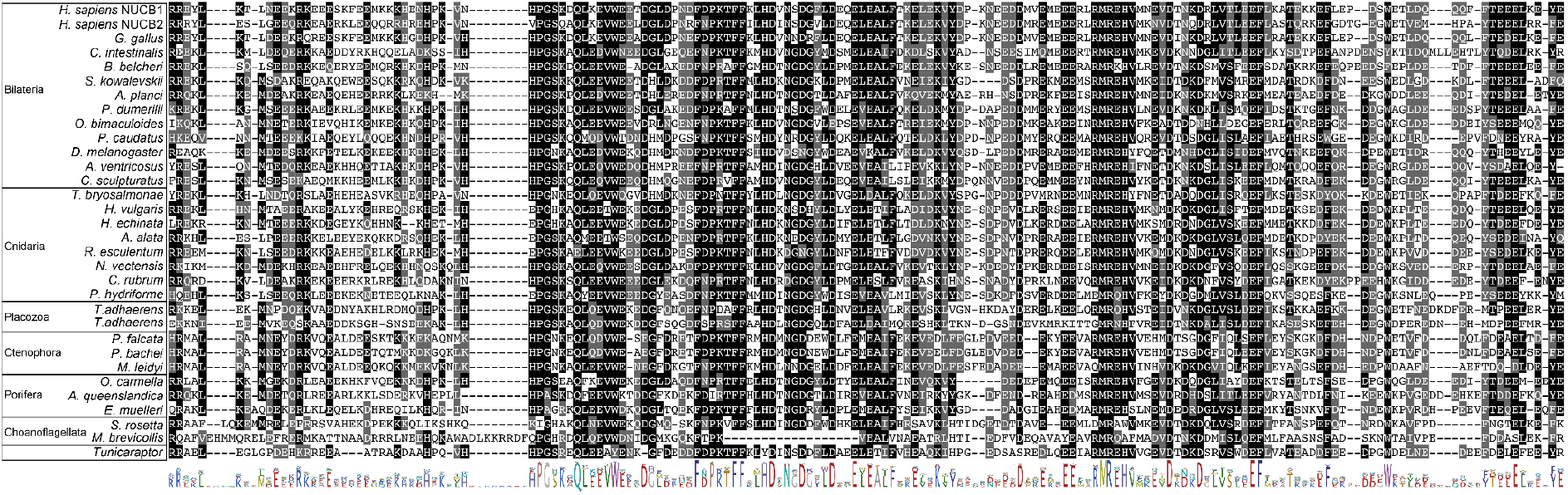
Alignment of the Nesfatin-3 region of NUCB precursors. Alignment of the C-terminal region of the NUCB precursor corresponding to the nesfatin-3 region of the propeptides. The conserved residues are highlighted, with conservation in more than 50% of sequences shown in black and conservative substitutions shown in grey. A consensus sequence is shown below the alignment.

(see supplementary files)

**Supplementary Figure 3. Whole-precursor alignment of NUCB precursors in all the species in which an homolog was identified.** The conserved residues are highlighted, with conservation in more than 50% of sequences shown in black and conservative substitutions shown in grey

**Supplementary Figure 4.**
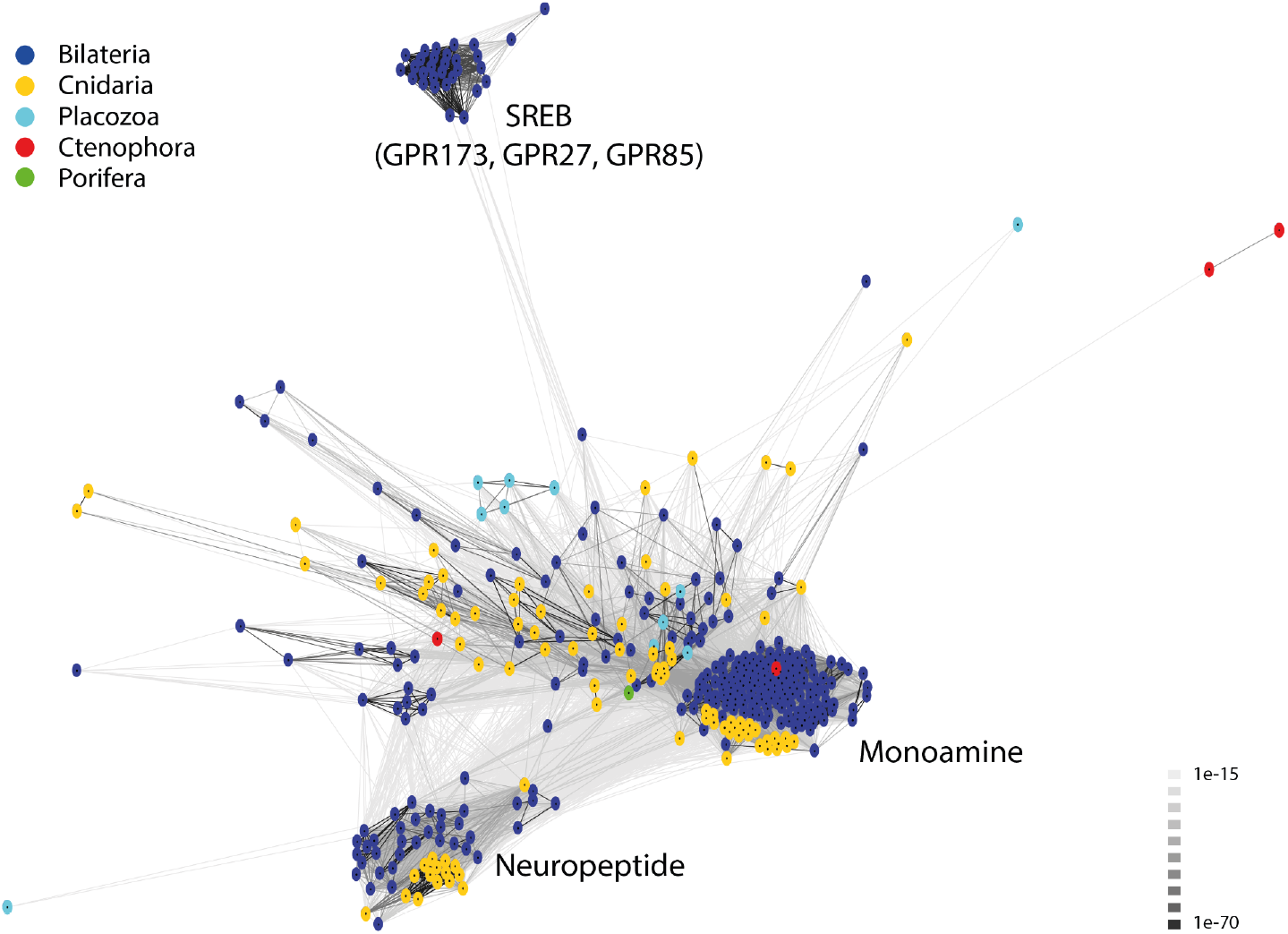
BLOSUM62 cluster map of metazoan receptors families. Nodes correspond to pNPs and are coloured by taxon as indicated on the key. Edges correspond to BLAST connections of p-value >1e-27. The cluster marked as SREB contains the GPR173, GPR27 and GPR85 homologs.

**Supplementary Figure 5.**
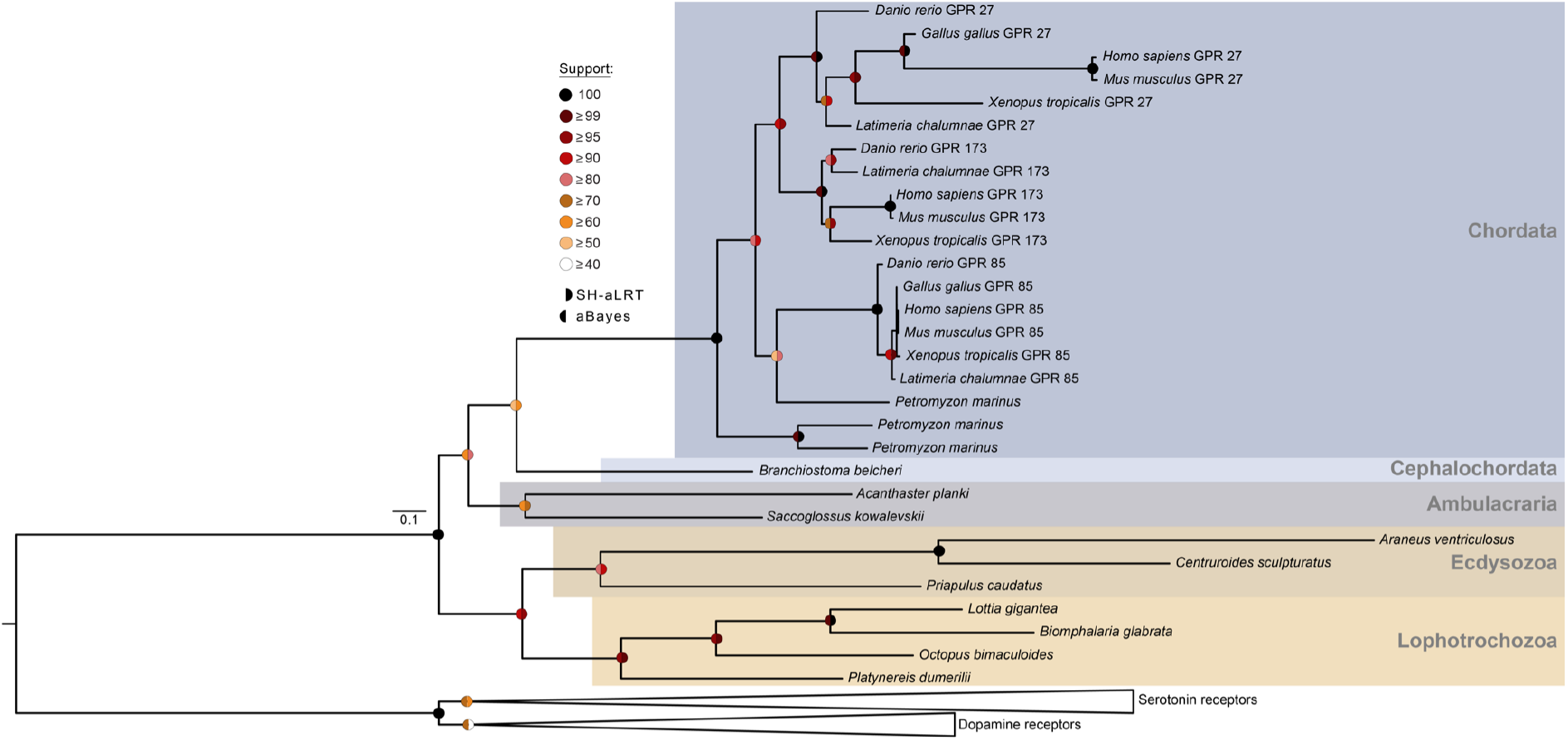
Phylogenetic tree showing the relationship of SREB receptors, including GPR173, GPR27 and GPR85. SREB receptors were identified in cephalochordates, ambulacrarians, ecdysozoans and lophotrocozoans but they are absent in non-bilaterians. The circles represent branch support based on 1000 replicates as explained in the legend, and the coloured backgrounds represent different taxonomic groups, as shown in the key. Species names are as follows: Apla (*Acanthaster planci*), Aven (*Araneus ventricosus*), Bbel (*Branchiostoma belcheri*), Bgla (*Biomphalaria glabrata*), Csul (*Centruroides sculpturatus*), Drer (*Danio rerio)*, Ggal (*Gallus gallus*), Hsap (*Homo sapiens*), Lcha (*Latimeria chalumnae*), Lgig (*Lottia gigantea*), Mmus (*Mus musculus*), Obim (*Octopus bimaculoides*), Pcau (*Priapulus caudatus*), Pdum (*Platynereis dumerilii*), Pmar (*Petromyzon marinus*), Skow (*Saccoglossus kowalevskii*), Xtro (*Xenopus tropicalis*).

**Supplementary figure 6.**
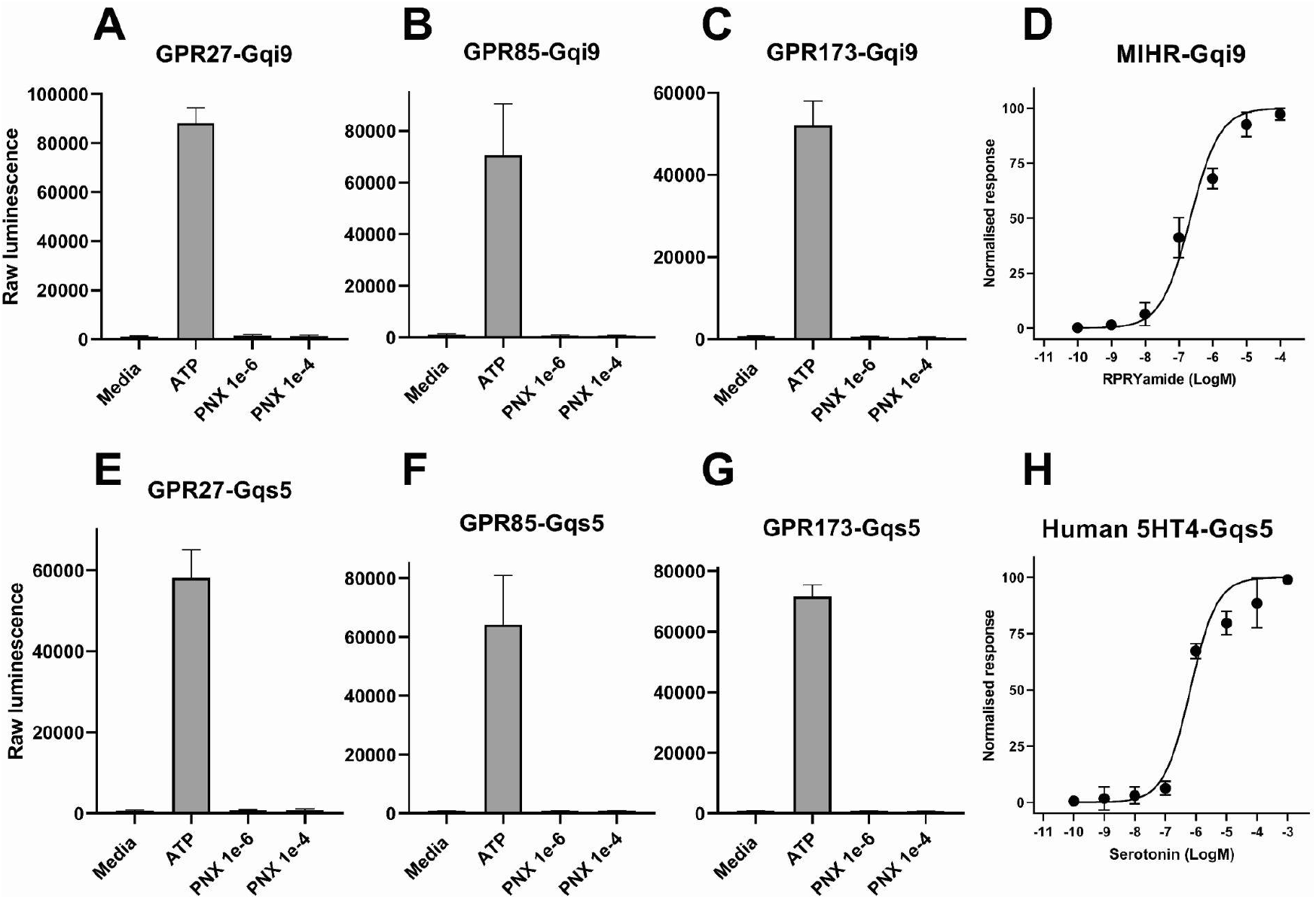
Receptor-ligand activation assay for human PNX-14 and human SREB receptors with different promiscuous G-proteins and positive controls. (A) GPR27 with Gqi9. (B) GPR85 with Gqi9. (C) GPR173 with Gqi9. (D) Positive control for Gqi9, the deorphanisation of the *Clytia hemisphaerica* MIH receptor is shown. (E) GPR27 with Gqs5. (F) GPR85 with Gqs5. (G) GPR173 with Gqi9. (H) Positive control for Gqs5 protein, the deorphanisation of human serotonin receptor 4 is shown.

**Supplementary file 1.** Source of the transcriptomes used for this analysis.

**Supplementary file 2.** Supermatrix used for the species tree in figure 3.

**Supplementary file 3.** Sequences of phoenixin precursors identified in different species with annotation.

**Supplementary file 4.** Data for the gene-structure of phoenixin precursors and accession number of the transcriptome and genome sequences used for the gene-structure analysis.

**Supplementary file 5.** Sequences of nesfatin-1 precursors identified in different species with annotation.

**Supplementary file 6.** Data for the gene-structure of NUCB precursors and accession number of the transcriptome and genome sequences used for the gene-structure analysis.

**Supplementary file 7.** Sequences used for the phylogenetic analysis of SREB receptors in supplementary figure 5.

**Supplementary file 8.** Aligned and trimmed sequences used for the reconstruction of SREB receptors in supplementary figure 5.

**Supplementary file 9.** Nexus file of the tree shown in supplementary figure 5

**Supplementary file 10.** Raw data for the receptor deorphanisation in supplementary figure 6.

**Supplementary file 11**. Detailed methodology of the research performed here

## Methods in brief

We obtained transcriptomes from public databases for 49 different taxa and assessed their completeness by a BUSCO analysis. To calculate a maximum-likelihood species tree, we used the aligned and trimmed BUSCO proteins as input for tree reconstruction with IQ-tree2 and the LG+G4 model. We searched the transcriptomes for Nesfatin and Phoenixin precursors by BLAST with bilaterian reference sequences as queries. Positive hits were tested for the presence of a signal peptide with SignalP-3.0. We analysed the exon-intron gene structure of the precursors with Splign. SREB GPCRs were retrieved by Hidden Markov Models and BLAST using vertebrate SREB reference sequences. We analysed the identified SREB sequences with CLANS and IQ-tree2. To test the activation of the human SREB receptors GPR27, GPR85 and GPR173 by the human Phoenixin-14 peptide, we transfected HEK cells with the receptors to carry out ligand-activation assays. Cells were stably expressing the calcium-sensitive bioluminescent reporter GFP-aequorin and were co-transfected with promiscuous Gq-proteins. A detailed description of the Material and Methods is given in Supplementary file 11.

## Detailed Materials and Methods

### Transcriptomic resources

Transcriptomes from different clades of metazoans, choanoflagellates, a filasterean and the flagellate *Tunicaraptor unikontum* were obtained from different public databases (see Supplementary file 1). We translated the transcripts into protein sequences with TransDecoder (TransDecoder; http://transdecoder.github.io/) with a minimum length of 50 amino acids. To assess the completeness of the transcriptomes, we ran BUSCO v5.2.1 (Manni et al. 2021) in protein mode and with the lineage set to ‘eukaryote’ with the database ‘eukaryota_odb10’ (Creation date of the database: Sep 2021, number of BUSCOs: 255).

### Phylogenomic analysis

To build a tree representing the relationships of the 49 species studied, we carried out a phylogenomic analysis with the output of the BUSCO analysis. BUSCO datasets comprise genes evolving under “single-copy control” (Waterhouse et al. 2011) and are near-universally present as single-copy orthologs across lineages. The eukaryotic database has 255 single-copy orthologs. We aligned these orthologs from each species individually with MAFFT v7 using the iterative refinement method L-INS-i (Katoh et al. 2002). The alignment was trimmed with the TrimAl software using the gappy-out method (Capella-Gutiérrez et al. 2009). Then, we concatenated the trimmed alignments with FASconcatG (Kück and Longo 2014), to assemble a concatenated supermatrix of 114,163 amino acid positions (Supplementary file 2). To build a species tree, we used IQ-TREE2 with the maximum-likelihood method under the LG+G4 model (Nguyen et al. 2015). The tree was rooted in the filasterean+*Tunicaraptor* clades. It is important to note that this phylogenomic analysis does not account for compositional bias and has been run with a homogeneous model (LG) only. The tree is merely used as a guide to map the evolutionary pattern of phoenixin, nesfatin-1 and the GPR173 across the species tree.

### Phoenixin and nesfatin-1 precursor identification and alignment

We identified the phoenixin precursor sequences by using the phoenixin precursor (SMIM20) from human and *Nephrops norvegicus* (Yosten et al. 2013; Nguyen et al. 2018) as queries. To search for nesfatin-1 precursor sequences, we used the human precursor as a query. We used a BLASTP with an e-value of e1-2 as the threshold to collect homologous sequences. In order to minimize the possibility of false positives, we manually curated the sequence list. To detect signal peptides, we used signalP-3.0. To align the full-length precursors and predicted mature peptides derived from them, we used MUSCLE (Edgar 2004). The lists of the sequences used for these alignments are available in Supplementary file 3 and Supplementary file 5.

### Gene-structure analyses of phoenixin and nesfatin-1 precursor sequences

In all the species in which we identified phoenixin and/or nefastin-1 precursors, we also searched for the corresponding genes with BLASTP in the GenBank database. For gene structure analysis, we selected at least one species from each of the major clades of metazoans and the choanoflagellate *S. rosetta* and we retrieved the transcripts and genomic regions. We used the tool Splign (Kapustin et al., 2008) to determine the exon/intron structure of the genes (https://www.ncbi.nlm.nih.gov/sutils/splign/splign.cgi).Based on these data, the gene-structure diagrams were drawn in Adobe Illustrator CS6. The output of the Splign analysis is available in supplementary files 4 and 6 (for phoenixin and nesfatin-1, respectively).

### GPR173 identification and phylogenetic analysis

To identify GPR173 receptors, we obtained a database of vertebrate SREB sequences, including GPR173, GPR85 and GPR27 from (Breton et al. 2021). From these sequences, we produced a Hidden Markov Model (HMM)-model and used this model to mine the 49 species investigated. HMM models were run in HMMR3 with an e-value of 1e-15. The same SREB sequences were used to carry out similarity-based searches using BLASTP with an e-value cutoff of 1e-15. We merged these two databases and ran CD-Hit to eliminate redundant sequences (at a 99% threshold). To identify the sequences that are closely related to the GPR173 sequences, we ran a cluster-based analysis in CLANS. To identify clusters, we used the convex-clustering option with 100 jackknife replicates. The SREB receptors are extremely well conserved and form an easily recognisable cluster. The CLANS analysis is available as Supplementary figure 4.

To analyse the phylogeny of SREB receptors, the cluster containing these receptors together with monoaminergic receptors were parsed and used for tree building. We aligned the sequences with MAFFT version 7, with the iterative refinement method E-INS-i. Alignments were trimmed with TrimAl in gappy-out mode (Capella-Gutiérrez et al. 2009). To calculate maximum-likelihood trees, we used IQ-tree2 with the LG+G4 model. To calculate branch support, we ran 1,000 replicates with the aLRT-SH-like and aBayes methods (Minh et al. 2020). The sequences used for the phylogenetic analysis are available in Supplementary file 7, the trimmed alignment is available in Supplementary file 8. The raw nexus tree of the SREB receptors is available in Supplementary file 9.

### GPR173 deorphanisation assays

We ordered synthetic mature peptides PNX14 from GenScript with a purity of >95%. The receptor GPR173 was purchased from the GenScript GenEZ human ORFs database (Accession No. NM_018969.6) and cloned into a pCDNA3.1(+) vector with EcoRV and a blunt cloning strategy. We expressed the receptor in HEK293 cells that were stably expressing the calcium-sensitive bioluminescent reporter GFP-aequorin fusion protein (G5A). This cell line was purchased from Angio-proteomie (CAT no. cAP-0200GFP-AEQ-Cyto). The HEK293-G5a cells were maintained at 37 °C in 96 well-plates containing 100 μl of DMEM, high glucose glutamax medium (Thermo; Cat. No. 10566016) supplemented with 10% fetal bovine serum (Thermo; Cat. No. 10082147). Upon reaching confluency of approximately 85%, we transfected the cells with the plasmid containing GPR173 and a plasmid containing the promiscuous Gαqi9 [Addgene; Cat. No. 125711 (Masharina et al. 2012)] or Gαqs5 [Addgene; Cat. No. 24498] (Conklin et al. 1996).

Transfections were carried out with 60 ng of each plasmid and 0.35 μl of the transfection reagent Transfectamine 5000 (AAT-bioquest; Cat. No. 60022). Two days post-transfection, we removed the culture medium and substituted it for fresh DMEM-medium supplemented with 4 mM coelenterazine-H (Thermo Fisher Scientific; Cat. No. C6780). After an incubation period of 3 hr, we exposed the cells to synthetic PNX-14 peptide diluted in DMEM-medium in concentrations ranging from 10-4 M to 10-6 M. Luminescence levels were recorded over a 60-second period in a FlexStation 3 Multi-Mode Microplate Reader (Molecular Devices). We integrated the luminescence data over a 60-second measurement period. A minimum of two independent transfections with triplicate measurements were made for each concentration, and the average of each was used to normalise the responses. We normalised the responses to the maximum response obtained by the addition of 100 μM ATP in each experiment (100% activation) and to the response obtained with the vehicle media (0% activation). As positive control for the Gαqi/9 protein, we used the *Clytia hemisphaerica* MIH receptor and its MIH peptide RPRYamide (Quiroga Artigas et al. 2020). As positive control for the Gαqs5 protein, we used the human serotonin receptor 4 (5-HTR4) purchased from the GenScript GenEZ human ORFs database (Accession No. NM_000870.6) and deorphanised with serotonin hydrochloride purchased from Sigma-Aldrich (Cat No.H9523). For the positive controls, dose-response curves were fitted with a four-parameter curve based on the normalized data from the average of three independent transfections using Prism 8 (GraphPad, La Jolla, USA). The raw data obtained from the deorphanisation assays shown in Supplementary Figure 6 are available in Supplementary file 10.

## Bibliography

Ayada C, Toru Ü, Korkut Y. 2015. Nesfatin-1 and its effects on different systems. Hippokratia 19:4–10.

Bendtsen JD, Nielsen H, von Heijne G, Brunak S. 2004. Improved prediction of signal peptides: SignalP 3.0. J. Mol. Biol. 340:783–795.

Bersani M, Johnsen AH, Højrup P, Dunning BE, Andreasen JJ, Holst JJ. 1991. Human galanin: primary structure and identification of two molecular forms. FEBS Lett. 283:189–194.

Billert M, Rak A, Nowak KW, Skrzypski M. 2020. Phoenixin: More than Reproductive Peptide. Int. J. Mol. Sci. [Internet] 21. Available from: http://dx.doi.org/10.3390/ijms21218378

Breton TS, Sampson WGB, Clifford B, Phaneuf AM, Smidt I, True T, Wilcox AR, Lipscomb T, Murray C, DiMaggio MA. 2021. Characterization of the G protein-coupled receptor family SREB across fish evolution. Scientific Reports [Internet] 11. Available from: http://dx.doi.org/10.1038/s41598-021-91590-9

Burkhardt P, Grønborg M, McDonald K, Sulur T, Wang Q, King N. 2014. Evolutionary insights into premetazoan functions of the neuronal protein homer. Mol. Biol. Evol. 31:2342–2355.

Burkhardt P, Jékely G. 2021. Evolution of Synapses and Neurotransmitter Systems: The Divide-and-Conquer Model for Early Neural Cell-Type Evolution. Preprints [Internet]. Available from: http://dx.doi.org/10.20944/preprints202110.0019.v1

Capella-Gutiérrez S, Silla-Martínez JM, Gabaldón T. 2009. trimAl: a tool for automated alignment trimming in large-scale phylogenetic analyses. Bioinformatics 25:1972–1973.

Clarke SA, Dhillo WS. 2019. Phoenixin and Its Role in Reproductive Hormone Release. Seminars in Reproductive Medicine [Internet] 37:191–196. Available from: http://dx.doi.org/10.1055/s-0039-3400964

Conklin BR, Herzmark P, Ishida S, Voyno-Yasenetskaya TA, Sun Y, Farfel Z, Bourne HR. 1996. Carboxyl-terminal mutations of Gq alpha and Gs alpha that alter the fidelity of receptor activation. Mol. Pharmacol. 50:885–890.

Edgar RC. 2004. MUSCLE: multiple sequence alignment with high accuracy and high throughput. Nucleic Acids Res. 32:1792–1797.

Elphick MR, Mirabeau O, Larhammar D. 2018. Evolution of neuropeptide signalling systems. J. Exp. Biol. [Internet] 221. Available from: http://dx.doi.org/10.1242/jeb.151092

Foster SR, Hauser AS, Vedel L, Strachan RT, Huang X-P, Gavin AC, Shah SD, Nayak AP, Haugaard-Kedström LM, Penn RB, et al. 2019. Discovery of Human Signaling Systems: Pairing Peptides to G Protein-Coupled Receptors. Cell [Internet] 179:895–908.e21. Available from: http://dx.doi.org/10.1016/j.cell.2019.10.010

Friedrich T, Schalla MA, Scharner S, Kühne SG, Goebel-Stengel M, Kobelt P, Rose M, Stengel A. 2019. Intracerebroventricular injection of phoenixin alters feeding behavior and activates nesfatin-1 immunoreactive neurons in rats. Brain Res. 1715:188–195.

Friedrich T, Stengel A. 2021. Role of the Novel Peptide Phoenixin in Stress Response and Possible Interactions with Nesfatin-1. Int. J. Mol. Sci. [Internet] 22. Available from: http://dx.doi.org/10.3390/ijms22179156

Göhde R, Naumann B, Laundon D, Imig C, McDonald K, Cooper BH, Varoqueaux F, Fasshauer D, Burkhardt P. 2021. Choanoflagellates and the ancestry of neurosecretory vesicles. Philos. Trans. R. Soc. Lond. B Biol. Sci. 376:20190759.

Haddock CJ, Almeida-Pereira G, Stein LM, Yosten GLC, Samson WK. 2020. A novel regulator of thirst behavior: phoenixin. Am. J. Physiol. Regul. Integr. Comp. Physiol. 318:R1027–R1035.

Hauser AS, Gloriam DE, Bräuner-Osborne H, Foster SR. 2020. Novel approaches leading towards peptide GPCR de-orphanisation. Br. J. Pharmacol. 177:961–968.

Hofmann T, Ahnis A, Elbelt U, Rose M, Klapp BF, Stengel A. 2015. NUCB2/nesfatin-1 Is Associated with Elevated Levels of Anxiety in Anorexia Nervosa. PLoS One 10:e0132058.

Hofmann T, Weibert E, Ahnis A, Elbelt U, Rose M, Klapp BF, Stengel A. 2017. Phoenixin is negatively associated with anxiety in obese men. Peptides 88:32–36.

Hook V, Funkelstein L, Lu D, Bark S, Wegrzyn J, Hwang S-R. 2008. Proteases for processing proneuropeptides into peptide neurotransmitters and hormones. Annu. Rev. Pharmacol. Toxicol. 48:393–423.

Jékely G. 2013. Global view of the evolution and diversity of metazoan neuropeptide signaling. Proc. Natl. Acad. Sci. U. S. A. 110:8702–8707.

Jékely G. The chemical brain hypothesis for the origin of nervous systems. Available from: http://dx.doi.org/10.31234/osf.io/3tv7p

Jiang JH, He Z, Peng YL, Jin WD, Mu J, Xue HX, Wang Z, Chang M, Wang R. 2015. Effects of Phoenixin-14 on anxiolytic-like behavior in mice. Behav. Brain Res. 286:39–48.

Kmiecik AM, Dzięgiel P, Podhorska-Okołów M. 2021. Nucleobindin-2/Nesfatin-1—A New Cancer Related Molecule? International Journal of Molecular Sciences [Internet] 22:8313. Available from: http://dx.doi.org/10.3390/ijms22158313

Koch TL, Grimmelikhuijzen CJP. 2020. A comparative genomics study of neuropeptide genes in the cnidarian subclasses Hexacorallia and Ceriantharia. BMC Genomics 21:666.

Kück P, Longo GC. 2014. FASconCAT-G: extensive functions for multiple sequence alignment preparations concerning phylogenetic studies. Front. Zool. 11:81.

Kumar D, Blaby-Haas CE, Merchant SS, Mains RE, King SM, Eipper BA. 2016. Early eukaryotic origins for cilia-associated bioactive peptide amidating activity. Journal of Cell Science [Internet]. Available from: http://dx.doi.org/10.1242/jcs.177410

Kumar D, Strenkert D, Patel-King RS, Leonard MT, Merchant SS, Mains RE, King SM, Eipper BA. 2017. A bioactive peptide amidating enzyme is required for ciliogenesis. eLife [Internet] 6. Available from: http://dx.doi.org/10.7554/elife.25728

Luxmi R, Blaby-Haas C, Kumar D, Rauniyar N, King SM, Mains RE, Eipper BA. 2018. Proteases Shape the Chlamydomonas Secretome: Comparison to Classical Neuropeptide Processing Machinery. Proteomes [Internet] 6. Available from: https://www.ncbi.nlm.nih.gov/pmc/articles/PMC6313938/

Luxmi R, Kumar D, Mains RE, King SM, Eipper BA. 2019. Cilia-based peptidergic signaling. PLOS Biology [Internet] 17:e3000566. Available from: http://dx.doi.org/10.1371/journal.pbio.3000566

Lyu R-M, Huang X-F, Zhang Y, Dun SL, Luo JJ, Chang J-K, Dun NJ. 2013. Phoenixin: a novel peptide in rodent sensory ganglia. Neuroscience 250:622–631.

Maejima Y, Sedbazar U, Suyama S, Kohno D, Onaka T, Takano E, Yoshida N, Koike M, Uchiyama Y, Fujiwara K, et al. 2009. Nesfatin-1-regulated oxytocinergic signaling in the paraventricular nucleus causes anorexia through a leptin-independent melanocortin pathway. Cell Metab. 10:355–365.

Ma H, Su D, Wang Q, Chong Z, Zhu Q, He W, Wang W. 2020. Phoenixin 14 inhibits ischemia/reperfusion-induced cytotoxicity in microglia. Arch. Biochem. Biophys. 689:108411.

Manni M, Berkeley MR, Seppey M, Simão FA, Zdobnov EM. 2021. BUSCO Update: Novel and Streamlined Workflows along with Broader and Deeper Phylogenetic Coverage for Scoring of Eukaryotic, Prokaryotic, and Viral Genomes. Mol. Biol. Evol. 38:4647–4654.

Masharina A, Reymond L, Maurel D, Umezawa K, Johnsson K. 2012. A fluorescent sensor for GABA and synthetic GABA(B) receptor ligands. J. Am. Chem. Soc. 134:19026–19034.

Matsumoto M, Saito T, Takasaki J, Kamohara M, Sugimoto T, Kobayashi M, Tadokoro M, Matsumoto S, Ohishi T, Furuichi K. 2000. An evolutionarily conserved G-protein coupled receptor family, SREB, expressed in the central nervous system. Biochem. Biophys. Res. Commun. 272:576–582.

Merali Z, Cayer C, Kent P, Anisman H. 2008. Nesfatin-1 increases anxiety- and fear-related behaviors in the rat. Psychopharmacology [Internet] 201:115–123. Available from: http://dx.doi.org/10.1007/s00213-008-1252-2

Minh BQ, Schmidt HA, Chernomor O, Schrempf D, Woodhams MD, von Haeseler A, Lanfear R. 2020. IQ-TREE 2: New Models and Efficient Methods for Phylogenetic Inference in the Genomic Era. Mol. Biol. Evol. 37:1530–1534.

Mirabeau O, Joly J-S. 2013. Molecular evolution of peptidergic signaling systems in bilaterians. Proc. Natl. Acad. Sci. U. S. A. 110:E2028–E2037.

Miura K, Kurosawa Y, Kanai Y. 1994. Calcium-binding activity of nucleobindin mediated by an EF hand moiety. Biochem. Biophys. Res. Commun. 199:1388–1393.

Miura K, Titani K, Kurosawa Y, Kanai Y. 1992. Molecular cloning of nucleobindin, a novel DNA-binding protein that contains both a signal peptide and a leucine zipper structure. Biochem. Biophys. Res. Commun. 187:375–380.

Moran Y, Zakon HH. 2014. The evolution of the four subunits of voltage-gated calcium channels: ancient roots, increasing complexity, and multiple losses. Genome Biol. Evol. 6:2210–2217.

Nguyen TV, Rotllant GE, Cummins SF, Elizur A, Ventura T. 2018. Insights Into Sexual Maturation and Reproduction in the Norway Lobster () via Prediction and Characterization of Neuropeptides and G Protein-coupled Receptors. Front. Endocrinol. 9:430.

Oh-I S, Shimizu H, Satoh T, Okada S, Adachi S, Inoue K, Eguchi H, Yamamoto M, Imaki T, Hashimoto K, et al. 2006. Identification of nesfatin-1 as a satiety molecule in the hypothalamus. Nature 443:709–712.

de Oliveira AL, Calcino A, Wanninger A. 2019. Ancient origins of arthropod moulting pathway components. Elife [Internet] 8. Available from: http://dx.doi.org/10.7554/eLife.46113

Otte S, Barnikol-Watanabe S, Vorbrüggen G, Hilschmann N. 1999. NUCB1, the Drosophila melanogaster homolog of the mammalian EF-hand proteins NEFA and nucleobindin. Mech. Dev. 86:155–158.

Pałasz A, Rojczyk E, Bogus K, Worthington JJ, Wiaderkiewicz R. 2015. The novel neuropeptide phoenixin is highly co-expressed with nesfatin-1 in the rat hypothalamus, an immunohistochemical study. Neurosci. Lett. 592:17–21.

Quiroga Artigas G, Lapébie P, Leclère L, Bauknecht P, Uveira J, Chevalier S, Jékely G, Momose T, Houliston E. 2020. A G protein-coupled receptor mediates neuropeptide-induced oocyte maturation in the jellyfish Clytia. PLoS Biol. 18:e3000614.

Roch GJ, Sherwood NM. 2014. Glycoprotein Hormones and Their Receptors Emerged at the Origin of Metazoans. Genome Biology and Evolution [Internet] 6:1466–1479. Available from: http://dx.doi.org/10.1093/gbe/evu118

Rupp SK, Wölk E, Stengel A. 2021. Nesfatin-1 Receptor: Distribution, Signaling and Increasing Evidence for a G Protein-Coupled Receptor - A Systematic Review. Front. Endocrinol. 12:740174.

Sachkova MY, Nordmann E-L, Soto-Àngel JJ, Meeda Y, Górski B, Naumann B, Dondorp D, Chatzigeorgiou M, Kittelmann M, Burkhardt P. 2021. Neuropeptide repertoire and 3D anatomy of the ctenophore nervous system. Curr. Biol. [Internet]. Available from: http://dx.doi.org/10.1016/j.cub.2021.09.005

Schalla MA, Goebel-Stengel M, Friedrich T, Kühne SG, Kobelt P, Rose M, Stengel A. 2020. Restraint stress affects circulating NUCB2/nesfatin-1 and phoenixin levels in male rats. Psychoneuroendocrinology 122:104906.

Schalla MA, Stengel A. 2018. Current Understanding of the Role of Nesfatin-1. J Endocr Soc 2:1188–1206.

Schalla MA, Stengel A. 2019. The role of phoenixin in behavior and food intake. Peptides [Internet] 114:38–43. Available from: http://dx.doi.org/10.1016/j.peptides.2019.04.002

Schalla M, Prinz P, Friedrich T, Scharner S, Kobelt P, Goebel-Stengel M, Rose M, Stengel A. 2017. Phoenixin-14 injected intracerebroventricularly but not intraperitoneally stimulates food intake in rats. Peptides [Internet] 96:53–60. Available from: http://dx.doi.org/10.1016/j.peptides.2017.08.004

Senatore A, Reese TS, Smith CL. 2017. Neuropeptidergic integration of behavior in, an animal without synapses. J. Exp. Biol. 220:3381–3390.

Sobrido-Cameán D, Yáñez-Guerra LA, Lamanna F, Conde-Fernández C, Kaessmann H, Elphick MR, Anadón R, Rodicio MC, Barreiro-Iglesias A. 2019. Galanin in an Agnathan: Precursor Identification and Localisation of Expression in the Brain of the Sea Lamprey Petromyzon marinus. Frontiers in Neuroanatomy [Internet] 13. Available from: http://dx.doi.org/10.3389/fnana.2019.00083

Stein LM, Tullock CW, Mathews SK, Garcia-Galiano D, Elias CF, Samson WK, Yosten GLC. 2016. Hypothalamic action of phoenixin to control reproductive hormone secretion in females: importance of the orphan G protein-coupled receptor Gpr173. Am. J. Physiol. Regul. Integr. Comp. Physiol. 311:R489–R496.

Stengel A, Goebel-Stengel M, Wang L, Kato I, Mori M, Taché Y. 2012. Nesfatin-130–59 but not the N- and C-terminal fragments, nesfatin-11–29 and nesfatin-160–82 injected intracerebroventricularly decreases dark phase food intake by increasing inter-meal intervals in mice. Peptides [Internet] 35:143–148. Available from: http://dx.doi.org/10.1016/j.peptides.2012.03.015

Sundarrajan L, Blanco AM, Bertucci JI, Ramesh N, Canosa LF, Unniappan S. 2016. Nesfatin-1-Like Peptide Encoded in Nucleobindin-1 in Goldfish is a Novel Anorexigen Modulated by Sex Steroids, Macronutrients and Daily Rhythm. Sci. Rep. 6:28377.

Thiel D, Yañez-Guerra LA, Franz-Wachtel M, Hejnol A, Jékely G. 2021. Nemertean, Brachiopod, and Phoronid Neuropeptidomics Reveals Ancestral Spiralian Signaling Systems. Mol. Biol. Evol. 38:4847–4866.

Tikhonenkov, et al. 2020. New Lineage of Microbial Predators Adds Complexity to Reconstructing the Evolutionary Origin of Animals. Curr. Biol. 30:4500–4509.e5.

Treen AK, Luo V, Belsham DD. 2016. Phoenixin Activates Immortalized GnRH and Kisspeptin Neurons Through the Novel Receptor GPR173. Mol. Endocrinol. 30:872–888.

Varoqueaux F, Williams EA, Grandemange S, Truscello L, Kamm K, Schierwater B, Jékely G, Fasshauer D. 2018. High Cell Diversity and Complex Peptidergic Signaling Underlie Placozoan Behavior. Curr. Biol. 28:3495–3501.e2.

Veenstra JA. 2000. Mono- and dibasic proteolytic cleavage sites in insect neuroendocrine peptide precursors. Arch. Insect Biochem. Physiol. 43:49–63.

Yañez-Guerra LA, Zhong X, Moghul I, Butts T, Zampronio CG, Jones AM, Mirabeau O, Elphick MR. 2020. Echinoderms provide missing link in the evolution of PrRP/sNPF-type neuropeptide signalling. eLife [Internet] 9. Available from: http://dx.doi.org/10.7554/elife.57640

Yao B, Lv J, Du L, Zhang H, Xu Z. 2021. Phoenixin-14 protects cardiac damages in a streptozotocin-induced diabetes mice model through SIRT3. Arch. Physiol. Biochem.:1–9.

Yosten GLC, Kolar GR, Salvemini D, Samson WK. 2021. The Deductive Reasoning Strategy Enables Biomedical Breakthroughs. Mo. Med. 118:352–357.

Yosten GLC, Lyu R-M, Hsueh AJW, Avsian-Kretchmer O, Chang J-K, Tullock CW, Dun SL, Dun N, Samson WK. 2013. A novel reproductive peptide, phoenixin. J. Neuroendocrinol. 25:206–215.

Zandawala M, Moghul I, Yañez Guerra LA, Delroisse J, Abylkassimova N, Hugall AF, O’Hara TD, Elphick MR. 2017. Discovery of novel representatives of bilaterian neuropeptide families and reconstruction of neuropeptide precursor evolution in ophiuroid echinoderms. Open Biol. [Internet] 7. Available from: http://dx.doi.org/10.1098/rsob.170129

