## Supplementary file 3 for "Pre-metazoan origin of neuropeptide signalling"

Potential cleavage site

Potential cleavage and amidation sites

Signal peptide

>Homo sapiens _Q8N5G0_SIM20_Small_integral_membrane_protein_20

MSRNLRTALIFGGFISLIGAAFYPIYFRPLMRLEEYKKEQAINRAGIVQEDVQPPGLKVWSDPFGRK

>Mus_musculus_D3Z7Q2

MAAARNLRTALIFGGFISMVGAAFYPIYFRPLMRLEEYQKEQAVNRAGIVQEDVQPPGLKVWSDPFGRK

>Gallus_gallus_F1NQM1

MARLFRTLVIFGGFAAVVGAAFYPIYFRPLLLPEEYKREQSINRAGIVQENIQPPGLKVWSDPFGRK

>Xenopus_tropicalis_A0A803K4P1

MSKNTRIVLIFGGFVTAVAAAFYPIFFHPLMHIDEYKKEQAVNRTDVIQENVQPTGLKVWSDPFSRK

>Danio_rerio_A0A2R8Q9H3

MSSSKRITLIFGGFVAAVAAAFYPIFFHPLTHSEDYKQVQKVNRAGVNQADIQPVGVKVWSDPYKPKS

>Deutero_Cephalo_118270R.t1

MSKDIPFLATMSGTSMRRMVLGIGAFAAVVGGVMYPIFIAPLLHSQEYQKMQKQNREGIKQDEIQPGGMVVWSNPFGEKKDQKKDS*

>Mnemiopsis_leidyi_GFAT01018176_1_p1

MSYRMSGWKYGAFIGGIVGFIGLALYPTVIYPMQHIDEYKEIQKSNRAGIIQENIQPGGMKVWSDPFGRKKSE

>Araneus_ventricosus_A0A4Y2BV56

MGRPVKLKGLRAFAFYGGLGGLVGACVYAVVIYPYLHIDEYKEIQKITRAGIDQEKIQPGGMKVWSDPFDKRK

>Cteno_Puk_Fal_comp9277_c0_seq2

MYNVNKPVSRGLRRDKNSKKSFTVMVLLKGWRYFFFIGSLVGFTGAAIYPVIIQPLMDPEPYKKIQAETRKGIRQEDIQPGNMKIWTDPFDRKK*

>Daphnia_pulex_E9FSU6

MVLLKGWKYAAMVGTVVGAIGLAIYPIIVSPMMNPEPYKKQQAVNRSGIKQEEVQPGNMKVWSDPFARPKAEEK

>Tribolium_castaneum_D6WAU5

MTILRGWRYNVFIGGLVGVIALTLYPIAIEPMLNSEKYKKIQERNRRGIKQEDVQPGNMKVWSDPFGRK

>Phoronis_ijimai_comp96582_c0_seq1

MGFKASEGHLMTGWRFGLFFGGLIGLIGLAVYPVIIEPYLNQDKWKKIQQSARQGIKQDEVQPGNMKVWSDPFEKKN*

>Cnidar_AcrMil_XR_003823459.1

MSHKVKTGVFIAGFTALTLAALYPIVVYPKLHPEVYKDIQMETRKSIKQEDIQPGNMKVWSDPFSRK*

>Cnidar_NemVec_comp12848_c0_seq2

MSSQVRSGIVIGGFVALTVAMMFPIAIYPKLFPEVYQDIQRDRRKGIKQEEIQPGNMKVWSDPFKKD*

>Amphimedon_queenslandica_qu2.1.41283_001

MGVVKWEGCSTCFMPSNRTLKNAGLAAGFLTLLGAALYSVIIYPKLHPEIYREKQKWIRDKVPLEKTQPGGMKVWTDPFGRKDSKK

>Corallium_rubrum_TR16776_c0_g1_i1.p4

MANSKGIVVIGALVSAIGLALYPIAVHPKLFPQKYKNIQKENRKEIVQEDIQPGGMKVWTDPFKKN*

>Saccoglossus_kowalevskii_v30017359m

MPVRNWWVAAGMVTLISAALLPVVVQPWLNPKTWQDVQKESRKGIKQEEIQPGGMRVWSDPFEPRK*

>Clytia_hemisphaerica_TCONS_00046557.p1

MNRNVIFVGGLVAAIGIALVPIYFVPMKRPDIYHNIQKDTRKDIVQEEVQPGNMKVWSDPFDRKKE*

>Hydra_vulg_Sc4wPfr_945.g15400.t1

MSLISKLAVYGLVLATGLSLVPIYFVPKAVPEKYRDIQKVSRKDIVQSEVQPGNMKIWSDPFDRKK

>Hydractina_echinatA_TRINITY_DN51421_c0_g1_i1.p1

MVKGKWVGVAFVVAITGALLPVYFMPKINPDMYRHIQKETRKDIVQNEVQPGGMKIWSDPFDRKK*

>XR_001216320.1 protein Biomphalaria glabrata

MNSIRENFRSIMVIATFVGGIVAASYPIIISPYLNPAPWKKEQSIGRRGIDREAIQPGGMKVWTDPFDRK

>Cnidar_Rhopilema_esculentum_TR100627_c0_g1_i1.p2

MAKRDLFLFGGIAAAIGLALLPIYAYPKFNPNGYNNVQERNREGLDREAIQPGGMKIWNDPFDRKKNSSGEE*

>Cnidar_AlaAla_agalma_gene_44948.1_

MGRREKVFIGSICGAILVTHLFVYVIPKMNKNKYKDIQSETRSNIAREDIQPGEMKIWSDPFEREKK*

>tr_F2UJQ3_F2UJQ3_SALR5_Uncharacterized_protein_OS_Salpingoeca_rosetta__strain

MARGGKSAAAVVGVVALTMAALYPIVFAPLMDASEYRGRQQEFLREQGLSKKDIQPEGLPVWSDPFDQKKSK

>Porifera_OscCar_comp20502_c0_seq1.p1

MKKNVAVIGGLVGLIGISLYAVVVHPRLHPEKWRKMQEPVRNKTTLEETQPGGMKVWEDPFGRKKKSD

>Protostome_ecdy_Pdum_comp424107_c0_seq4

MANMTGWRFALFIGSMTGAVGAVMYSVAIDPYLHPEKWQDVQRETRRGIDQASVQPAGLKVWSDPFDRPKKDS

>XR_004405356.1_Petromyzon marinus uncharacterized LOC116958355

MGRITRTTLVFGGFVAALAVTIYPIAIHPFLNIADFQVVQKVNREGVDQESIQPGGMKVWSNPFEPKK
