## Supplementary file 4 for "Pre-metazoan origin of neuropeptide signalling"

Homo sapiens

>NP_001138904.1 small integral membrane protein 20 [Homo sapiens]

MSRNLRTALIFGGFISLIGAAFYPIYFRPLMRLEEYKKEQAINRAGIVQEDVQPPGLKVWSDPFGRK

>NP_001138904.1 small integral membrane protein 20 [Homo sapiens]

MSRNLRTALIFGGFISLIGAAFYPIYFRPLMRLEEY--(1)-[13891bp]--

KKEQAINRAGIVQEDVQPP--(1)-[785bp]--

GLKVWSDPFGRK

>NM_001145432.3 Homo sapiens small integral membrane protein 20 (SMIM20), transcript variant 1

ATGTCCCGGAACCTGCGCACCGCGCTCATTTTCGGCGGCTTCATCTCCCTGATCGGCGCCGCCTTCTATCCCATCTACTTCCGGCCCCTAATGAGATTGGAGGAGTACAAGAAGGAACAAGCTATAAATCGGGCTGGAATTGTTCAAGAGGATGTGCAGCCACCAGGGTTAAAAGTGTGGTCTGATCCATTTGGCAGGAAATGA

Genome: splign

Gallus gallus

>NP_001138902.1 [Gallus gallus]

MARLFRTLVIFGGFAAVVGAAFYPIYFRPLLLPEEYKREQSINRAGIVQENIQPPGLKVWSDPFGRK

>NP_001138902.1 [Gallus gallus]

MARLFRTLVIFGGFAAVVGAAFYPIYFRPLLLPEEY--(1)-[1916bp]--

KREQSINRAGIVQENIQPP--(1)-[1228bp]--

GLKVWSDPFGRK

Transcript: NM_001145430.1

Genome: splign

Branchiostoma belcheri

>XP_019625233.1 [Branchiostoma belcheri]

MSGTSMRRMVLGIGAFAAVVGGVMYPIFIAPLLHSQEYQKMQKQNREGIKQDEIQPGGMVVWSNPFGEKKDQKKDS

>XP_019625233.1 [Branchiostoma belcheri]

MSGTSMRRMVLGIGAFAAVVGGVMYPIFIAPLLHSQEY--(1)-[398bp]--

QKMQKQNREGIKQDEIQPG--(1)-[337bp]--

GMVVWSNPFGEKKDQKKDS

Transcript: XM_019769674.1

Genome: NW_017803824.1

Tribolium castaneum

>XP_001808390.2 [Tribolium castaneum]

MTILRGWRYNVFIGGLVGVIALTLYPIAIEPMLNSEKYKKIQERNRRGIKQEDVQPGNMKVWSDPFGRK

>XP_001808390.2 [Tribolium castaneum]

MTILRGWRYNVFIGGLVGVIALTLYPIAIEPMLNSEKY--(1)-[48bp]--

KKIQERNRRGIKQEDVQPG--(1)-[52bp]--

NMKVWSDPFGRK

Transcript: XM_001808338.3

Genome: splign

Biomphalaria glabrata

>XR_001216320.1 [Biomphalaria glabrata]

MNSIRENFRSIMVIATFVGGIVAASYPIIISPYLNPAPWKKEQSIGRRGIDREAIQPGGMKVWTDPFDRK

>XR_001216320.1 [Biomphalaria glabrata]

MNSIRENFRSIMVIATFVGGIVAASYPIIISPYLNPAPW--(1)-[1759bp]--

KKEQSIGRRGIDREAIQPG--(1)-[1603bp]--

GMKVWTDPFDRK

Transcript: XR_001216320.1

Genome: JABXJC010000399.1

Nematostella vectensis

>comp12848_c0_seq2_.p1 [Nematostella vectensis]

MSSQVRSGIVIGGFVALTVAMMFPIAIYPKLFPEVYQDIQRDRRKGIKQEEIQPGNMKVWSDPFKKD

>comp12848_c0_seq2_.p1 [Nematostella vectensis]

MSSQVRSGIVIGGFVALTVAMMFPIAIYPKLFPEVY--(1)-[158bp]--

QDIQRDRRKGIKQEEIQPG--(1)-[1064bp]--

NMKVWSDPFKKD

>HADP01171693.1 Nematostella vectensis, contig TR67875

ATGTCCAGTCAAGTGAGATCTGGTATTGTTATCGGGGGTTTTGTGGCGCTAACGGTGGCTATGATGTTTCCAATCGCCATCTACCCAAAGCTGTTCCCCGAAGTCTACCAGGATATACAAAGAGATAGGAGAAAGGGAATAAAACAAGAGGAAATTCAGCCTGGAAACATGAAAGTATGGTCAGATCCATTTAAGAAGGACTAG

Genome: <https://simrbase.stowers.org/starletseaanemone>, chromosome 10

(Zimmermann et al 2020, bioRxiv: https://www.biorxiv.org/content/10.1101/2020.10.30.359448v1)

Hydra vulgaris

>XR_001140944.1 (translated) [Hydra vulgaris]

MSLISKLAVYGLVLATGLSLVPIYFVPKAVPEKYRDIQKVSRKDIVQSEVQPGNMKIWSDPFDRKK

>XR_001140944.1 (translated) [Hydra vulgaris]

MSLISKLAVYGLVLATGLSLVPIYFVPKAVPEKY--(1)-[10041bp]--

RDIQKVSRKDIVQSEVQPG--(1)-[1642bp]--

NMKIWSDPFDRKK

Transcript: XR_001140944.1

Genome: NW_004166935.1

Amphimedon queenslandica

>Aqu2.1.41283_001.p1 [Amphimedon queenslandica]

MGVVKWEGCSTCFMPSNRTLKNAGLAAGFLTLLGAALYSVIIYPKLHPEIYREKQKWIRDKVPLEKTQPGGMKVWTDPFGRKDSKK

>Aqu2.1.41283_001.p1 [Amphimedon queenslandica]

MGVVKWE--(2)-[76bp]--

GCSTCFMPSNRTLKNAGLAAGFLTLLGAALYSVIIYPKLHPEIY--(1)-[183bp]--

REKQKWIRDKVPLEKTQPG--(1)-[50bp]--

GMKVWTDPFGRKDSKK

>Aqu2.1.41283_001 cdna

ATGGGCGTGGTTAAGTGGGAGGGCTGCTCAACCTGCTTCATGCCTAGCAATAGGACATTAAAGAATGCTGGCCTTGCTGCTGGCTTCCTAACTCTCCTTGGAGCGGCTCTTTACTCAGTCATCATTTATCCAAAGCTCCACCCTGAAATCTATAGAGAAAAGCAAAAGTGGATACGTGATAAAGTTCCATTGGAGAAAACTCAACCAGGAGGTATGAAGGTCTGGACTGATCCATTTGGAAGAAAGGACTCAAAAAAATAA

Genome: ACUQ01000725.1

Salpingoeca rosetta

>XP_004990696.1 [Salpingoeca rosetta]

MARGGKSAAAVVGVVALTMAALYPIVFAPLMDASEYRGRQQEFLREQGLSKKDIQPEGLPVWSDPFDQKK

SK

>XP_004990696.1 [Salpingoeca rosetta]

MARGGKSAAAVVGVVALTMAALYPIVFAPLMDASEY--(1)-[227bp]--

RGRQQEFLREQGLSKKDIQPE--(1)-[213bp]--

GLPVWSDPFDQKK

SK

Transcript: XM_004990639.1

Genome: ACSY01002029.1
