## Supplementary file 5 for "Pre-metazoan origin of neuropeptide signalling"

Potential cleavage site

Potential cleavage and amidation sites

>Cnidaria_H.echinata_DN32624_c2_g1_i1.p1

MDTQIKLILLSFLIGQLVAPPPVERNQAEMTDNWKYNTEGLTKEQAQDQEYFRYLTQVVGELEKDADFKSVLNNATEDDIRSGKIAEHFDLVGHHIRQKLDEVKRLEVEYQRDLLRQKKEHMLGIDRNYWNPIHHEDQKTFNKEDLRKLLSKHNDMMIDQDKKRKDEFKSYEMQKEHELREKRKNMTEEERKKDEGEYKQHHNKKHETMHEPGHKAQLEEVWEKEDGLDPDSFDPKTFFNLHDKNGDGYLDIFELETLFLTDLDKNYNESDPDVDLKERDEELARMREHVMKSMDRDNDGLVSKEEFMMETKKDDFEKDDEWKPLTEEDQYTDEEFDEYEKMLQQDGERHGDEHHEDDKEKEHPSNNNSEGH*

>Cnidaria_H.vulgaris_Sc4wPfr_518.g19401.t1

MKELLTFGIFLALISLHVAPPVGKKEPELPANETGTVEDKQDAEYFRYLSQVVEVLEKDPEFKSKLHNASEEDIRSGKIANYLDLVGHGVRLKLDEIKRTEVEYQRELLRQRQDFMSGIERNYWNPIHHDNKDSFEMEDLKKLLSKHNDMMSAQDAKRHEEFKAYEMEKEHERREKLHNMTAEERAKEEALYKEHREQRSKHEKIHEPGHKAQLEETWEKEDGLDPESFDPRTFFNLHDKNSDHYLDLYELETIFLADIDKVYNESNPEVDLRERSEEIERMREHVMKNMDKDKDGLISFTEFMDETKSEDFEKDEDWKPLTEQDQFTEEELQEYEKMLSENPHGENVKQEASH

>Cnidaria_H.vulgaris_Sc4wPfr_518.g19401.t2

MKELLTFGIFLALISLHVAPPVGKKEPELPANETGTVEDKQDAEYFRYLSQVVEVLEKDPEFKSKLHNASEEDIRSGKIANYLDLVGHGVRLKLDEIKRTEVEYQRELLRQRQDFMSGIERNYWNPIHHDNKDSFEMEDLKKLLSKHNDMMSAQDAKRHEEFKAYEMEKEHERREKLHNMTAEERAKEEALYKEHREQRSKHEKIHEPGHKAQLEETWEKEDGLDPESFDPRTFFNLHDKNGDSYLDMYEIETVFLADIDKVYNESNPEVDLRERSEEIERMREHVMKNMDKDKDGLISFTEFMDETKSEDFEKDEDWKPLTEQDQFTEEELQEYEKMLSENPHGENVKQEASH

>Bilateria_L.anatina_comp148428_c0_seq11.p1

MLSLNLLAVLVIGLLVQFIVCPPVQKQVKEEEPVEGPEEDALPENLEYERYLKEVVDALEQDENFRKRLESANVSEIKSGEIAKHLDFVHHSVRTKLDEIKRREIGRLKEAQRLRMREMNGVGKVNFDDKKHLDHENPHTFEVEDLAKLIKQATTDLEEADKKRKEDFKKYEIQKEVLEREKLKNMPEEERKKEEQKIKEMEQKHKEHPKVHHPGSKEQLKEVWEEEDHMDPNDFDPATFFHMHDTNGDGVLTPDEVEALFQKELDKVYDPNAPEDDMVEREEEMQRMREHVYKEIDKNRDAMISMDEFLDSTKQAEFQKDEGWKTLDEQEPEYSQEELEQFQREYEEQLARHHAQPGEINVPPMDHAQQQGVQHMGQQPQPGHHPDQFGQPPHPGQFEQPPHPGQFQQPPPHPGQFQQPPHPDQFQQGHPNMIGSEHAGQPQIGHQQPQQGQKPEQGQPKVVVGQPQKVQQQPQQHVQAQQQVQQQAQQPQKTAQTQEQKASPQQQAQQQGQQQVQQQAQQQGQQQVQQQAQQQAQQQVQQHAQQQAQQQAQQPAQQQAQQLAQQQAQQQAQQQQQPMQNLQQQAQHGQVQQEQPQKQQ*

>Bilateria_L.anatina_comp148428_c0_seq2.p1

MLSLNLLAVLVIGLLVQFIVCPPVQKQVKEEEPVEGPEEDALPENLEYERYLKEVVDALEQDENFRKRLESANVSEIKSGEIAKHLDFVHHSVRTKLDEIKRREIGRLKEAQRLRMREMNGKQILHNVVYHGGVGKVNFDDKKHLDHENPHTFEVEDLAKLIKQATTDLEEADKKRKEDFKKYEIQKEVLEREKLKNMPEEERKKEEQKIKEMEQKHKEHPKVHHPGSKEQLKEVWEEEDHMDPNDFDPATFFHMHDTNGDGVLTPDEVEALFQKELDKVYDPNAPEDDMVEREEEMQRMREHVYKEIDKNRDAMISMDEFLDSTKQAEFQKDEGWKTLDEQEPEYSQEELEQFQREYEEQLARHHAQPGEINVPPMDHAQQQGVQHMGQQPQPGHHPDQFGQPPHPGQFEQPPHPGQFQQPPPHPGQFQQPPHPDQFQQGHPNMIGSEHAGQPQIGHQQPQQGQKPEQGQPKVVVGQPQKVQQQPQQHVQAQQQVQQQAQQPQKTAQTQEQKASPQQQAQQQGQQQVQQQAQQQGQQQVQQQAQQQAQQQVQQHAQQQAQQQAQQPAQQQAQQLAQQQAQQQAQQQQQPMQNLQQQAQHGQVQQEQPQKQQ*

>Bilateria_L.anatina_comp148428_c0_seq4.p1

MLSLNLLAVLVIGLLVQFIVCPPVQKQVKEEEPVEGPEEDALPENLEYERYLKEVVDALEQDENFRKRLESANVSEIKSGEIAKHLDFVHHSVRTKLDEIKRREIGRLKEAQRLRMREMNGVGKVNFDDKKHLDHENPHTFEVEDLAKLIKQATTDLEEADKKRKEDFKKYEIQKEVLEREKLKNMPEEERKKEEQKIKEMEQKHKEHPKVHHPGSKEQLKEVWEEEDHMDPNDFDPATFFHMHDTNGDGVLTPDEVEALFQKELDKVYDPNAPEDDMVEREEEMQRMREHVYKEIDKNRDAMISMDEFLDSTKQAEFQKDEGWKTLDEQEPEYSQEELEQFQREYEEQLARHHAQPGEINVPPMDHAQQQGVQHMGQQPQPGHHPDQFGQPPHPGQFEQPPHPGQFQQPPPHPGQFQQPPHPGQFQQPPHPGQFQQPPHPDQFQQGHPNMIGSEHAGQPQIGHQQPQQGQKPEQGQPKVVVGQPQKVQQQPQQHVQAQQQVQQQAQQPQKTAQTQEQKASPQQQAQQQGQQQVQQQAQQQGQQQVQQQAQQQAQQQVQQHAQQQAQQQAQQPAQQQAQQLAQQQAQQQAQQQQQPMQNLQQQAQHGQVQQEQPQKQQ*

>Bilateria_P.ijimai_comp94105_c0_seq1.p1

MISYWRLIALCAALLPVVLSAPVKPNVEVPVEKKEGEDDAATNLEYDRYLREVVSILETDPTFKKKLEEANITDIKSGKIARHLELVQHGYRSQLDEIKRKEIDRLKKYHKQQVELQNVGNLKIPEHIDHRNSETFEEEDLRKLIMKATADLDAVDTKRKEEFKDYEMEKEHERREKLKGMNEEQRKAEEKRMEELKKKHAEHPKINHPGSKDQLNEVWEEKDGMKGEEFDPKTFFNMHDANGDGSLDPIELEALFEKELEKSFDDKNPEEDDPVERDENRQRMREHVLKEIDTNKDHMVSLEEFLKSTKNEDFDNKEEWKQDVEDQEHLFTDDEFDEFEKENGYEDLDEHGQVEGDHDKVPPAGDHQQPPVVSDSHPNQQPQGHEGQA

>Filasterea_Tunicaraptor_GIQG01079875.1_.p1

IRFNIFGFFLQTHLSIGTRITWREIKPAKMRRAGGCPLLLLVLTALFQILMKSTLAAPLKAENSVEEMPEFVRYIEEMLQADPHLRAEFEKRFGGGPVAKAHPAELRELIETVQKRGLGGDAALDQLDVRSPSEEHERLVRERMRTEYRRRRERELGLAEGSELPEVVESEIEERHQLLEELDEIRRDGYKQRLMERERQRRAELEGLGPDEHKRREEAATRAKDAAHPQVHHPGSREQLEEAYENKGFDEDDFDPKTFFKLYDINSDDFLDAEELETIFVHEAQKIHPGEDSASREDLQEEIARMREHVVGEVDTDKDSRVSWDEFLVATEADDFEEDPGWDELNEDEEFDELEFEEYRARWEEEQRAKAAHDELPPLPDDMDPEERAVAEQEREHLRAGNVDGLPDHEFVPAKHPDEVRARAGEESED*

>Choanoflagellata_M.brevicollis_A9V878

MGRLSVLRLSMAWVALVMLLAAAYAAPVGTPKEEPAPPAPQDTQPSPPDPASEDLPDIHLAADVVDYDEYDDEDDQYYQEQEQEQQEQEQEQQEQQEQQAQALGNAKADQAPVRAEPKPKKLWPSVEEMRLLRVKEFDKEYERYEEETADQLVKDPNMAKSVLEALRANKDDSVAEKAVEAQVAQKLQRGFRSRLDEQPRLAGQAQAIFLGMLNTCLSSTGTGLIRCAISQKRQVIERMRRAAKRADNTEPERPDQAKFGDKGEGSHNEALVRNVVEKRVRLMGQLDERSRQAFVEHMMQRELEFRERMKATTNAADRRRLNEEHQKAWADLKKRRDFQPGHRDQLQEVWDNIDGMKGQKFTPKVEALVNAEATRLHTIEDFVDEQAVAYEAVRMRQAFMADVDTNKDDMISLQEFMLFAASNSFADSKNWTAIVPEFDDASLEKFRERVEALKEDRDGGAVPQGVAETADLVQARLAKERSDGFKRLQKMQADHQERIKTRRAQKMAHLARKKRAAQEPPAE

>Bilateria_A.planci_gbr122_8_t1

MERWRSLVLLLALLAVCRAAPVLPEKTDTDDEGDIEDSEHTGLEYDRYLRKVVKTLENDPVMKKKLDDLSLDDLKTGKMMEILDGVSDKMRARLDDLKRSEIQRLRQVARQRFQQANHRPMDQKQIEDMTGHLDINNMDQFGGKDFEKLIKKATEDLDKADKERKEEFKRYEMQKEIERRQKLKEMDEAKRKEAEQEHEERRKKLKEKHMKHPGSKPQLEDVWEETDHLEREDFNPRTFFNLHDTNGDGTLDAFELEALFVKQVEKMYAERHNSDPREKFEEISRMREHVMNEIDKNKDKMVSRKEFMEATEAEDFDEDKGWDDLEEQDIYTEDELETYEKNIQEEMKKLKLRLQQEHDQYKENRVPQPGDPVVMNQAKAAISNPQQLADQVIQAAADHEAEKFHQASQQMKKLVDEVANQVKANAQGGQKQQDNTQHQQQQQQ

>Bilateria_S.kowalevskii_30011267m

MASWKILLLLAALLPLIFAAPLDPKKQRESGEEEEQEKDDETDDIREQEEDYDTGLEYDRYLQEVINVLEKDPDFKKKLEEADIEHIKSGRIADELNLVDHQVRSKLDELKRHEINRLRLVAREQLDQKNGVKRMDKKSITDLLGHVDLGNTKSFEEEDLAKLIQKATKDLDEMDRRRKTDFKKYEMEKEIKRREKLKQMSDAKREQAKQEWEESQKKRKQHDKVKHPGSKKQLEEVWEETDHLDKDDFDPRTFFMLHDKNGDGKLDPMELEALFVNEIEKIYGDDSDPREKMEEMSRMREHVMKEIDTDKDFMVSMREFMDATRDKDFDNEESWEDLGDKDLFTEEELADFQRQLQEMEARRMRNNVKKQQQQQLGGEDDKLKFQAEAVAQVAAQQAAQQKSAQELAQQQKDAQLRNAQEAARQNQQAAQQQTEQKEQMKQQFDEAHAAIQQQHHQQQQQFDEAREAIKQQHHQNDANVVVMEGGQPLQAQAAQVNENKPQQ*

>Bilateria_B.belcheri_233850F.t1

MNKEWKKRRQHPFIFLLNTRMLTLRRCLLLAALVLLVSHDAMLAPVDPKKQQVEENETEKKEESQDTGLHYDSYLQEVVRVLETDPEFKKKLQEADIEDIKSGKLSAELNLVAHNVRSQLDELKRTEVTRLRKLQRVRMDMQNGRNGLHQGDLRDVKKMYEEFNHVDHSNPRSFEEEDLNKLIQAVVQDLENYDQKRHLEFKRYEMVKEHQRREKLSQLSEEDRKKEQERYEEMQRKHKDHPKMNHPGSKDQLEEVWEADGLAKEDFNPRAFFGMHDTNGDGYLDPMELEALFEKELEKVYKESNEEDDLREMEEERARMRKHVLREVDTNKDSMVSFEEFSDATKRKEFEQPEEDSFEPLDETDFFTEEELEEFEQQLAREADDLRQKEVELARQRAEHQQKRQQLQQVKMKQKQVVQMSQEQERQAEAQGQQVAVETNQESDQAAAGQDAVPAEVAQHAADGEGGQPAEMHEPVAAAVAADAVAVGGNLPPEGQLGEEQVQHQEAQQPAS*

>Bilateria_D.rerio_Q6NX06

MWRPSHCCLILALWACLDALPVAVDKTKVSPPAEAQEPPENADTGLHYDRYLREVIDFLEKDPHFREKLHNTDMEDIKQGKLAKELDFVSHNVRTKLDELKRQEVNRLRTLIKAKQDLNGEKGMTVDHQALLKQFEHLNHMNPHTFEVEDLDRLIKSATNDLENFDKERHEEFKRYEMMKEHERREHLKTMNEEERKKEEEHYEEMKKKHADHPKLNHPGSQDQLKEVWEEADGLDPNDFDPKTFFNLHDTNGDGYFDEQELEALFTKELEKIYDPAQEEDDMVEMEEERLRMREHVMNEVDTNKDRLVSLEEFITATNKKEFLEPDEWETLDQNPVYTEEELREFEQHLAREEQDLNLRTNDLQKQREELERQQDQLNAQKMELQQAVQHMERLKAQKTEPPVQPKALSAVEILPGDGQQLSQDLPPHS

>Bilateria_D.rerio_A0A2R8RUA0

MCHLNICRLFQMSCLKGLLTSCLVLASVLSWAQAVPISIDKTKVKIPEETVKEPPQSVDTGLHYDRYLREVIDFLEKDQHFREKLHNTDMEDIKQGKLAKELDFVSHHVRSKLDELKRQEVSRLRTLIKAKQDIEGGNDIAVDHQALLKQFEYLNHMNPHTFEVEDLDRLIKSATKDLENYDKERHEEFKKYEMMKEHERREHLKTLDEDGRKKEEEHYEEMKKKHADHPKVNHPGSKDQLKEVWEEADGLDPEDFDPKTFFNLHDTNGDGFFDEQELESLFTKELEKIYDPTNEEDDMVEMEEERLRMREHVMNEVDSNKDRLVSLDEFLVATKKKEFLEPDSWETLEQNQAYTEEEMREFEEQLVRQEEDLNQKAADLQKQREDLERQQEQLNAQKIELQQAVEHMERLKTQKVPLPSEILEGNAVPESVGQDQPPVPLEHQPLPPGHQDVPPPAAQQHHDELQNQQALGQEHQNTPHENQPLPPGHNNPPEDPPLVPLDHNIVP

>Bilateria_G.gallus_F1NGB1

MKWQSLLPQQCILLIPCLLMALEAVPIDIDKTKVKGEGHVEGEKIENPDTGLYYDEYLRQVIDVLETDKHFREKLQTADIEEIKSGKLSRELDLVSHHVRTRLDELKRQEVARLRMLIKAKMDSVQDTGIDHQALLKQFEHLNHQNPDTFEPKDLDMLIKAATSDLENYDKTRHEEFKKYEMMKEHERREYLKTLDEEKRQREESKFEEMKKKHGDHPKVHHPGSKDQLKEVWEEADGLDPNEFDPKTFFKLHDVNNDRFLDEQELEALFTKELEKVYDPKNEEDDMVEMEEERLRMREHVMNEVDINKDRLVTLEEFLRATEKKEFLEPDSWETLDQQQLFTEDELKEFESHISQQEDELRKKAEELQKQKEELQRQHDQLQAQKQELQQVVKQMEQKKLQQANPPAGPAGELKFQPPGEHKIEEAPKHPAGGDQPLPPGHIQEPAARTDQVHP

>Bilateria_H.sapiens_P80303

MRWRTILLQYCFLLITCLLTALEAVPIDIDKTKVQNIHPVESAKIEPPDTGLYYDEYLKQVIDVLETDKHFREKLQKADIEEIKSGRLSKELDLVSHHVRTKLDELKRQEVGRLRMLIKAKLDSLQDIGMDHQALLKQFDHLNHLNPDKFESTDLDMLIKAATSDLEHYDKTRHEEFKKYEMMKEHERREYLKTLNEEKRKEEESKFEEMKKKHENHPKVNHPGSKDQLKEVWEETDGLDPNDFDPKTFFKLHDVNSDGFLDEQELEALFTKELEKVYDPKNEEDDMVEMEEERLRMREHVMNEVDTNKDRLVTLEEFLKATEKKEFLEPDSWETLDQQQFFTEEELKEYENIIALQENELKKKADELQKQKEELQRQHDQLEAQKLEYHQVIQQMEQKKLQQGIPPSGPAGELKFEPHI

>Bilateria_L.chalumnae_H3BBJ0

MLHYFLLICLLVVSDAVPIDIDKTKVKDVPPVEAEKIETPDTGLYYDRYLREVIDVLETDKHFREKLQTANLEEIKSGKLSKELDLVSHHVRTKLDELKRQEVARLRMLIRAKMDAQQELGVDHQALLKQFQHLNHHNPDTFEPKDLDMLIKTATNDLENYDQERHEEFKNYEMKKEHERREYLKTLDEEKRKEEETRYEEMRKKHSEHPKVNHPGSKDQLKEVWEETDGLDPNDFDPKTFFKLHDTNSDGFLDEQELEALFTKELAKVYDPKNEEDDMAEMEEERLRMREHVMNEVDINKDRLITMEEFLKATEKKEFLEPDSWETLDKQQLFTEEELKEFEQHISNQENELQKKAEDLNKQREELQRQQDELQAQKEELQQVVQQMEQKKIQQGLPPSGPNGELKFHPPGEQSVQNVDPHPVGGNLPLTGGHMPVQEVQKEGATGAEQSNLHP

>Bilateria_M.musculus_P81117

MRWRIIQVQYCFLLVPCMLTALEAVPIDVDKTKVHNTEPVENARIEPPDTGLYYDEYLKQVIEVLETDPHFREKLQKADIEEIRSGRLSQELDLVSHKVRTRLDELKRQEVGRLRMLIKAKLDALQDTGMNHHLLLKQFEHLNHQNPNTFESRDLDMLIKAATADLEQYDRTRHEEFKKYEMMKEHERREYLKTLSEEKRKEEESKFEEMKRKHEDHPKVNHPGSKDQLKEVWEETDGLDPNDFDPKTFFKLHDVNNDGFLDEQELEALFTRELEKVYNPQNAEDDMIEMEEERLRMREHVMSEIDNNKDRLVTLEEFLRATEKKEFLEPDSWETLDQQQLFTEDELKEYESIIAIQENELKKRAEELQKQKEDLQRQHDHLEAQKQEYHQAVQHLEQKKLQQGIAPSGPAGELKFEPHT

>Bilateria_P.marinus_S4RIM9

MKCLLQMRMALESRALALTLLLALLSFVSAIPIDKSKLKEHNEVKDTKDEKQIEDTGLYYDRYLREVIDVLEKDTHFKQKLETADIEDIKNGRLSKELDFVSHHVRSKLDELKRQEVARLRMLIKAKMDAAQGKDLRMDHQSLLKQFEHLDHNNPHSFEPQDLEKLIKTATKDLENYDAARHEEFKRYEMLKEHERREFLKTLDEERRQEEEKRFLDMKQKHKDHPKVNHPGSKDQLQEVWEESDGLDPEQFEPKTFFKLH

>Bilateria_X.tropicalis_A0A803JHK8

MGTGQVRWGRMSLICSSLLLLTSLAAVSAVPIDKDKAKVKVEPMEEEQKSQSADTGLFYDRYLREVVEVLETDRHFREKLQTADIEDIKSGKISKELDLVSHHIRTKLDELKRQEVARLRMLIRAKMDQGDDTGMDHRALLRQFEHLNQNNPHSFEPQDLDLLIRTATKDLENYDKTRHDEFKRYEMMKEHERREYLKTLDEEKRKEEEEKYEEMKKKHREHPKINHPGSKDQLKEVWEETDGLDPTDFDPKTFFKLHDTNSDGFIDEQELEALFTKELEKVYDPNNEEDDMVEMEEERLRMREHVMTEVDLNRDRLIDLQEFIRATEKREFLEPDGWETVADQPIYTEEELQEFEKQILQQEEELKRKADELYRQREDLQKQHEYIQAQKQELQQVVQQMEQKKLEQQQPVPPAVPDANFQQGGDQAVHLGSQQPLPPGHLPEAGVPQSHPVP

>Cnidaria_A.millepora_XM_029344017.1

HWNQRMDIRSLVVPRFGKQPRALPLRSNSQSENCLKVPQFSWIRNTKDLNICCQKICPKMWQKATLICVFLVGLAVLCVAPPVRKGDKSDSKTDDVKGDPEYARYLKQVIEILESDDDYVKKLLNASEEELRTGKVADDLDLVKHDVRTKLDELKRQEVERQRMIRRQMNDHLNGLKEREYWNPLFDDENPNFFGPEDFKKLLWKHHEEMDRQDSERREEFKKHEMEKEHKRKEHLKELDDKARKEEELKFEQLQQKHLNASKNIHHPGSKAQLEQVWEETDGLDAKDFDPRTFFKLHDTNGDDYLDTAELEVLFVKEVDKLYDEKDEDYDPREKDEEIARMREHVMSEIDKDKDGFVSLEEFMKASKGDEFEKDEGWKTVEEEPGYTDEELAEFEKQLQEEERKHKAATETMQNFAEEVKQVAGTLVEHGEGQQQKQEETHDGTQQQQQQQQQQQQSHDGKEQEHPSP*

>Cnidaria_C.hemisphaerica_TCONS_00052731.p1

MNKFSLVILLIFGLVQQHVAPPVEKEKKEFQDNMRGNTDGLSHEQAQDQEYFRYLTQIVGELEKDAKFKAKLNNASEDDIKSGKIADFFPEVDVSIRRKLDEIKRMEVEYQRDLVRQKKDHMAGIERNYWNPIHHDNKDSFEMADLKKLLSKHNDMMSEQDKKRKEDFKKHEMDKEHERREKQKNMSEEERKKDDEEYKQHHEQKHESMHEPGHKKQLEEVWEKEDGLDPDSFDPRTFFNLHGNDFHLKKHISI*

>Cnidaria_C.hemisphaerica_TCONS_00064061.p1

MDKEHERREKQKNMSEEERKKDDEEYKQHHEQKHESMHEPGHKKQLEEVWEKEDGLDPDSFDPRTFFNLHDKNNDGHLDTYELETMFLNDLDKVYNESDPNTDKQEREEEMERMREHVLKNMDTDKDGLVSYDEFIAETKDTDFDKDEEWKPLTEEEQFSDEEYKEYERLLSEQEGHHEDENHVEGGAQAEVHH*

>Cnidaria_C.hemisphaerica_TCONS_00064062.p1

MDKEHERREKQKNMSEEERKKDDEEYKQHHEQKHESMHEPGHKKQLEEVWEKEDGLDPDSFDPRTFFNLHDKNGDGYLDTYELETFFLSDLDKVYNESDPNTDKQEREEEMERMREHVLKNMDTDKDGLVSYDEFIAETKDTDFDKDEEWKPLTEEEQFSDEEYKEYERLLSEQEGHHEDENHVEGGAQAEVHH*

>Cnidaria_C.rubrum_TR81664_c0_g1_i1.p1

MTWTHTGKQLAFVFIFVLIITVVHCKPVRKAAKKVDEEWKKEEDPSYVKYLKQVIEILEGDENFAKELENVTEEDIRSGKIAEQIDFANHRVRTKLDELKRKEIESQRQLLRQQQDLRKGIHREYWNPLNDKENPNTFEVGDLKKLLSKHHEEMDEIDKKRKQDFQKYEMEKEHERRQRDKVLDEAKRKEKEEERKRLREKHLQDAKNINHPGSKSQLEEVWEEEDGLEKDQFNPRTFFNLHDRDGDGYLDPMELESLFVREASKIHNSSNADYDPRKLNEEVQRMREHVVKEIDKDKDNFVSYDEFIGATKEKDYEKPPEEHWKGIDDEDEYTDEEFENYEKEFYDDEDEDAVPGHEEPPHPDQEKREDESNTAKHVPMAHNQEQEKREHQQDHQQEQQHQQ*

>Cnidaria_N.vectensis_comp283_c0_seq1_.p1

MDSKLSLLSLLVVLVLVTLCYAPPVKRGKPAEKAPEPNKDDPEYARYLRQVIEILEKDEDYVRKLMNASDDDLRSGRIAEDIDLVKHDVRSKLDELKRQEVERQRMIRRQMNDHLNGIKEREYWNPLFDDENPDFFGADDFKKLLWKHHEEMDKQDRERREEFKKHEMQKEHERKIKMKDMDEKHRKEAEEHFRELQEKHHNQSKQLHHPGSKAQLEQVWEESDGLDAKDFDPKTFFKLHDVNGDGFLDTGELEALFVKEVTKLYNPKDEDYDPKERDEEISRMREHVMNEIDKDKDGFVSQDEFLQSSKGEEFDKDDGWKSVEDERPYTDEELAEFEKSLEQDQHPSEQETDHEQPAQHDQQQQNTGQQQQHDKQQQHDQQQQHDQQQQHDQQQQHDQQQQHDQQ

>Bilateria_D.rerio_F1QLH9

MTGSFKALLLLSLCLLVWAVPIDRNPDPPQEEKAEENVDTGLYYDRYLREVIEVLETDPHFREKLQTANTEDIKNGRLSKELDLVGHHVRTRLDELKRQEVSRLRMLLKAKLDSTNAQSVQMDHASLLKQFEHLDPHNQNTFEAKDLELLIATATKDLENYDAERHEEFKRYEMLKEHERREYLKSLDQEKREKEEKRMDELKEKHRQHPKVNSPGSVDQLREVWEETDGLDPQEFNPKTFFKLHDTNSDGVLDVQELEALFTKELEKVYDPKNEEDDMVEMEEERLRMREHVMQNVDVNHDRLVSLEEFLKSTEKRELNNQKEWETLDDTKPVYTEEELQRFETELRDKELELGRRAEKLRQEQELLKERSKALEAQKREYQQAVMEMSQKQKEQQALNKQPPSGPNGELKFQEGIQKLINEDSQVKEPDQQTGDVQNNLPAEPPQNLPQHTS

>Bilateria_M.musculus_Q02819_NUCB1

MPTSVPRGAPFLLLPPLLMLSAVLAVPVDRAAPPQEDSQATETPDTGLYYHRYLQEVINVLETDGHFREKLQAANAEDIKSGKLSQELDFVSHNVRTKLDELKRQEVSRLRMLLKAKMDAKQEPNLQVDHMNLLKQFEHLDPQNQHTFEARDLELLIQTATRDLAQYDAAHHEEFKRYEMLKEHERRRYLESLGEEQRKEAERKLQEQQRRHREHPKVNVPGSQAQLKEVWEELDGLDPNRFNPKTFFILHDINSDGVLDEQELEALFTKELEKVYDPKNEEDDMREMEEERLRMREHVMKNVDTNQDRLVTLEEFLASTQRKEFGDTGEGWKTVEMSPAYTEEELKRFEEELAAREAELNARAQRLSQETEALGRSQDRLEAQKRELQQAVLQMEQRKQQLQEQSAPPSKPDGQLQFRADTDDAPVPAPAGDQKDVPASEKKVPEQPPELPQLDSQHL

>Bilateria_X.tropicalis_A0A6I8QNX2_NUCB1

MKICTLLLFSFLGLGLAVPIERPTPKKEAEPPPPPEQPDTGLYYDRYLREVIDVLETDGHFREKLQAANADDIKSGKLSKELDFVSHHVRTKLDELKRQEVSRLRMLIKAKMDATMEENVQIDHMSLLKQFEHLDPQNQHTFEARDLELLIQAATKDLENYDAARHEEFKRYEMMKEHERREYLKSLDEEKRKMEEAHFEEMKKKHKEHPKVNVPGSIDQLKEVWEETDGLDPNEFNPKTFFKLHDTNGDGVLDEQELEALFTKELEKVYDPKNEEDDMVEMEEERLRMREHVMKNVDANHDRLVTLDEFLKSTERKEFNEADGWETVDETQVYTEEELRKFEQELSAQETALNQRAEELRKEHEQLQQQQIELDAQKKEYQQAVMQMEQKKAQQTGAEAAAGPNGELKFQPEAPHAESEHAKPVEEQPKLAEQQQKSDVHPAAPGEADGHPAGQKEPEETALNQRAEELRKEHEQLQQQQIELDAQKKEYQQAVMQMEQKKAQQTGAEAAAGPNGELKFQPEAPHAESEHAKPVEEQPKLAEQQQKSDVHPAAPGEADGHPAGQKEPEETALNQRAEELRKEHEQLQQQQIELDAQKKEYQQAVMQMEQKKAQQTGAEAAAGPNGELKFQPEAPHAESEHGKSRQTDICSDRT

>Cnidaria_A.alata_agalma_gene_56247.1_isoform_1.p1

MEKILILLVCLTFVSYGLSKPVPVEKTEEEKKGEADAEYLRYLGQVVQILEEDEEFKKKLHNASEDDIRSGRIVNDFEFLSHKVRTKLDELKRKEVEYQRQLQRQMRDHKNGLTERQFWNPVHDENQMSFEASDFHKLLQKHNEMMEEQDKKRREEFHHYEVEKEAKRRKHLESLPEEERKKELEEYEKQKKDRSQHEKLHEPGSKAQMEETWSEQDGLDPENFDPRTFFKLHDKNEDGYLDMYELETLFLGDVNKVYNESDPNVDPRERAEEIERMREHVVKEMDKDKDGLVSLQEFMDETKDPDYEKDEEWKPLTDQEQYSEDEINAYQKQLGEHDDPEPDDHAPETIGEHHEPEIMKHDEKKEEHHEPEKKEQKGEQHH*

>Cnidaria_P.hydriforme_GBGH01000495.1_.p1

DPSATEFVSRRTELDRMHWALLVMAVCLLRSSRAPPVTPSVPEIDLSKFLDKDESYRHYIQEVLSVLEEDPVYRQKLMTASIDDIVTGKLADDFHLIEHNIRTRLDEVKRLEVSRQRELVRLYADQVHGIKEREHWVDLDNQHIFSEQDFRKLLKKEATTIQSLDEKRHEEYVKHEMEKERLHQEHLKSLSEEQRKLEEEKEKNETEEQLKNAKLHEPGSKAQYEEVWEEEDGYEASDFNPKTFFMMHDINGDGYWDISEVEAVLMIEVSKLYNESDKDFDSVERDEELMEMRQHVFKEYDKDGDMLVSLDEFQKVSSQESFKEDEGWKSNLEQPEYSEEEYKKYMSEYEVNHAANATAPIH*

>Cnidaria_P.hydriforme_GBGH01003830.1_.p1

ATEFVSRRTELDRMHWALLVMAVCLLRSSRAPPVTPSVPEIDLSKFLDKDESYRHYIQEVLSVLEEDPVYRQKLMTASIDDIVTGKLADDFHLIEHNIRTRLDEVKRLEVSRQRELVRLYADQVHGIKEREHWVDLDNQHIFSEQDFRKLLKKEATTIQSLDEKRHEEYVKHEMEKERLHQEHLKSLSEEQRKLEEEKEKNETEEQLKNAKLHEPGSKAQYEEVWEEEDGYEASDFNPKTFFMMHDINGDGYWDISEVEAVLMIEVSKLYNESDKDFDSVERDEELMEMRQHVFKEYDKDGDMLVSLDEFQKVSSQESFKEDEGWKSNLEQPEYSEEEYKKYMSEYEVNHAANATAPIH*

>Cnidaria_R.esculentum_TR45503_c0_g3_i3.p1

MAHKTILLLLFCVLAINEIYGKPIRKEKEEKEHPKKEEEAEYLRYLGQVVEILEEDDEFKKKLHNASEDEIRSGAIADHIDLVSHKVRSKLDELKRMEIEFQKRLRRQERDHLNGINERNFWSPLFDENSDTFEQEDLKKLLHKHNIMMEEQDKKRREDFHKYELEKEHKRREEMKNLSEEDRKKKEAEHEDELKKLRKHEKMHEPGSKAELEEVWKEEDGLDPESFDPKTFFNLHDKDGNGYLDNFELETFFVDDVDKVYNESDPNTDPRERQEELERMREHVMKQMDKDNDGLVSKEEFIKSTNDPEFEKDDEWKPVVDDEEYSEDELKEYEKQLEEEAERNREGHHGENNEHESDEHHEKPEEVHVEEKKDEHHAKQEQAQQQHHEQPKQQQQQHDQQQQQHNEQQQQQTQQQPQHHEQQQQQQHEQQQPGSH*

>Cnidaria_T.bryosalmonae_contig_12406_1_1182

RQAAILTLWTLSLLGYIVCPPVVPSDDSKAAPASDAQQLANPLDDLEYNKYIQEVVNVLDQDDEFRKRLEAANASSSEAGSVAMSLEFANKNTRTRLDELKRVELRRLQELHIRLLRQQAESIRSGKHEGKLDLEKVDMEHVDHANPHSFEVEDLKKLIKKTTEDLHKIDSQRREEFKRYEMLKEHEYREKLKHLNDTQRSLAEHEHEASVKRHEQHPAVNHPGNKEQLQEVWQGVDHMDKNEFDPNTFFYLHDLNGDGYLDVGELDALFQKELDKVYSPGNPDDDPVMRNEEMNRMREHYFNETDADQDGLISRQEFLKSTESKDYQKKDEWETIEKQPAPFTDEEFKQFEEQIAASELKEKVDEHIKGAQAAEGIVHAASKLQPGHRQEAR

>Ctenophora_M.leidyi_comp17847_c0_seq1_.p1

LYTNNLSMKLLIASFLLLFLAAEGAENPESVIPTKTEYERYLDQILLVFEKDPKVKEILKNATPQDIKNGVVEKTLRRLHPDIRSKLDEVKRNIIMEYRQKLRRAKMDDKVPDDLYKFQDEDLKHLLKERHDRLAQMNDERKRQFKKYEMEKELKHRMALRAMNEYDRKVQEALDEEQKQKKMEKVKNLKHPGNKAQLEEVWENEGFDKGTFNPKTFFKMHDLNNDGELDDFELEAIFEKEVEDLFESFEDADEEEMAEEVAQMREHVINHVDKDKDGVIQLKEFIEYANGSEFDHEDPWDAAFNAEFTDQDLDEYEKTLDDIEREDVKAQAEIEKQKRLETIKKYEEESKINMEMLKAHSEKLKEALMAANKAKDLEKAKKLVAEAQADQEANKASP*

>Ctenophora_P.bachei_comp48688_c1_seq10.p1

MIRIWTILCLTIVSVACSESVIPTKSEYERYLDQVLLVFSHEPQIVNLLKNASPEDIKNGVIEKTLRRINPQIRSRLDEVKRNIIMEYKQKLRKAKMDDNVPEELYTFTHDDLKHLLNERHERLKQMDDQRRKEFKKFEMEKELKHRMALRAMNEYDRKVQEALDEETQTMKKDKGQKLKHPGNKEQLQDVWEAEGFDRETFDPKTFFRMHDMNGDNEWDLFEMEAIFEKEVEELFDGLEDVDEDEKYEEVARMREHVVEHMDTSGDGFIQLEEFVEYSKGKDFDHNDPWETVFDDNLFDEAELSEFEKNMDKTDTKELEELAKKEALTREEKIKKYEDESKVNMDMLKEHSEKLKEAIMAANKFKDLEQAKKIVADAAAGEVS*

>Ctenophora_P.falcata_comp27370_c0_seq11_.p1

SVIPTKTEYERYLDQVLLVFSNDPQVVNLLKNATPEDIKNGVIEKTLRRINPQIRSRLDEVKRNIIMEYRQKLRKAKMDDNVPEELYTFTNDDLKHLLNERHERLRQMDDQRKKEFKKFEMEKELRHRMALRAMNEYDRKVQEALDEESKTKKKEKAQNMKHPGNKEQLQDVWESEGFDRETFDPKTFFRMHDMNGDDEWDLFEMEAIFEKEVEDLFEGLPDVEEDEKYEEVARMREHVVEHMDTSGDGFIQLSEFLEYSKGKDFDHNDPWETVFDDQLFDEAELTDFEQKLDAGETKDLENLAKLEAAKREEKIKKYEEESKINMEMLKEHSEKLKEAIMAANKFKDLEAAKKIIADAAKEGGASGDKPAT*

>Ctenophora_P.falcata_comp27370_c0_seq20_.p1

SVIPTKTEYERYLDQVLLVFSNDPQVVNLLKNATPEDIKNGVIEKTLRRINPQIRSRLDEVKRNIIMEYRQKLRKAKMDDNVPEELYTFTNDDLKHLLNERHERLRQMDDQRKKEFKKFEMEKELRHRMALRAMNEYDRKVQEALDEESKTKKKEKAQNMKHPGNKEQLQDVWESEGFDRETFDPKTFFRMHDMNGDDEWDLFEMEAIFEKEVEDLFEGLPDVEEDEKYEEVARMREHVVEHMDTSGDGFIQLSEFLEYSKGKDFDHNDPWETVFDDQLFDEAELTDFEQKLDAGETKDLENLAKLEAAKREEKIKKYEEESKINMEMLKEHSEKLKEAIMAANKFKDLEASKKIIADAAKEGGASGDKPAT*

>Ctenophora_P.falcata_comp27370_c0_seq4_.p1

MIRVWVILCLTLTLVAGNESVIPTKTEYERYLDQVLLVFSNDPQVVNLLKNATPEDIKNGVIEKTLRRINPQIRSRLDEVKRNIIMEYRQKLRKAKMDDNVPEELYTFTNDDLKHLLNERHERLRQMDDQRKKEFKKFEMEKELRHRMALRAMNEYDRKVQEALDEESKTKKKEKAQNMKHPGNKEQLQDVWESEGFDRETFDPKTFFRMHDMNGDDEWDLFEMEAIFEKEVEDLFEGLPDVEEDEKYEEVARMREHVVEHMDTSGDGFIQLSEFLEYSKGKDFDHNDPWETVFDDQLFDEAELTDFEQKLDAGETKDLENLAKLEAAKREEKIKKYEEESKINMEMLKEHSEKLKEAIMAANKFKDLEASKKIIADAAKEGGASGDKPAT*

>Bilateria_A.ventricosus_A0A4Y2D5N5

MSNLLFTLLLVLVALELAVAPPVDQKKKDEEKKDKGEELDDFGLEYGRYLQQVVQALEEDKEFAKKLENISAEQIRSGHIAKELEFVKHNVRSKLDELKRIEVDRLRKLIQEQMERTELGLDRKGVKIPKHLDIENPHSFEIDDLKKLIQTATSDLEELDRQRREEFKKYELEKEATYRESLQNMTEDQRKEAEKHHQETIAKHKDHPKVHHPGSKPQLEQVWEDQDHMPREEFNPRTFFAMHDINGDGHLDVEEVEAILIPEVKKLYNPNNEEDDPVEMEEEFHRMREHIFNETDKNKDSLISLEEFLQMTQQQEFQRDEGWRGLDEQQVYSDAELQEYMRQRQMVLLFI

>Bilateria_C.sculpturatus_XP_023229830

MNRSQFVFLLIFALLAIHLVGAPPVDPKKKKEKEKEHEEDEGNELDDYGLEYGRYLQQVVQALEEDKDFAKKLENISTDQIKSGDVAKELEFVKHNVRTKLDELKRIELERLRKLSREEMERLELGLYRNEDGHLAPPSIDGRRWRTISSAGDGEGINRKHIKVPHHLDVDNKHTFEIEDLRKLMKAATRDLEELDRRRREEFKQYELEKEYQFRESLKNMSESEKAEQMKKHEEMLKKHKDHPKVHHPGSKQQLEQVWEEQDHMQGNEFDPRVFFAMHDVNGDGYLDEQEVEAILSLEIKKMYDPQNNVEDDPQEMMEEYNRMREHVYKETDKNKDGLISKEEFMDMTKRADFEKDDGWRGLDEQQIYTEEELKAYEKKRIEELQKLNAYYGYQNAHGGAPVPHGMPQYQGVPPPHPGYQPHPQQMPQGYQPHPQQMPQGAHQQGYQSHHQQAQMGGAHPQGYQSQQQGEAHHQPSQQQQGYQPQMQQMPQPGAHQIPQPGYPPHQQQMPQGIHQPMPGMQQQPHPGAQYGIPNNQNPQMRHPAAQGIHPGVQIVQGNAHGLPTHAPSDPPPRQPQPTQYNPQQRQPAVEQKQAEDTNANTANQKQGN

>Bilateria_D.pulex_E9GUA2

MELEMQQETGLEYNRYLQEVVQLLESDPDFRQKLEKSDPEDIRTGKVAKELEYVNHHVRNKLDELKRQEMERLRHLAMEEYERARGLGIPHDGRLKIPGHLDHKSPSFESEDLRKLIVQTSKDLEEADKQRKEEFKEYEMQKEFEHQNKLKGLDEEKRKTEEKEWEEAQAKHKQHPKMHHPGSKQQLEEVWEEQDHLSPDSFNPKTFFALHDLDGNGYWDPDEVKALFSKELDKAYDPNAPEDDMAERYEEMERMREHVFNETDTNRDYLISFPEFLEQTRKQEFERDPGWNTIDEQPVYSQQEYMEFEHQRQMEIQRLIDQGMLPPHPGMMPNHPRMPNVPYGVAPGQYAGQQPYYPSPVAQGKPITPEEAIRMQQQQYAQQPQFAPQQQYYGQPQFAQQQQYAGHPQQQFAANPQYAAQPQQPQFAQPEQQNFQQAAQQQQQNFQQASQQQQPIQQVPQQQPIAQQPVQQQQIPAQQQPLVQPQVPVQQQAPIQSQPVALPPAPQQIPAAKTV

>Bilateria_D.melanogaster_tr_Q9VVK7

MVQNVALLGLALIAISASIVALPVTQNKKDHKEAAESSTPATADVETALEYERYLREVVEALEADPEFRKKLDKAPEADIRSGKIAQELDYVNHHVRTKLDEIKRREVERLRELANQAYELSNDIDRKHLKVSQHLDHDNEHTFEIEDLRKLIQKTSDDLAEADRKRRGEFKEYEMQKEFEREAQKKEMDEESRKKFETELKEKEEKHKDHEKLHHPGNKAQLEDVWEKQDHMDKNDFDPKTFFSIHDVDSNGYWDEAEVKALFVKELDKVYQSDLPEDDMRERAEEMERMREHYFQETDMNHDGLISIDEFMVQTNKEEFQKDPEWETIDRQQQYTHEEYLEYERRRQEEVQRLIAQGQLPPHPNMPQGYYAAPPPGGVAYQQAPPGAQLHYQHPDQVHAQQQQQYAQQQQQYAQQYQQQQYGNGQQPVQLQPNQVYQHAGQIPQQQQPVYQNQPVYQQQQPVYQQQQPVQQQQKPVQQPVQQQQQPVQQQQQPVQQQQQTVQQQQPVQQQQQTVQQQQPVQQQQQTAQQQPVAQQQIHNQSPPPVLNQQVPVQQQQKQHQESLNQQH

>Bilateria_P.caudatus_XP_014669982.1

MARLTWLILICLVASIVCPPVDKNSKKSEEDKKKENGTTDNNDIDLLEYDRYLKEIIGALEEDGDFKKKLESADFNDVQSGKIANELQFVNHNVRTKLDEIKRREMERLRQTTKKMLQKKEGRDVLKQGDIGGHIDHMNPRGFDMSDLAKLIQKTIADLETLDTQRKDNFKTYEMEKEYHHKEQVNNMTEEEKKIAEQEYLQQQEKHNDHPRVHHPGSKQQMQDVWTDNDHMDPGSFNPKTFFSMHDYNRDGYLDQKELEALFQTELDKLYDPDNPEDDMYERQEEMARMREHVQREVDTDSDGLISLAEFIAETHRSEWGEDEGWKDIRDEPVFDNEEYRAYEQFMAEQRHLASMDPQRLNTVPPHILEDFAKLPPDQQQQVMLQQHLQNQQQQQHPGQYQQHPGQVDPAQVDPRYQQHPGQVDQQYQQHHDQVDPRYQQHPDQVDQQ

>Bilateria_T.castaneum_D6WHU5

MNRYIPFAFFLISVLQVCFAPPVTQNKDKDKEKGEDEIGGLEDYMEYHRYLQEVVNALESDPQFRQKLEKADETDIRSGKIAEELEFVSHHVRSRLDEIKRVELVRLKELTEKKRQLQQNIDLEDPSHHHLDHSNPHTFEIDDLKKLIAKTTADLAEADKKRREEFKQYELQKEFEKQEKLNHTNGEDREKLEREFREREEKHKKHEKLHEPGHKAQLEEVWKEQDQMQQEFDPKTFFMLHDIDGNGLWDQDEVKALFIKELQKMYAAGEPEDDMRERAEEMERMRESVFSEVDVNRDGFIDYEEFLAQTKRNDFQQDHGWQGLDERKPYTDEELEEYIRRHQAANQIPHGYPPQGYAQHPPPGYHPNVAVHPNGVPVQQYHPGQVPQQQFHPGQLSQQHPQQLQGHLPDLNTNEVYPQQRQQQYQQPPQHYQQQQYQQMNYQQHPQQVNYQQQHPQVPVQNQQQQHPQVPVQQNQQQHPQAVNQQQVPVQNQQQAANQQVPVQNQQIPVQNQQQQIPVQNQPQGVPQVQGNVQQGAPQASNPNLNNAQANQV

>Bilateria_L.gigantea_V4A928

DTGLEYDRYLREIVSVLEEDEDFRKKLEAANVSEIKDGSIARHLEFVGHNVRTKLDEIKRREVDRLQELARLKMKQMKGTEILHNIAYHGDIRIPGHLDVMNPHSFEMKDLESLIKKTTNDLEELDKQRRDEFKEYEMEKQYEEKEKLKHLSEEERKKEEERLAELKKKHGEHEKINHPASKDQFEEVWENTDHLDKEDFDPKTFFLLHDANGDGVWSIQEVEALLQHELDKAYDEKNADEDDPVERVEEMNRMREHIFTEIDQNHDKFITRDEFLIYTGPKGDRKEFEENEDWKVSCYSTNLV

>Bilateria_B.glabrata_A0A2C9KTG4

MVNHTVRTSLDEIKRREISRLQELARLQMQGMSAHDGVKKFEIPSYLDVRNPHSFEVKDLENLIIKTTSDLEELDKQRKEEFKEYEMEKTFEQQEHLKALKEEERKREEARLEELKKKHAQHPKVNHPGSKDQFEEVWEKVDHLEDQEFNPKTFFYTHDVNGDMEWSVDEVDAVLQLELDKVYDAKNSPDEDDPVERQEEMNRMREHVFQEMDKDKNWRISFQEFIDYTGSQH

>Bilateria_O.bimaculoides_A0A0L8HQS8

MFITILVSLVLFVINGIVCPPVKPGTKPKEGASVNDTNETGLAYDRYLNEIVTVLEEDPAFKKKLETSDLQYIKSGAISKDLDLVHHGIRQQLDEIKRREVIRLKSLVQGQFKYKSEERGNRKVALHHININNPDSFEQDDLAKLIKQATHDLDELDRERKEEFKQYEMKKEHEIKQKLANMNETERKIEVQHIKEMKEKHKQHPKIHHPGSKAQLEEVWEEVDRLNGENFNPKTFFKLHDVNGDGVLEPSEVEALFTREIEKMYDPNNPEDDMKEAKEEINRMREHVFKEADTDDNHLLDEGEFERLTQREEFGKDEGWKGLDEDDEIYSEEEMQQYEKMLREEEERLKKQQIYDNLAQPGQVVQPVDPNVVHMQQPHQEQQQQHQEQQNLQYAHQEKLRQQQLTDAQHQQQFAQQQAALHQQQQQQPVGHQQQQQPVGHQQQQPVVHQQQQPVVHQQQQPVVQQHQQQQPVVQQHQQQPVVQQQQQQHQPVVQQHQ

>Bilateria_P.dumerilii_comp411621_c0_seq2

MKLQSGKDSSIMEKLALTALLLVIVLPGIFCPPVTKTPEPSKSKNDHSTPAPEEDVNFGLEYERYLKEVVNILETDEIFKKKLEESNISDIKSGAIAKHLEFVSHAVRTKLDELKRQEMDRLRNLIKIKTRNKQGIPTHDLEKIVGHLDGDNHETFEENDLAKLVKKATQDLEDLDKKRKEEFKMYEMEKEHLKREKLKGMSEEERKAEEKRLEEMKEKHKKHPKLHHPGSKDQLEEVWEESDGLAKEDFDPKAFFMLHDTNSDGFWDELEVEALFQKELDKMYDPDAPEDDMMERYEEMSRMREHVLNEVDKNKDKLISMQEFIDSTKTGEFNKDDGWAGLDEEDSPYTEEELAAFEREQEQLGGHAQPGEVNFHGQGLAPPPLAPHLGGSPQFQQRPPGGFQPPPPMGHQGYPNQGYPNQGYPNQGYPNQPLQYQQPPPYNGAQPPPPYQQQPQYQQAPNIYQQRGGAPGGQQQQQQQQTQHGFQQPYQQQPPQGHQFQHQAPPAGQPVQFGQNNNQFQQNQGQNNKNNQFQGQNNQNNQFQGQNKQQQQGQVNSQGQNSQGQGQNAKQSGGQDPNSVNQQATGQPATKTLPKSQKQKQFDAANDFGRV

>Choanoflagellata_S.rosetta_F2U7V5

MRTKQAALLVLALVGVLAVVGCMQGMTGAHAAPVRTAVEDTDTDNAAAADAADAAESTAPPADDAPVDADSAQDTTDAGTDNGAAKPVPVSKDEPYKAYDPYDPYDYDDYDYGEYEELYNEQYDLYADEQLDLKDEESYVGGDEDDDADPDVMSVRKEAAEKRRKADEEEYQRYLDEMMEEMARDSDLREDLIEDIQGVQEAERAPGDKDILKIALRLKNEERVRNIIEEKERQLLEAQRHEARRRAEARMHVPRPHDAMPEGGEEEDEDLFLDPEAAFDKLTEIVKRRAAVLGEIDTKRRAAFLQKEMMRELEFRRSVAHEKDPEKKKQLIKSHKQKIGHAKQLNEVWEKQDGMQSKFNPKVFFSLHDINGDDFWDHKELEAIFHRSAVKLHTIDGETDTDAVEEEMLDMRAWVMKEVDTDKDQLVSRDEFITFANSPEFQTNRDWKAVFPDFNGTELKEFKDRARKAGDPRQEHVQARVAEASKRFKDKFAKTMKLGTNALKEAIKPFAKVGGNKKPAAQGDEGAAANEQQQQQQQQEQK

>Placozoa_T.adhaerens_evg1220999

MKKLVSVILLITVGLVILIDAAPPRLPVKADKVKEVPKEDVTNIEKTKIDVGVDKSVQLDSKMAHDQQNMDQHKTDQRNMDQHKTDQQNTDHQEPEENYDDTFQFLRDQPDLINKFKGLDDDVIGRDLANPAFRDLDEKMLDRFDDQEYDQVRHQRDLEHQIQEANEREEHPESFRLHHSDDNRYTMEELNEKLQEANKKLRETDDVRHKAFLRYELEKELLRKKELEKMNPDQKKVAEDNYAKHLQDMQDHPKLHHPGSKEQLQEVWEKEDGFQNQEFNPDAFFRLHDTNGDGHLDNVELEAVLRKEVSKLVGNHKDAYDERELKEELQRMRQHVSTEIDVNKDGLVSLDEFIKSTKKAEFKKDEGWETFNEDKDFERMTPEELERYERKVRDAARNQFDKIKIRTGEETDPRR

>Porifera_A.queenslandica_tr_A0A1X7UY00

MSSWVLVFLFLSVVSLSSSEASPFRNEKKEEGKRERPLHDTIMERMKKIRESLSMKGQDEEYKRYWEELIANNPDLAKRFREQKAEDLAAGLGDRLSHRDGGIRSQMEELQRRNIEELRDLHRQKMAGESHEPIEIRRQRERGRHLSDKEMEEKIREHQRIQEQLDHARREDFKNFEMKLEHRRRQKLKEMDETQRLREEARLKKLSDERKVHEPLLHPASKDQFEEVWKTDDGFKDEKFDIRTFFHLHDTNGDGTLDIREVEALFVNEIRKYYGKDFENDREAHEDMSRMREHVMSEVDRDHDSLITLEEFVRYANTDLFNIKEEWKPVGEDDEHKEFTDDEFKQYENEMEEDDYDYDDEGNLIHKPKMENHDGVPPEHHEVPTEHHEVPPEHQEVPTEHHEVPAAHDTKPDTGLPNDHGLPPNDHGLPNDHGLPPYDHRLPNDHGLPSDHGSPKGEKGLPEHVPPQDVPHDVPVGGANHNENPKAPIVETKQDDTAPPPHAASDDEKKFHFCYNQTLEGVEYSYSLGIPVFLNFAFRSQGACPSVAIKTEVYSPESYD

>Porifera_E.muelleri_m.21273

MFVCCIVVRSTCSFVVIHYVIPNPYNVKMPAGVMKAGLVVAALFVSVLSLPVDQRKDQPGKQDPFPGLDKEEFRRYWDDAVKNNPELAKRLGKMRPEDIPGAHPRLADKIAREFPDLRGQFDEARRMNIDELRQFQKVNNWKNGAGFPQQPPQRGQQQWPSMSGDPQWPDGRRGGGPQRGPQGGMPQWPGRHEADDLENRLKEKQKLLEEMDRKRKEDFKRYEMRMEHQQRAKLKEAQDEKERLKLEQELKDHREQILKHQRINHPAGRKQLEEVWDKQDGLTQEKFDPKTFFFLHDTNGDRYLDPLEMEALFYSEIKKVYGDDADGIEAHEDMARMREHSLNEMDEDRDGLVSLEEFMKYTSNKVFDTNDEWKPVRDHPEEVFTEQELEQFEKHYADYEADYDGIVSTNGDEPKGEEEHKGEGLKEKADHKDEVKELKAEKVEDHKVEVKAEEKKDENMVVPNEHAPAVGNAA

>Porifera_O.carmella_comp41841_c0_seq1.p1

MKTALYALLLSLLACFVFSVPVRPAKKAKSDTDNDDTAAHMDEYKRYVQQVLEVLGEHPDLREKIEAASEEEIANGELGKEISKLAPTYIRSQLDEAKRQTIQRQRLILRKRLEKQEGTRQRLDAHDEELFGGEVPEFDASDLKQVMLSRQEELKKEDKTRHDQFVKLEMDKEHQRRLALKKMGEKDRLEAEEKHKFVQEKKKDHPKLHHPGSEAQFKEVWEKEDGLDAQDFNPRTFFELHDTNGDGYLDTYELEALFINEVQKVYDEDEPEMQEEISRMREHVFGEVDKDQDGLIAYDEFKTSTELTSFSEDPGWQGLDEEDIYTDEEMEEYERMLQEEMESIDQESQNLIEKNKTPDEHGEKTVEVENAGEHHENVDVGEHREEVKTEE

>Porifera_T.wilhelma_c28025_g1_i1

QKGTILGWLVIIDVFGIVYSGSIKGGVMTVPSFLVTVAMRAVQVWICAVCLFTLVGLAAPFDAEAHRKQMDELRERMERLRDLPGLNREEYARYWEEAMKNDPRNFDRASKEFIKRGADLRGMNLDKLGANHPLVGDRLARGSDDIRSRLDEARRVNIEEMRQLQRLQALREDNLLKDGDWMAKREAMREKLPPRFTNEDIQARLKEQQRIMEELDQRRKDEFKRYEMQMEHVRRQKLKEMNEKQRLEAEEKYRQQQEEMKRHEKLKHPASKEQLEDVWANEDGFGKEAFDPKTFFHLHDKNGDKHLDPVELEALFYAEIRKAYPKQMEGMEAHEDMARMREHAMRELDTNKDGMVSLSEFMSYTGKEEFNEEEAWKPVVEEPLFSEEELKKFEDDYDYYYDYTYDDDGNIIDIKTRQRPARTNPPVDADPPKPDEENQSPPNEQEQPPAENLPHPVTPAIDDPKAVHINGVNPEQLMEKNANEEAPVDQEKQAEPNQL*

>Bilateria_C.intestinalis_F6U6Z4

MYRTICISCVVLLSFISLVNSAPAPTKRPQQPVSQNEAQAPPDANPVNTGLYYDQYLQKVIKLLESDESFKKKMETADLDDIKNGALSNELYSVSGEIRTKLDALKKEELNRLRKILHAKVELDKGRKVQRSGYLKQIANHLDHGSPHTFEADDLTKLIKTATSDLENFDRDRHEEFKKFEMRREMKREEKLKMLDEQERKKAEDDYRKHQQELADKSSIHHPGSKAQLEDVWNEEDGLGEQEFNPKTFFKMHDVNGDGYMDSMELEAIFDKDLSKVYADNSEESIMQMEEERTRMREHVMKEVDKNNDGLITLEEFLKYSDTPEFANPDENSYKTIDQMLLEHTLYTQDELRKYRENISDQEQEIKAKLEQAQLAAQQAQQAQAAQQAQAAQQAQAAQQAQAAQQAQAAQQAQAAQQAQAAQQAQAAQQAQAAQQAQAAQQAQAAQQAQ
