## Supplementary file 6 for "Pre-metazoan origin of neuropeptide signalling"

Homo sapiens – NUCB2

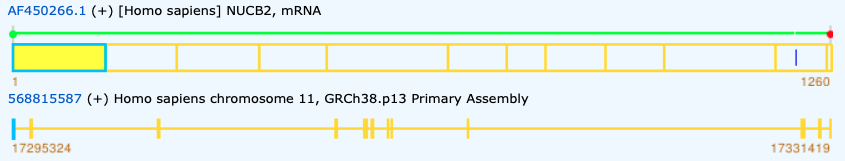

>AAM73810.1 [Homo sapiens] NUCB2 protein

MRWRTILLQYCFLLITCLLTALEAVPIDIDKTKVQNIHPVESAKIEPPDTGLYYDEYLKQVIDVLETDKHFREKLQKADIEEIKSGRLSKELDLVSHHVRTKLDELKRQEVGRLRMLIKAKLDSLQDIGMDHQALLKQFDHLNHLNPDKFESTDLDMLIKAATSDLEHYDKTRHEEFKKYEMMKEHERREYLKTLNEEKRKEEESKFEEMKKKHENHPKVNHPGSKDQLKEVWEETDGLDPNDFDPKTFFKLHDVNSDGFLDEQELEALFTKELEKVYDPKNEEDDMVEMEEERLRMREHVMNEVDTNKDRLVTLEEFLKATEKKEFLEPDSWETLDQQQFFTEEELKEYENIIALQENELKKKADELQKQKEELQRQHDQLEAQKLEYHQVIQQMEQKKLQGIPPSGPAGELKFEPHI

>AAM73810.1 [Homo sapiens] NUCB2 protein

MRWRTILLQYCFLLITCLLTALEAVPIDIDKTKVQNIHPVESAKIEPP--[0]--DTGLYYDEYLKQVIDVLETDKHFREKLQKADIEEIK--[0]--SGRLSKELDLVSHHVRTKLDELKRQEVGRLRMLIKAKLDSLQ--[1]--DIGMDHQALLKQFDHLNHLNPDKFESTDLDMLIKA--[0]--ATSDLEHYDKTRHEEFKKYEMMKEHERREYLKTLNEEKRKEEESKFEEMKKKHENHPKVNHP--[0]--GSKDQLKEVWEETDGLDPNDFDPKTFFKLH--[1]--DVNSDGFLDEQELEALFTKE--[0]--LEKVYDPKNEEDDMVEMEEERLRMREHVMNE--[0]--VDTNKDRLVTLEEFLKATEKKEFLEPDSWE--[0]--TLDQQQFFTEEELKEYENIIALQENELKKKADELQKQKEELQRQHDQLEAQKLEYHQ--[0]--VIQQMEQKKLQGIPPSGPAGELKFEP--[1]--HI

>AF450266.1 [Homo sapiens] NUCB2, mRNA

ATGAGGTGGAGGACCATCCTGCTACAGTATTGCTTTCTCTTGATTACATGTTTACTTACTGCTCTTGAAGCTGTGCCTATTGACATAGACAAGACAAAAGTACAAAATATTCACCCTGTGGAAAGTGCGAAGATAGAACCACCAGATACTGGACTTTATTATGATGAATATCTCAAGCAAGTGATTGATGTGCTGGAAACAGATAAACACTTCAGAGAAAAGCTCCAGAAAGCAGACATAGAGGAAATAAAGAGTGGGAGGCTAAGCAAAGAACTGGATTTAGTAAGTCACCATGTGAGGACAAAACTTGATGAACTGAAAAGGCAAGAAGTAGGAAGGTTAAGAATGTTAATTAAAGCTAAGTTGGATTCCCTTCAAGATATAGGCATGGACCACCAAGCTCTTCTAAAACAATTTGATCACCTAAACCACCTGAATCCTGACAAGTTTGAATCCACAGATTTAGATATGCTAATCAAAGCGGCAACAAGTGATCTGGAACACTATGACAAGACTCGTCATGAAGAATTTAAAAAATATGAAATGATGAAGGAACATGAAAGGAGAGAATATTTAAAAACATTGAATGAAGAAAAGAGAAAAGAAGAAGAGTCTAAATTTGAAGAAATGAAGAAAAAGCATGAAAATCACCCTAAAGTTAATCACCCAGGAAGCAAAGATCAACTAAAAGAGGTATGGGAAGAGACTGATGGATTGGATCCTAATGACTTTGACCCCAAGACATTTTTCAAATTACATGATGTCAATAGTGATGGATTCCTGGATGAACAAGAATTAGAAGCCCTATTTACTAAAGAGTTGGAGAAAGTATATGACCCTAAAAATGAAGAGGATGATATGGTAGAAATGGAAGAAGAAAGGCTTAGAATGAGGGAACATGTAATGAATGAGGTTGATACTAACAAAGACAGATTGGTGACTCTGGAGGAGTTTTTGAAAGCCACAGAAAAAAAAGAATTCTTGGAGCCAGATAGCTGGGAGACATTAGATCAGCAACAGTTCTTCACAGAGGAAGAACTAAAAGAATATGAAAATATTATTGCTTTACAAGAAAATGAACTTAAGAAGAAGGCAGATGAGCTTCAGAAACAAAAAGAAGAGCTACAACGTCAGCATGATCAACTGGAGGCTCAGAAGCTGGAATATCATCAGGTCATACAGCAGATGGAACAAAAAAAATTACAAGGAATTCCTCCATCAGGGCCAGCTGGAGAATTGAAGTTTGAGCCACACATTTAA

Homo sapiens – NUCB1

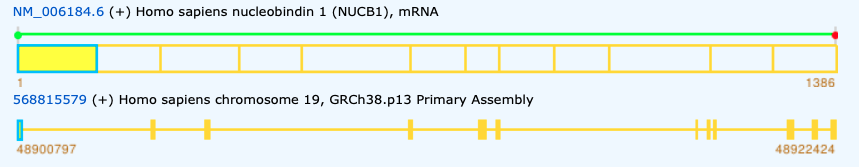

>NP_006175.2 nucleobindin-1 precursor [Homo sapiens]

MPPSGPRGTLLLLPLLLLLLLRAVLAVPLERGAPNKEETPATESPDTGLYYHRYLQEVIDVLETDGHFREKLQAANAEDIKSGKLSRELDFVSHHVRTKLDELKRQEVSRLRMLLKAKMDAEQDPNVQVDHLNLLKQFEHLDPQNQHTFEARDLELLIQTATRDLAQYDAAHHEEFKRYEMLKEHERRRYLESLGEEQRKEAERKLEEQQRRHREHPKVNVPGSQAQLKEVWEELDGLDPNRFNPKTFFILHDINSDGVLDEQELEALFTKELEKVYDPKNEEDDMREMEEERLRMREHVMKNVDTNQDRLVTLEEFLASTQRKEFGDTGEGWETVEMHPAYTEEELRRFEEELAAREAELNAKAQRLSQETEALGRSQGRLEAQKRELQQAVLHMEQRKQQQQQQQGHKAPAAHPEGQLKFHPDTDDVPVPAPAGDQKEVDTSEKKLLERLPEVEVPQHL

>NP_006175.2 nucleobindin-1 precursor [Homo sapiens]

MPPSGPRGTLLLLPLLLLLLLRAVLAVPLERGAPNKEETPATESP--[0]--DTGLYYHRYLQEVIDVLETDGHFREKLQAANAEDIK--[0]--SGKLSRELDFVSHHVRTKLDELKRQEVSRLRMLLKAKMDAEQDP--[1]--NVQVDHLNLLKQFEHLDPQNQHTFEARDLELLIQT--[0]--ATRDLAQYDAAHHEEFKRYEMLKEHERRRYLESLGEEQRKEAERKLEEQQRRHREHPKVNVP--[0]--GSQAQLKEVWEELDGLDPNRFNPKTFFILH--[1]--DINSDGVLDEQELEALFTKE--[0]--LEKVYDPKNEEDDMREMEEERLRMREHVMKN--[0]--VDTNQDRLVTLEEFLASTQRKEFGDTGEGWE--[0]--TVEMHPAYTEEELRRFEEELAAREAELNAKAQRLSQETEALGRSQGRLEAQKRELQQ--[0]--AVLHMEQRKQQQQQQQGHKAPAAHPEGQLKFHPDT--[1]--DDVPVPAPAGDQKEVDTSEKKLLERLPEVEVPQHL

>NM_006184.6 Homo sapiens nucleobindin 1 (NUCB1), mRNA

ATGCCTCCCTCTGGGCCCCGAGGAACCCTCCTTCTGTTGCCGCTGCTGCTGCTGCTCCTGCTTCGCGCCGTGCTGGCTGTCCCCCTGGAGCGAGGGGCGCCCAACAAGGAGGAGACCCCTGCGACTGAGAGTCCCGACACAGGCCTGTACTACCACCGGTACCTCCAGGAGGTCATCGATGTACTGGAGACGGATGGGCATTTCCGAGAGAAGCTGCAGGCTGCCAATGCGGAGGACATCAAGAGCGGGAAGCTGAGCCGAGAGCTGGACTTTGTCAGCCACCACGTCCGCACCAAGCTGGATGAGCTCAAGCGACAGGAGGTGTCACGGCTGCGGATGCTGCTCAAGGCCAAGATGGACGCCGAGCAGGATCCCAATGTACAGGTGGATCATCTGAATCTCCTGAAACAGTTTGAACACCTGGACCCTCAGAACCAGCATACATTCGAGGCCCGCGACCTGGAGCTGCTGATCCAGACGGCCACCCGGGACCTTGCCCAGTACGACGCAGCCCATCATGAAGAGTTCAAGCGCTACGAGATGCTTAAGGAACACGAGAGACGGCGTTATCTGGAGTCACTGGGAGAGGAGCAGAGAAAGGAGGCGGAGAGGAAGCTGGAAGAGCAACAGCGCCGGCACCGCGAGCACCCTAAAGTCAACGTGCCTGGCAGCCAAGCCCAGTTGAAGGAGGTGTGGGAGGAGCTGGATGGACTGGACCCCAACAGGTTTAACCCCAAGACCTTCTTCATACTGCATGATATCAACAGTGATGGTGTCCTGGATGAGCAGGAGCTGGAGGCACTCTTCACCAAGGAGCTGGAGAAAGTGTACGACCCAAAGAATGAGGAGGACGACATGCGGGAGATGGAGGAGGAGCGACTGCGCATGCGGGAGCATGTGATGAAGAATGTGGACACCAACCAGGACCGCCTCGTGACCCTGGAGGAGTTCCTCGCATCCACTCAGAGGAAGGAGTTTGGGGACACCGGGGAGGGCTGGGAGACAGTGGAGATGCACCCTGCCTACACCGAGGAAGAGCTGAGGCGCTTTGAAGAGGAGCTGGCTGCCCGGGAGGCAGAGCTGAATGCCAAGGCCCAGCGCCTCAGCCAGGAGACAGAGGCTCTAGGGCGGTCCCAGGGCCGCCTGGAGGCCCAGAAGAGAGAGCTGCAGCAGGCTGTGCTGCACATGGAGCAGCGGAAGCAGCAGCAGCAGCAGCAGCAAGGCCACAAGGCCCCGGCTGCCCACCCTGAGGGGCAGCTCAAGTTCCACCCAGACACAGACGATGTACCTGTCCCAGCTCCAGCCGGTGACCAGAAGGAGGTGGACACTTCAGAAAAGAAACTTCTCGAGCGGCTCCCTGAGGTTGAGGTGCCCCAGCATCTGTGA

Gallus gallus

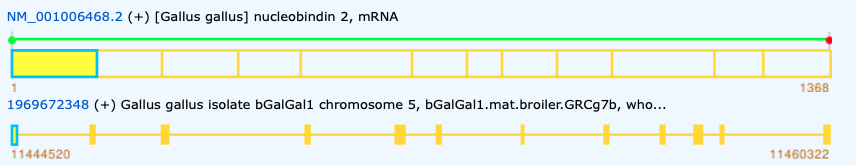

>NP_001006468.2 [Gallus gallus] nucleobindin-2 precursor

MKWQSLLPQQCILLIPCLLMALEAVPIDIDKTKVKGEGHVEGEKIENPDTGLYYDEYLRQVIDVLETDKHFREKLQTADIEEIKSGKLSRELDLVSHHVRTRLDELKRQEVARLRMLIKAKMDSVQDTGIDHQALLKQFEHLNHQNPDTFEPKDLDMLIKAATSDLENYDKTRHEEFKKYEMMKEHERREYLKTLDEEKRQREESKFEEMKKKHGDHPKVHHPGSKDQLKEVWEEADGLDPNEFDPKTFFKLHDVNNDRFLDEQELEALFTKELEKVYDPKNEEDDMVEMEEERLRMREHVMNEVDINKDRLVTLEEFLRATEKKEFLEPDSWETLDQQQLFTEDELKEFESHISQQEDELRKKAEELQKQKEELQRQHDQLQAQKQELQQVVKQMEQKKLQQANPPAGPAGELKFQPPGEHKIEEAPKHPAGGDQPLPPGHIQEPAARTDQVHP

>NP_001006468.2 [Gallus gallus] nucleobindin-2 precursor

MKWQSLLPQQCILLIPCLLMALEAVPIDIDKTKVKGEGHVEGEKIENP--[0]--DTGLYYDEYLRQVIDVLETDKHFREKLQTADIEEIK--[0]--SGKLSRELDLVSHHVRTRLDELKRQEVARLRMLIKAKMDSVQ--[1]--DTGIDHQALLKQFEHLNHQNPDTFEPKDLDMLIKA--[0]--ATSDLENYDKTRHEEFKKYEMMKEHERREYLKTLDEEKRQREESKFEEMKKKHGDHPKVHHP--[0]--GSKDQLKEVWEEADGLDPNEFDPKTFFKLH--[1]--DVNNDRFLDEQELEALFTKE--[0]--LEKVYDPKNEEDDMVEMEEERLRMREHVMNE--[0]--VDINKDRLVTLEEFLRATEKKEFLEPDSWE--[0]--TLDQQQLFTEDELKEFESHISQQEDELRKKAEELQKQKEELQRQHDQLQAQKQELQQ--[0]--VVKQMEQKKLQQANPPAGPAGELKFQP--[1]--PGEHKIEEAPKHPAGGDQPLPPGHIQEPAARTDQVHP

>NM_001006468.2 [Gallus gallus] nucleobindin 2, mRNA

ATGAAGTGGCAATCTCTCCTGCCTCAGCAATGTATTTTGCTGATACCATGTCTCCTAATGGCTTTGGAAGCTGTGCCTATAGACATAGATAAGACAAAAGTCAAAGGAGAAGGTCACGTAGAAGGAGAGAAGATAGAAAATCCGGATACAGGGCTTTATTATGATGAATATCTCAGGCAAGTAATTGATGTCTTGGAAACAGATAAGCATTTCAGGGAGAAACTCCAGACTGCTGACATAGAAGAAATAAAGAGTGGAAAACTAAGCAGAGAGCTTGACTTAGTAAGCCACCATGTGAGAACACGACTGGATGAACTCAAACGGCAAGAGGTGGCAAGATTAAGAATGTTAATTAAGGCCAAAATGGATTCTGTTCAAGATACTGGCATAGACCATCAGGCTCTCCTTAAACAGTTTGAGCACCTGAACCATCAGAATCCTGACACGTTTGAGCCTAAGGACTTAGATATGTTGATCAAAGCAGCTACAAGTGACCTGGAGAATTATGATAAAACCCGCCATGAGGAATTTAAGAAATACGAAATGATGAAAGAACACGAGAGGAGAGAGTATTTAAAAACATTGGATGAAGAAAAGAGACAGCGTGAAGAATCCAAATTTGAAGAGATGAAAAAGAAACATGGAGATCACCCTAAAGTTCACCATCCTGGAAGCAAAGATCAATTGAAAGAGGTGTGGGAAGAAGCAGATGGGCTAGATCCAAATGAGTTTGACCCCAAAACATTCTTCAAATTACATGATGTCAATAATGATCGTTTTCTGGATGAGCAGGAGTTGGAAGCTCTATTTACAAAAGAGCTAGAAAAAGTTTATGACCCGAAAAATGAAGAGGATGACATGGTTGAAATGGAAGAAGAGAGGCTGAGAATGAGAGAACATGTAATGAATGAGGTTGACATCAACAAGGACAGACTAGTAACTCTGGAAGAGTTCCTGCGAGCTACAGAAAAAAAAGAATTTTTGGAGCCAGATAGCTGGGAGACCCTAGATCAACAGCAGCTGTTCACTGAGGATGAACTGAAGGAATTTGAAAGTCATATTTCTCAGCAGGAAGATGAACTCAGAAAGAAGGCAGAAGAACTTCAGAAACAGAAGGAGGAGCTACAGAGGCAGCATGACCAGCTTCAGGCTCAGAAACAAGAGCTTCAGCAGGTTGTAAAACAGATGGAGCAAAAGAAACTACAGCAAGCAAATCCACCTGCAGGACCAGCTGGAGAACTGAAATTCCAGCCCCCTGGCGAGCACAAAATTGAAGAAGCACCGAAACATCCTGCAGGAGGTGATCAGCCTCTCCCACCAGGACATATCCAAGAACCAGCTGCTAGAACTGATCAAGTTCACCCCTAA

Branchiostoma belcheri

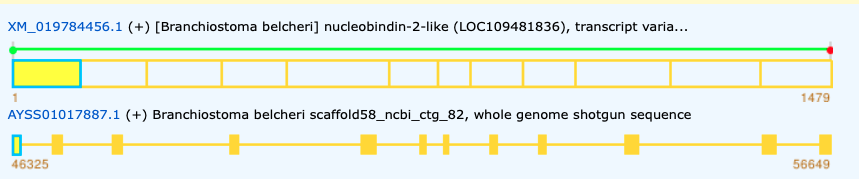

>XP_019640015.1 [Branchiostoma belcheri] nucleobindin-2-like isoform X1

MLTLRRCLLLAALVLLVSHDAMLAPVDPKKQQVEENETEKKEESQDTGLHYDSYLQEVVRVLETDPEFKKKLQEADIEDIKSGKLSAELNLVAHNVRSQLDELKRTEVTRLRKLQRVRMDMQNGRNGLHQGDLRDVKKMYEEFNHVDHSNPRSFEEEDLNKLIQAVVQDLENYDQKRHLEFKRYEMVKEHQRREKLSQLSEEDRKKEQERYEEMQRKHKDHPKMNHPGSKDQLEEVWEADGLAKEDFNPRAFFGMHDTNGDGYLDPMELEALFEKELEKVYKESNEEDDLREMEEERARMRKHVLREVDTNKDSMVSFEEFSDATKRKEFEQPEEDSFEPLDETDFFTEEELEEFEQQLAREADDLRQKEVELARQRAEHQQKRQQLQQVKMKQKQVVQMSQEQERQAEAQGQQVAVETNQESDQAAAGQDAVPAEVAQHAADGEGGQPAEMHEPVAAAVAADAVAVGGNLPPEGQLGEEQVQHQEAQQPAS

>XP_019640015.1 [Branchiostoma belcheri] nucleobindin-2-like isoform X1

MLTLRRCLLLAALVLLVSHDAMLAPVDPKKQQVEENETEKK--[0]--EESQDTGLHYDSYLQEVVRVLETDPEFKKKLQEADIEDIK--[0]--SGKLSAELNLVAHNVRSQLDELKRTEVTRLRKLQRVRMDMQNGRN--[1]--GLHQGDLRDVKKMYEEFNHVDHSNPRSFEEEDLNKLIQA--[0]--VVQDLENYDQKRHLEFKRYEMVKEHQRREKLSQLSEEDRKKEQERYEEMQRKHKDHPKMNHP--[0]--GSKDQLEEVWEADGLAKEDFNPRAFFGMH--[1]--DTNGDGYLDPMELEALFEKE--[0]--LEKVYKESNEEDDLREMEEERARMRKHVLRE--[0]--VDTNKDSMVSFEEFSDATKRKEFEQPEEDSFE--[0]--PLDETDFFTEEELEEFEQQLAREADDLRQKEVELARQRAEHQQKRQQLQQVKMKQKQ--[0]--VVQMSQEQERQAEAQGQQVAVETNQESDQAAAGQDAVPAEVAQHAADGEGGQPA--[1]--EMHEPVAAAVAADAVAVGGNLPPEGQLGEEQVQHQEAQQPAS

>XM_019784456.1 [Branchiostoma belcheri] nucleobindin-2-like (LOC109481836), transcript variant X1, mRNA

ATGTTGACCCTGAGACGATGTCTGCTGCTGGCAGCCCTGGTGCTGCTGGTATCCCATGATGCCATGCTTGCACCGGTCGACCCCAAGAAGCAGCAAGTGGAAGAGAATGAGACAGAGAAGAAGGAAGAGTCCCAGGACACAGGGCTACATTATGACAGCTACCTTCAGGAGGTGGTCAGAGTCCTGGAAACTGATCCTGAGTTCAAGAAAAAGCTTCAAGAGGCTGACATTGAGGACATCAAGTCTGGTAAACTGTCGGCTGAACTGAACCTGGTGGCGCACAACGTCCGGTCCCAGCTGGACGAACTGAAGAGAACGGAGGTCACCAGACTGCGCAAACTGCAGCGTGTACGCATGGACATGCAGAATGGCAGGAACGGATTACACCAGGGTGACTTGAGGGATGTGAAGAAAATGTACGAGGAGTTTAACCATGTGGACCACAGCAACCCTCGCTCATTCGAGGAGGAGGATCTCAACAAACTTATCCAGGCAGTGGTCCAGGACTTGGAGAACTACGACCAGAAGCGGCACCTGGAGTTTAAGAGGTACGAGATGGTGAAGGAACACCAGCGGAGAGAGAAGCTCAGCCAGCTGAGCGAGGAGGACAGGAAGAAGGAGCAGGAGAGATACGAGGAGATGCAGCGCAAACACAAGGACCACCCCAAGATGAACCATCCCGGCAGCAAAGATCAGCTGGAGGAAGTCTGGGAAGCCGATGGTCTTGCTAAGGAAGACTTCAACCCCAGGGCATTCTTTGGGATGCATGATACAAATGGCGATGGCTACCTGGATCCAATGGAGTTGGAGGCACTCTTTGAAAAAGAGTTGGAGAAGGTTTACAAGGAGAGTAACGAGGAAGATGACTTGCGAGAGATGGAGGAAGAGAGGGCTCGCATGAGGAAACACGTCTTACGAGAGGTGGACACCAACAAGGACTCCATGGTGAGCTTTGAGGAATTTTCAGATGCCACCAAAAGAAAAGAGTTTGAGCAGCCGGAAGAGGATAGTTTTGAGCCCCTGGATGAGACAGACTTCTTCACAGAAGAAGAACTTGAAGAGTTTGAGCAGCAGCTGGCGAGGGAAGCGGACGATCTGCGGCAGAAGGAGGTAGAACTGGCCCGGCAGCGCGCCGAGCACCAGCAGAAACGACAGCAACTTCAGCAGGTCAAAATGAAACAAAAACAGGTGGTTCAGATGAGTCAGGAGCAGGAGAGACAGGCGGAGGCTCAGGGTCAGCAGGTTGCCGTGGAGACTAACCAGGAGTCGGACCAGGCCGCGGCGGGACAGGACGCCGTGCCAGCCGAGGTGGCACAACATGCCGCAGATGGGGAGGGGGGGCAGCCCGCAGAGATGCATGAGCCTGTAGCTGCAGCTGTAGCAGCAGATGCCGTGGCTGTAGGTGGGAACCTGCCACCTGAGGGACAGCTGGGAGAGGAACAGGTGCAGCATCAGGAGGCACAGCAACCGGCATCATAA

Genomic: AYSS01017887.1

Priapulus caudatus

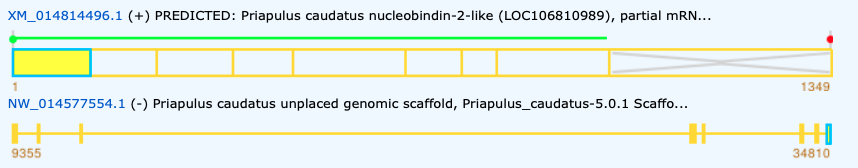

>Bilateria_P.caudatus_XP_014669982.1

MARLTWLILICLVASIVCPPVDKNSKKSEEDKKKENGTTDNNDIDLLEYDRYLKEIIGALEEDGDFKKKLESADFNDVQSGKIANELQFVNHNVRTKLDEIKRREMERLRQTTKKMLQKKEGRDVLKQGDIGGHIDHMNPRGFDMSDLAKLIQKTIADLETLDTQRKDNFKTYEMEKEYHHKEQVNNMTEEEKKIAEQEYLQQQEKHNDHPRVHHPGSKQQMQDVWTDNDHMDPGSFNPKTFFSMHDYNRDGYLDQKELEALFQTELDKLYDPDNPEDDMYERQEEMARMREHVQREVDTDSDGLISLAEFIAETHRSEWGEDEGWKDIRDEPVFDNEEYRAYEQFMAEQRHLASMDPQRLNTVPPHILEDFAKLPPDQQQQVMLQQHLQNQQQQQHPGQYQQHPGQVDPAQVDPRYQQHPGQVDQQYQQHHDQVDPRYQQHPDQVDQQ

>Bilateria_P.caudatus_XP_014669982.1

MARLTWLILICLVASIVCPPVDKNSKKSEEDKKKENGTTDNND--[0]--IDLLEYDRYLKEIIGALEEDGDFKKKLESADFNDVQ--[0]--SGKIANELQFVNHNVRTKLDEIKRREMERLRQTTKKMLQKKE--[1]--GRDVLKQGDIGGHIDHMNPRGFDMSDLAKLIQK--[0]--TIADLETLDTQRKDNFKTYEMEKEYHHKEQVNNMTEEEKKIAEQEYLQQQEKHNDHPRVHHP--[0]--GSKQQMQDVWTDNDHMDPGSFNPKTFFSMH--[1]--DYNRDGYLDQKELEALFQTE--[0]--LDKLYDPDNPEDDMYERQEEMARMREHVQREVDTDSDGLISLAEFIAETHRSEWGEDEGWKDIRDEPVFDNEEYRAYEQFMAEQRHLASMDPQRLNTVPPHILEDFAKLPPDQQQQVMLQQHLQNQQQQQHPGQYQQHPGQVDPAQVDPRYQQHPGQVDQQYQQHHDQVDPRYQQHPDQVDQQ

>XM_014814496.1 PREDICTED: Priapulus caudatus nucleobindin-2-like (LOC106810989), partial mRNA

ATGGCCAGACTTACCTGGCTCATACTTATTTGCCTCGTGGCAAGCATAGTTTGCCCACCGGTCGACAAAAACTCGAAAAAATCCGAAGAAGACAAAAAAAAGGAAAACGGAACTACTGATAACAACGACATCGATCTGCTCGAATATGATCGTTACCTGAAGGAGATTATCGGTGCACTGGAGGAGGATGGCGATTTCAAGAAAAAACTTGAGAGTGCTGACTTTAATGATGTCCAGTCTGGCAAGATCGCGAACGAACTCCAGTTTGTCAACCACAACGTCCGAACAAAGCTGGACGAGATCAAGCGCAGAGAGATGGAGCGACTTCGCCAGACGACGAAGAAGATGCTTCAGAAGAAAGAAGGCAGGGATGTGTTGAAGCAAGGAGACATCGGCGGCCACATTGACCACATGAACCCGAGAGGCTTCGACATGTCTGACTTGGCAAAGCTCATACAGAAGACGATCGCCGACCTTGAGACGCTCGACACGCAGCGCAAGGACAACTTCAAGACGTACGAGATGGAGAAGGAGTACCATCACAAGGAGCAGGTGAACAACATGACAGAGGAGGAGAAGAAGATCGCCGAGCAGGAGTACCTTCAGCAGCAGGAGAAGCACAACGACCACCCGCGCGTGCATCACCCGGGAAGCAAACAGCAGATGCAGGACGTGTGGACAGACAATGATCACATGGACCCAGGCTCTTTTAATCCGAAAACTTTCTTCAGTATGCACGATTACAACAGAGATGGATATTTGGATCAGAAGGAGCTAGAAGCCCTATTCCAGACCGAGCTGGACAAGCTGTACGACCCAGACAACCCGGAGGACGACATGTACGAGAGGCAGGAGGAGATGGCGCGCATGCGCGAGCACGTGCAGCGTGAAGTCGACACCGACAGCGACGGCCTCATCAGCCTCGCCGAGTTCATCGCCGAGACACACCGCAGCGAGTGGGGCGAGGACGAGGGCTGGAAGGACATACGCGACGAGCCGGTGTTCGATAACGAGGAGTACCGCGCCTACGAGCAGTTCATGGCCGAGCAGCGCCACCTAGCGAGCATGGACCCGCAACGCCTCAACACGGTTCCGCCACATATACTTGAGGACTTTGCCAAATTGCCTCCGGACCAGCAGCAGCAGGTTATGCTGCAGCAGCATCTTCAGAATCAGCAGCAGCAGCAGCATCCAGGGCAATACCAGCAACATCCCGGCCAAGTTGACCCGGCTCAAGTCGACCCACGTTACCAGCAACATCCTGGTCAAGTCGACCAACAGTACCAGCAACATCACGACCAAGTTGACCCTCGTTACCAGCAACATCCCGACCAAGTTGACCAACAATT...

Genomic: NW_014577554.1 (C-terminal part missing)

Centruroides sculpturatus

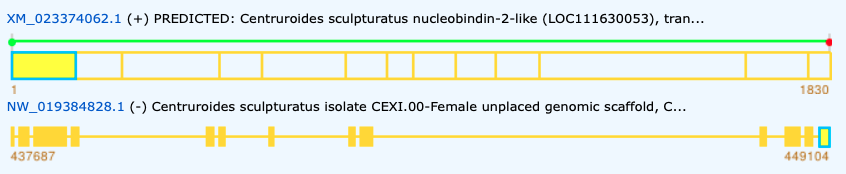

>Bilateria_C.sculpturatus_XP_023229830

MNRSQFVFLLIFALLAIHLVGAPPVDPKKKKEKEKEHEEDEGNELDDYGLEYGRYLQQVVQALEEDKDFAKKLENISTDQIKSGDVAKELEFVKHNVRTKLDELKRIELERLRKLSREEMERLELGLYRNEDGHLAPPSIDGRRWRTISSAGDGEGINRKHIKVPHHLDVDNKHTFEIEDLRKLMKAATRDLEELDRRRREEFKQYELEKEYQFRESLKNMSESEKAEQMKKHEEMLKKHKDHPKVHHPGSKQQLEQVWEEQDHMQGNEFDPRVFFAMHDVNGDGYLDEQEVEAILSLEIKKMYDPQNNVEDDPQEMMEEYNRMREHVYKETDKNKDGLISKEEFMDMTKRADFEKDDGWRGLDEQQIYTEEELKAYEKKRIEELQKLNAYYGYQNAHGGAPVPHGMPQYQGVPPPHPGYQPHPQQMPQGYQPHPQQMPQGAHQQGYQSHHQQAQMGGAHPQGYQSQQQGEAHHQPSQQQQGYQPQMQQMPQPGAHQIPQPGYPPHQQQMPQGIHQPMPGMQQQPHPGAQYGIPNNQNPQMRHPAAQGIHPGVQIVQGNAHGLPTHAPSDPPPRQPQPTQYNPQQRQPAVEQKQAEDTNANTANQKQGN

>Bilateria_C.sculpturatus_XP_023229830

MNRSQFVFLLIFALLAIHLVGAPPVDPKKKKEKEKEHEEDEGNELDDY--[0]--GLEYGRYLQQVVQALEEDKDFAKKLENISTDQIK--[0]--SGDVAKELEFVKHNVRTKLDELKRIELERLRKLSREEMERLELGLYRNEDGHLAPPSIDGRRWRTISSAGDGE--[1]--GINRKHIKVPHHLDVDNKHTFEIEDLRKLMKA--[0]--ATRDLEELDRRRREEFKQYELEKEYQFRESLKNMSESEKAEQMKKHEEMLKKHKDHPKVHHP--[0]--GSKQQLEQVWEEQDHMQGNEFDPRVFFAMH--[1]--DVNGDGYLDEQEVEAILSLE--[0]--IKKMYDPQNNVEDDPQEMMEEYNRMREHVYKE--[0]--TDKNKDGLISKEEFMDMTKRADFEKDDGWR--[0]--GLDEQQIYTEEELKAYEKKRIEELQKLNAYYG--[0]--YQNAHGGAPVPHGMPQYQGVPPPHPGYQPHPQQMPQGYQPHPQQMPQGAHQQGYQSHHQQAQMGGAHPQGYQSQQQGEAHHQPSQQQQGYQPQMQQMPQPGAHQIPQPGYPPHQQQMPQGIHQPMPGMQQQPHPGAQYGIPNNQNPQMRHPAAQ--[0]--GIHPGVQIVQGNAHGLPTHAPSDPPPRQPQPTQYNPQQRQPAVEQK--[0]--QAEDTNANTANQKQGN

>XM_023374062.1 PREDICTED: Centruroides sculpturatus nucleobindin-2-like (LOC111630053), transcript variant X2, mRNA

ATGAACAGGTCACAATTTGTTTTTCTACTTATTTTCGCACTTTTGGCCATTCATCTGGTCGGCGCACCACCCGTAGACCCTAAGAAAAAGAAAGAAAAAGAAAAGGAGCATGAGGAAGACGAAGGAAATGAATTGGACGATTATGGGTTAGAATACGGACGTTACCTCCAGCAAGTCGTTCAGGCTCTGGAGGAGGATAAAGACTTTGCTAAGAAGTTGGAAAATATTTCCACCGACCAAATTAAAAGTGGAGACGTGGCCAAGGAATTGGAGTTTGTCAAACACAATGTTCGCACGAAGCTAGACGAGTTGAAACGTATAGAACTAGAACGTCTGCGTAAGCTAAGTAGAGAGGAGATGGAGAGATTGGAATTAGGTCTGTACAGAAACGAAGATGGTCACTTAGCTCCGCCGTCCATCGACGGTCGTAGGTGGCGGACTATATCCTCGGCGGGTGACGGTGAAGGAATAAATCGAAAACATATAAAAGTGCCTCATCATTTGGACGTAGATAACAAGCATACATTTGAAATTGAAGATCTGAGAAAATTAATGAAAGCGGCCACGCGCGATCTGGAAGAATTAGACAGGAGACGGAGAGAAGAATTCAAACAGTACGAGTTAGAAAAGGAATATCAATTCCGGGAGAGCCTGAAAAATATGTCCGAGAGCGAAAAGGCCGAACAAATGAAGAAACACGAAGAAATGCTCAAGAAACACAAGGATCATCCCAAAGTCCATCATCCCGGAAGTAAGCAGCAATTGGAACAAGTGTGGGAAGAGCAGGATCACATGCAGGGCAACGAATTTGATCCAAGAGTCTTCTTTGCCATGCACGATGTTAACGGTGATGGTTATTTGGACGAACAAGAAGTAGAAGCTATCCTGAGTCTAGAGATCAAGAAAATGTACGACCCTCAGAACAACGTGGAGGACGATCCACAGGAGATGATGGAAGAATACAATAGAATGAGAGAGCACGTCTACAAAGAGACCGATAAAAACAAAGACGGACTCATCAGCAAGGAGGAGTTCATGGACATGACAAAGAGGGCCGATTTCGAGAAGGACGACGGGTGGAGGGGATTAGATGAACAACAAATTTATACCGAAGAAGAATTAAAAGCCTACGAAAAGAAAAGAATAGAAGAGTTACAGAAGTTGAATGCTTATTATGGATATCAGAATGCTCATGGTGGTGCTCCAGTGCCTCATGGAATGCCGCAGTATCAGGGCGTTCCTCCACCTCATCCAGGATATCAACCTCATCCACAGCAGATGCCACAAGGTTATCAGCCTCACCCGCAACAAATGCCGCAAGGTGCGCACCAGCAGGGATACCAGTCTCATCATCAACAGGCGCAGATGGGCGGAGCTCATCCGCAGGGCTATCAGTCTCAACAACAGGGAGAGGCGCACCACCAACCGTCACAGCAGCAGCAGGGTTATCAACCTCAGATGCAACAGATGCCACAGCCCGGAGCACATCAAATACCACAGCCCGGCTATCCTCCTCACCAGCAGCAGATGCCACAAGGTATTCATCAGCCGATGCCCGGAATGCAACAACAACCCCATCCCGGTGCCCAGTACGGAATACCGAACAATCAGAATCCTCAAATGCGCCACCCCGCAGCTCAGGGAATACATCCAGGAGTCCAAATAGTACAAGGAAATGCTCACGGTCTGCCCACTCACGCCCCAAGCGACCCGCCCCCACGACAACCCCAGCCCACTCAATATAATCCACAACAGCGGCAGCCTGCCGTAGAACAAAAACAGGCAGAAGATACCAACGCAAACACGGCAAACCAGAAACAAGGAAACTAG

Genomic: NW_019384828.1

Octopus bimaculoides

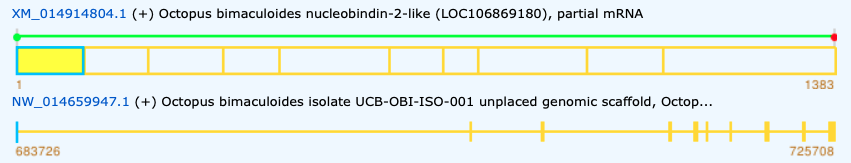

>Bilateria_O.bimaculoides_A0A0L8HQS8

MFITILVSLVLFVINGIVCPPVKPGTKPKEGASVNDTNETGLAYDRYLNEIVTVLEEDPAFKKKLETSDLQYIKSGAISKDLDLVHHGIRQQLDEIKRREVIRLKSLVQGQFKYKSEERGNRKVALHHININNPDSFEQDDLAKLIKQATHDLDELDRERKEEFKQYEMKKEHEIKQKLANMNETERKIEVQHIKEMKEKHKQHPKIHHPGSKAQLEEVWEEVDRLNGENFNPKTFFKLHDVNGDGVLEPSEVEALFTREIEKMYDPNNPEDDMKEAKEEINRMREHVFKEADTDDNHLLDEGEFERLTQREEFGKDEGWKGLDEDDEIYSEEEMQQYEKMLREEEERLKKQQIYDNLAQPGQVVQPVDPNVVHMQQPHQEQQQQHQEQQNLQYAHQEKLRQQQLTDAQHQQQFAQQQAALHQQQQQQPVGHQQQQQPVGHQQQQPVVHQQQQPVVHQQQQPVVQQHQQQQPVVQQHQQQPVVQQQQQQHQPVVQQHQ

>Bilateria_O.bimaculoides_A0A0L8HQS8

MFITILVSLVLFVINGIVCPPVKPGTKPKEGASVNDTN--[0]--ETGLAYDRYLNEIVTVLEEDPAFKKKLETSDLQYIK--[0]--SGAISKDLDLVHHGIRQQLDEIKRREVIRLKSLVQGQFKYKS--[1]--EERGNRKVALHHININNPDSFEQDDLAKLIKQ--[0]--ATHDLDELDRERKEEFKQYEMKKEHEIKQKLANMNETERKIEVQHIKEMKEKHKQHPKIHHP--[0]--GSKAQLEEVWEEVDRLNGENFNPKTFFKLH--[1]--DVNGDGVLEPSEVEALFTRE--[0]--IEKMYDPNNPEDDMKEAKEEINRMREHVFKEADTDDNHLLDEGEFERLTQREEFGKDEGWK--[0]--GLDEDDEIYSEEEMQQYEKMLREEEERLKKQQIYDNLAQPGQV--[0]--VQPVDPNVVHMQQPHQEQQQQHQEQQNLQYAHQEKLRQQQLTDAQHQQQFAQQQAALHQQQQQQPVGHQQQQQPVGHQQQQPVVHQQQQPVVHQQQQPVVQQHQQQQPVVQQHQQQPVVQQQQQQHQPVVQQHQ

>XM_014914804.1 Octopus bimaculoides nucleobindin-2-like (LOC106869180), partial mRNA

ATGTTTATAACAATATTAGTAAGTTTGGTGCTTTTTGTCATCAATGGTATAGTATGTCCACCAGTAAAACCGGGCACCAAACCAAAAGAAGGTGCATCAGTCAATGATACCAATGAAACAGGACTTGCTTATGACCGTTACTTAAATGAAATTGTTACTGTTTTGGAGGAAGATCCTGCCTTTAAGAAAAAATTAGAAACTTCTGATCTTCAGTATATAAAGAGTGGTGCAATATCAAAAGACTTAGACTTGGTGCATCATGGCATTCGACAACAATTGGATGAGATTAAGCGAAGAGAAGTAATTCGTCTGAAATCTCTTGTCCAAGGGCAGTTTAAATATAAATCTGAAGAAAGAGGTAATAGAAAAGTGGCTTTGCATCATATCAATATAAATAATCCAGATTCATTTGAACAAGATGACTTGGCAAAACTCATCAAACAGGCTACTCATGATTTGGATGAATTGGATCGAGAGAGGAAAGAAGAGTTTAAACAATATGAGATGAAGAAGGAGCATGAAATAAAACAAAAATTAGCTAATATGAATGAAACTGAACGAAAAATTGAAGTTCAACATATAAAAGAAATGAAAGAAAAACATAAGCAACATCCCAAAATACATCATCCGGGTAGCAAAGCCCAGCTGGAGGAAGTCTGGGAAGAAGTTGACCGTCTAAATGGTGAAAACTTTAATCCCAAGACCTTTTTCAAGCTACATGATGTCAATGGTGATGGTGTCCTGGAGCCAAGTGAAGTTGAAGCTCTTTTTACAAGAGAGATTGAAAAGATGTATGATCCAAATAATCCTGAGGATGATATGAAAGAAGCCAAAGAAGAAATTAATCGGATGAGGGAACATGTCTTCAAAGAAGCTGATACTGATGATAACCACCTTTTGGATGAAGGAGAGTTTGAACGTCTTACTCAACGAGAAGAGTTTGGGAAAGATGAAGGATGGAAGGGCTTGGATGAAGATGATGAAATATATTCAGAAGAAGAAATGCAACAATATGAGAAGATGTTACGTGAAGAAGAAGAGCGCCTTAAAAAGCAACAAATCTATGACAATTTGGCTCAACCGGGACAAGTGGTACAGCCTGTTGATCCAAACGTTGTTCATATGCAGCAACCACATCAAGAACAACAACAGCAACATCAGGAGCAACAAAATCTGCAATATGCACATCAAGAAAAACTCAGACAACAGCAATTGACTGATGCTCAACATCAACAACAGTTTGCTCAACAGCAAGCTGCTTTACATCAACAGCAGCAACAGCAGCCTGTAGGTCATCAGCAGCAACAGCAGCCTGTAGGTCATCAGCAACAACAGCCTGTAGTTCATCAGCAACAACAGCCTGTAGTTCATCAGCAACAACAG...

Genomic: NW_014659947.1

Nematostella vectensis

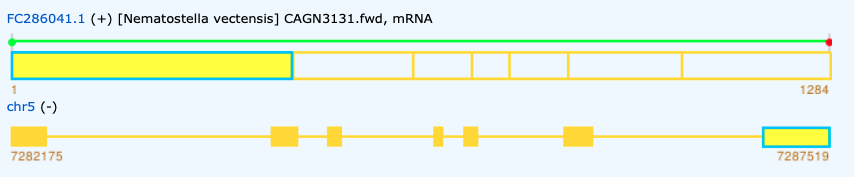

>FC286041.1 [Nematostella vectensis] CAGN3131.fwd

MDSKLSLLSLLVVLVLVTLCYAPPVKRGKPAEKAPEPNKDDPEYARYLRQVIEILEKDEDYVRKLMNASDDDLRSGRIAEDIDLVKHDVRSKLDELKRQEVERQRMIRRQMNDHLNGIKEREYWNPLFDDENPDFFGADDFKKLLWKHHEEMDKQDRERREEFKKHEMQKEHERKIKMKDMDEKHRKEAEEHFRELQEKHHNQSKQLHHPGSKAQLEQVWEESDGLDAKDFDPKTFFKLHDVNGDGFLDTGELEALFVKEVTKLYNPKDEDYDPKERDEEISRMREHVMNEIDKDKDGFVSQDEFLQSSKGEEFDKDDGWKSVEDERPYTDEELAEFEKSLEQDQHPSEQETDHEQPAQHDQQQQNTGQQQQHDKQQQHDQQQQHDQQQQHDQQQQQRHDQQQQHHDQQQQQHHDQQQQHHEQQKPQ

>FC286041.1 [Nematostella vectensis] CAGN3131.fwd

MDSKLSLLSLLVVLVLVTLCYAPPVKRGKPAEKAPEPNKDDPEYARYLRQVIEILEKDEDYVRKLMNASDDDLRSGRIAEDIDLVKHDVRSKLDELKRQEVERQRMIRRQMNDHLNGIKEREYWNPLFDDENPDFFGADDFKKLLWK--[0]--HHEEMDKQDRERREEFKKHEMQKEHERKIKMKDMDEKHRKEAEEHFRELQEKHHNQSKQLHHP--[0]--GSKAQLEQVWEESDGLDAKDFDPKTFFKLH--[1]--DVNGDGFLDTGELEALFVKE--[0]--VTKLYNPKDEDYDPKERDEEISRMREHVMNE--[0]--IDKDKDGFVSQDEFLQSSKGEEFDKDDGWKSVEDERPYTDEELAEFEKSLEQDQHPSEQ--[0]--ETDHEQPAQHDQQQQNTGQQQQHDKQQQHDQQQQHDQQQQHDQQQQQRHDQQQQHHDQQQQQHHDQQQQHHEQQKPQ

>FC286041.1 [Nematostella vectensis] CAGN3131.fwd, mRNA

ATGGACTCGAAATTATCACTGCTTAGTTTACTAGTTGTATTGGTGCTTGTCACATTGTGTTATGCGCCTCCAGTCAAACGTGGAAAACCCGCTGAAAAAGCACCTGAACCAAACAAGGACGATCCTGAATATGCTAGATATCTTAGGCAAGTAATAGAGATTTTAGAAAAGGACGAAGACTATGTTCGTAAGCTTATGAACGCTTCTGATGATGATCTTCGATCAGGCCGAATCGCAGAGGACATTGATTTAGTCAAACATGATGTGCGTTCTAAACTCGATGAGTTGAAACGACAAGAAGTAGAAAGACAACGTATGATTCGAAGGCAAATGAACGATCATCTTAATGGTATCAAAGAACGAGAATACTGGAATCCTCTATTCGACGATGAAAATCCAGATTTCTTTGGCGCGGATGATTTCAAGAAACTTTTATGGAAGCATCATGAAGAGATGGACAAACAAGATCGTGAGAGAAGAGAGGAATTTAAAAAACATGAAATGCAAAAAGAACACGAAAGGAAAATTAAAATGAAAGACATGGACGAGAAACATAGGAAGGAAGCAGAAGAGCACTTCAGGGAGCTGCAAGAGAAGCATCACAACCAATCCAAGCAGCTACACCATCCAGGGAGCAAAGCACAATTGGAACAAGTCTGGGAAGAGTCTGATGGTTTGGACGCCAAAGACTTTGATCCAAAAACATTCTTTAAGCTTCATGATGTTAATGGGGATGGCTTCTTGGATACAGGAGAGCTAGAGGCATTGTTTGTTAAAGAGGTTACCAAGCTGTACAACCCCAAAGATGAAGACTATGACCCCAAGGAAAGAGATGAAGAAATCTCCCGGATGAGAGAACATGTCATGAATGAGATTGACAAGGATAAAGATGGCTTTGTGTCACAAGATGAGTTCCTCCAATCAAGCAAAGGAGAAGAGTTTGATAAAGATGATGGCTGGAAGAGTGTGGAAGACGAGCGGCCGTACACAGACGAGGAACTAGCTGAGTTTGAGAAGAGTCTAGAGCAGGACCAACACCCTTCAGAACAGGAAACTGACCATGAACAACCTGCACAACATGATCAGCAACAGCAGAACACAGGGCAGCAACAACAGCATGATAAACAACAACAACATGATCAACAACAACAACATGATCAACAACAACAACATGATCAACAACAGCAACAACGCCATGATCAACAACAACAACACCATGATCAACAACAACAACAGCACCATGATCAACAACAACAACACCATGAGCAGCAAAAGCCGCAATAA

Genomic: New genome, chromosome 5

Hydra vulgaris

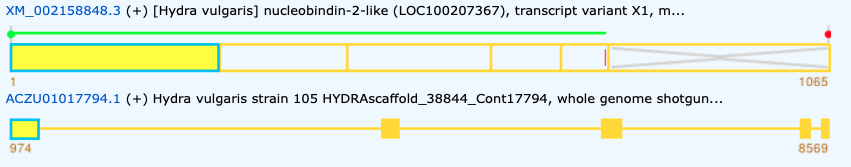

>XP_002158884.2 [Hydra vulgaris] nucleobindin-2-like, isoform X1

MKELLTFGIFLALISLHVAPPVGKKEPELPANETGTVEDKQDAEYFRYLSQVVEVLEKDPEFKSKLHNASEEDIRSGKIANYLDLVGHGVRLKLDEIKRTEVEYQRELLRQRQDFMSGIERNYWNPIHHDNKDSFEMEDLKKLLSKHNDMMSAQDAKRHEEFKAYEMEKEHERREKLHNMTAEERAKEEALYKEHREQRSKHEKIHEPGHKAQLEETWEKEDGLDPESFDPRTFFNLHDKNSDHYLDLYELETIFLADIDKVYNESNPEVDLRERSEEIERMREHVMKNMDKDKDGLISFTEFMDETKSEDFEKDEDWKPLTEQDQFTEEELQEYEKMLSENPHGENVKQEASH

>XP_002158884.2 [Hydra vulgaris] nucleobindin-2-like, isoform X1

MKELLTFGIFLALISLHVAPPVGKKEPELPANETGTVEDKQDAEYFRYLSQVVEVLEKDPEFKSKLHNASEEDIRSGKIANYLDLVGHGV--[2]--RLKLDEIKRTEVEYQRELLRQRQDFMSGIERNYWNPIHHDNKDSFEMEDLKKLLSK--[0]--HNDMMSAQDAKRHEEFKAYEMEKEHERREKLHNMTAEERAKEEALYKEHREQRSKHEKIHEP--[0]--GHKAQLEETWEKEDGLDPESFDPRTFFNLH--[1]--DKNSDHYLDLYELETIFLADI--[0]--DKVYNESNPEVDLRERSEEIERMREHVMKNMDKDKDGLISFTEFMDETKSEDFEKDEDWKPLTEQDQFTEEELQEYEKMLSENPHGENVKQEASH

>XM_002158848.3 [Hydra vulgaris] nucleobindin-2-like (LOC100207367), transcript variant X1, mRNA

ATGAAGGAACTTCTTACTTTTGGTATTTTTTTGGCACTAATTAGTTTGCATGTTGCACCGCCTGTTGGTAAAAAAGAACCAGAACTACCAGCAAATGAGACTGGAACTGTAGAAGACAAGCAAGATGCTGAATACTTTAGATATTTATCACAAGTTGTAGAAGTACTTGAAAAAGATCCTGAATTTAAATCCAAGCTACATAATGCTTCAGAAGAAGATATTCGTTCAGGAAAAATTGCCAATTATTTAGATTTAGTTGGGCATGGAGTGAGATTAAAACTTGATGAGATAAAAAGAACAGAAGTTGAATATCAAAGAGAGTTGTTACGACAAAGACAAGACTTCATGTCTGGCATAGAAAGAAACTATTGGAACCCTATACATCATGACAATAAAGACTCTTTTGAAATGGAGGATTTGAAAAAACTATTATCAAAACACAATGATATGATGAGTGCACAAGATGCCAAACGTCATGAAGAATTTAAAGCTTATGAGATGGAAAAAGAACATGAACGCAGGGAAAAATTACATAATATGACTGCTGAAGAGCGTGCTAAGGAAGAAGCATTATACAAAGAACATCGTGAGCAGAGAAGTAAACATGAGAAGATACATGAACCCGGACATAAAGCTCAGTTGGAAGAAACATGGGAAAAAGAAGATGGTTTAGATCCTGAAAGCTTTGATCCAAGAACTTTTTTTAACTTGCATGATAAAAATTCTGACCATTACTTGGATCTTTACGAGTTGGAGACCATATTTTTGGCTGATATAGATAAAGTCTACAATGAAAGTAATCCTGAAGTTGATCTTAGAGAAAGGTCTGAAGAAATAGAAAGAATGAGAGAACATGTCATGAAGAATATGGATAAAGATAAAGATGGATTAATATCATTTACGGAGTTTATGGATGAAACCAAATCTGAAGACTTCGAAAAGGATGAAGATTGGAAACCTTTAACTGAGCAAGATCAATTTACTGAAGAAGAACTTCAAGAATACGAAAAAATGCTTTCAGAAAACCCTCATGGAGAAAATGTGAAACAGGAAGCCTCGCACTAA

Genomic: ACZU01017794.1 (last part was not aligned)

Mnemiopsis leidyi

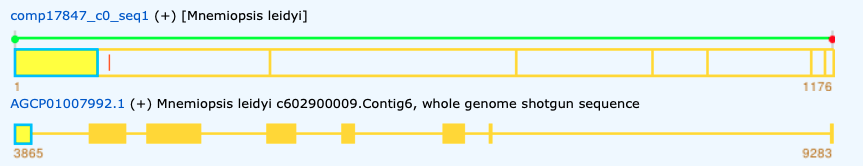

>Ctenophora_M.leidyi_comp17847_c0_seq1_.p1

MKLLIASFLLLFLAAEGAENPESVIPTKTEYERYLDQILLVFEKDPKVKEILKNATPQDIKNGVVEKTLRRLHPDIRSKLDEVKRNIIMEYRQKLRRAKMDDKVPDDLYKFQDEDLKHLLKERHDRLAQMNDERKRQFKKYEMEKELKHRMALRAMNEYDRKVQEALDEEQKQKKMEKVKNLKHPGNKAQLEEVWENEGFDKGTFNPKTFFKMHDLNNDGELDDFELEAIFEKEVEDLFESFEDADEEEMAEEVAQMREHVINHVDKDKDGVIQLKEFIEYANGSEFDHEDPWDAAFNAEFTDQDLDEYEKTLDDIEREDVKAQAEIEKQKRLETIKKYEEESKINMEMLKAHSEKLKEALMAANKAKDLEKAKKLVAEAQADQEANKASP

>Ctenophora_M.leidyi_comp17847_c0_seq1_.p1

MKLLIASFLLLFLAAEGAENPESVIPTKTEYERYLDQILL--[0]--VFEKDPKVKEILKNATPQDIKNGVVEKTLRRLHPDIRSKLDEVKRNIIMEYRQKLRRAKMDDKVPDDLYKFQDEDLKHLLKE--[0]--RHDRLAQMNDERKRQFKKYEMEKELKHRMALRAMNEYDRKVQEALDEEQKQKKMEKVKNLKHPGNKAQLEEVWENEGFDKGTFNPKTFFKMHDLNNDGELDDFELEAIFEKEVEDLFE--[0]--SFEDADEEEMAEEVAQMREHVINHVDKDKDGVIQLKEFIEYANGSEFDHEDPWDAAFNAEFTDQD--[0]--LDEYEKTLDDIEREDVKAQAEIEKQK--[2]--RLETIKKYEEESKINMEMLKAHSEKLKEALMAANKAKDLEKAKKLVAEAQ--[0]--ADQEANK--[0]--ASP

>comp17847_c0_seq1 [Mnemiopsis leidyi]

ATGAAGCTGTTAATTGCAAGCTTCCTTTTATTATTTCTGGCAGCTGAAGGGGCAGAGAATCCTGAAAGTGTGATACCAACCAAAACTGAGTATGAGAGATATTTGGATCAGATTCTTCTGGTATTCGAAAAAGATCCAAAAGTGAAAGAAATCCTGAAGAATGCAACACCTCAAGACATCAAGAATGGTGTAGTAGAAAAAACTTTACGCAGATTGCATCCAGACATTCGAAGCAAACTAGATGAAGTTAAACGGAACATAATAATGGAGTATAGACAGAAGTTGCGGAGAGCCAAAATGGATGACAAGGTCCCTGATGACCTGTACAAGTTCCAGGATGAGGATTTGAAACATCTTCTTAAGGAGCGCCATGACAGACTGGCTCAAATGAATGACGAAAGAAAGAGGCAATTTAAAAAATATGAAATGGAAAAAGAGTTGAAACACAGAATGGCTCTGCGGGCTATGAACGAATATGATAGAAAAGTACAAGAAGCTCTCGACGAAGAGCAGAAGCAAAAGAAGATGGAAAAAGTGAAGAATCTCAAGCATCCTGGTAATAAGGCTCAGTTAGAAGAAGTGTGGGAGAATGAAGGTTTCGACAAGGGTACCTTTAACCCCAAAACATTTTTCAAAATGCATGATTTGAACAACGATGGTGAATTGGACGACTTTGAACTTGAAGCAATTTTTGAAAAAGAGGTTGAAGACCTCTTTGAGTCATTCGAGGATGCTGATGAAGAAGAAATGGCCGAGGAGGTAGCTCAAATGCGTGAACATGTCATTAATCATGTAGATAAAGATAAAGATGGGGTAATTCAGTTGAAAGAGTTTATTGAATATGCTAACGGAAGTGAGTTCGATCATGAGGACCCCTGGGATGCTGCTTTCAATGCAGAGTTTACCGATCAAGATTTAGACGAATATGAGAAAACTCTTGATGACATAGAAAGAGAAGATGTAAAAGCTCAAGCGGAGATCGAAAAACAAAAAAGACTGGAAACTATCAAGAAATACGAGGAAGAAAGCAAGATAAATATGGAAATGCTAAAAGCACATTCTGAAAAACTGAAAGAGGCTCTCATGGCTGCAAATAAGGCCAAGGATCTTGAAAAGGCCAAAAAATTAGTCGCTGAGGCTCAAGCAGATCAAGAAGCCAATAAAGCATCACCGTGA

Genomic: AGCP01007992.1

Amphimedon queenslandica

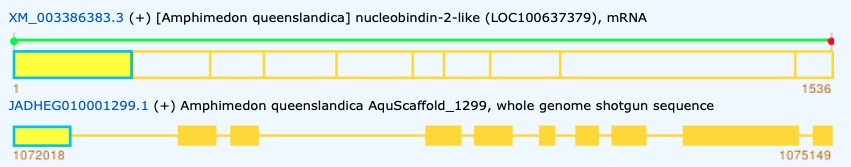

>XP_003386431.1 [Amphimedon queenslandica] nucleobindin-2-like

MSSWVLVFLFLSVVSLSSSEASPFRNEKKEEGKRERPLHDTIMERMKKIRESLSMKGQDEEYKRYWEELIANNPDLAKRFREQKAEDLAAGLGDRLSHRDGGIRSQMEELQRRNIEELRDLHRQKMAGESHEPIEIRRQRERGRHLSDKEMEEKIREHQRIQEQLDHARREDFKNFEMKLEHRRRQKLKEMDETQRLREEARLKKLSDERKVHEPLLHPASKDQFEEVWKTDDGFKDEKFDIRTFFHLHDTNGDGTLDIREVEALFVNEIRKYYGKDFENDREAHEDMSRMREHVMSEVDRDHDSLITLEEFVRYANTDLFNIKEEWKPVGEDDEHKEFTDDEFKQYENEMEEDDYDYDDEGNLIHKPKMENHDGVPPEHHEVPTEHHEVPPEHQEVPTEHHEVPAAHDTKPDTGLPNDHGLPPNDHGLPNDHGLPPYDHRLPNDHGLPSDHGSPKGEKGLPEHVPPQDVPHDVPVGGANHNENPKAPIVETKQDDTAPPPHAASDDEKKL

>XP_003386431.1 [Amphimedon queenslandica] nucleobindin-2-like

MSSWVLVFLFLSVVSLSSSEASPFRNEKKEEGKRERPLHDTIMERMKKIRESLSMKGQDEEYKRYWEELIANNP--[1]--DLAKRFREQKAEDLAAGLGDRLSHRDGGIRSQMEELQRRNIEELRDLH--[2]--RQKMAGESHEPIEIRRQRERGRHLSDKEMEEKIRE--[0]--HQRIQEQLDHARREDFKNFEMKLEHRRRQKLKEMDETQRLREEA--[2]--RLKKLSDERKVHEPLLHPASKDQFEEVWKTDDGFKDEKFDIRTFFHLH--[1]--DTNGDGTLDIREVEALFVNE--[0]--IRKYYGKDFENDREAHEDMSRMREHVMSE--[0]--VDRDHDSLITLEEFVRYANTDLFNIKEEWKPVGEDDEHKEFTDD--[0]--EFKQYENEMEEDDYDYDDEGNLIHKPKMENHDGVPPEHHEVPTEHHEVPPEHQEVPTEHHEVPAAHDTKPDTGLPNDHGLPPNDHGLPNDHGLPPYDHRLPNDHGLPSDHGSPKGEKGLPEHVPPQDVPHDVPVGGANHNENPKAP--[1]--IVETKQDDTAPPPHAASDDEKKL

>XM_003386383.3 [Amphimedon queenslandica] nucleobindin-2-like (LOC100637379), mRNA

ATGTCCTCTTGGGTTCTTGTGTTTCTGTTTCTATCCGTTGTTTCTTTGTCTTCATCAGAAGCCTCTCCTTTTAGAAATGAGAAGAAAGAAGAAGGAAAAAGGGAGCGTCCATTGCATGATACTATAATGGAGAGGATGAAGAAGATAAGAGAGTCTTTGTCAATGAAAGGACAGGACGAGGAGTACAAGAGGTACTGGGAAGAACTCATCGCTAATAATCCTGACTTGGCTAAGCGGTTTCGTGAGCAGAAAGCTGAAGATCTAGCAGCTGGTCTCGGAGACAGACTCTCTCACCGTGACGGAGGAATCCGGTCTCAAATGGAGGAACTGCAGAGGAGGAACATTGAAGAGTTGAGAGATCTTCACAGGCAAAAAATGGCAGGAGAATCCCATGAGCCCATTGAGATTAGGAGACAGAGAGAGAGAGGGAGACACTTGTCAGATAAAGAAATGGAAGAGAAGATCAGAGAGCATCAAAGGATTCAGGAGCAGTTAGATCACGCTCGTCGTGAAGATTTCAAAAATTTTGAAATGAAACTCGAGCACAGAAGAAGACAGAAATTAAAGGAAATGGACGAAACTCAAAGACTCAGAGAAGAAGCAAGATTAAAGAAGCTATCAGACGAGAGAAAGGTTCATGAGCCATTGCTCCATCCAGCAAGTAAAGATCAGTTTGAGGAGGTATGGAAGACTGATGATGGATTTAAAGATGAAAAGTTCGACATTAGGACATTCTTTCACCTTCACGATACAAATGGAGATGGGACTTTAGACATTCGTGAAGTTGAAGCACTTTTTGTCAATGAAATAAGGAAGTATTATGGTAAGGATTTTGAAAATGACAGAGAAGCTCACGAAGATATGTCAAGAATGAGAGAACACGTTATGAGTGAAGTTGATCGAGATCATGATTCCTTGATCACGTTAGAAGAGTTTGTGAGATACGCTAACACTGATCTGTTCAATATAAAGGAAGAATGGAAGCCAGTTGGGGAGGATGATGAGCACAAGGAGTTTACTGATGATGAATTCAAACAGTATGAAAACGAAATGGAAGAAGATGATTATGATTATGACGACGAAGGTAACCTCATTCATAAACCAAAAATGGAAAACCATGACGGAGTACCCCCTGAACACCACGAAGTACCCACTGAACATCATGAAGTACCTCCTGAGCACCAGGAAGTACCCACTGAACATCATGAAGTACCTGCAGCACATGATACAAAACCTGATACTGGATTACCTAATGATCATGGATTACCTCCTAATGATCACGGGTTACCTAATGATCACGGGTTACCTCCTTATGATCATAGGTTACCTAATGATCATGGATTACCTAGTGATCATGGGTCACCAAAAGGTGAGAAAGGACTGCCCGAGCATGTTCCTCCACAGGATGTGCCTCATGATGTTCCTGTAGGAGGAGCTAATCACAATGAGAACCCAAAAGCACCAATTGTTGAAACTAAGCAAGATGATACTGCTCCTCCTCCTCATGCTGCTTCTGATGATGAAAAGAAATTGTAA

Genomic: JADHEG010001299.1

Salpingoeca rosetta

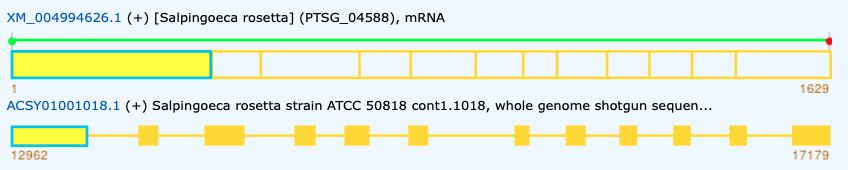

>XP_004994683.1 [Salpingoeca rosetta] PTSG_04588

MRTKQAALLVLALVGVLAVVGCMQGMTGAHAAPVRTAVEDTDTDNAAAADAADAAESTAPPADDAPVDADSAQDTTDAGTDNGAAKPVPVSKDEPYKAYDPYDPYDYDDYDYGEYEELYNEQYDLYADEQLDLKDEESYVGGDEDDDADPDVMSVRKEAAEKRRKADEEEYQRYLDEMMEEMARDSDLREDLIEDIQGVQEAERAPGDKDILKIALRLKNEERVRNIIEEKERQLLEAQRHEARRRAEARMHVPRPHDAMPEGGEEEDEDLFLDPEAAFDKLTEIVKRRAAVLGEIDTKRRAAFLQKEMMRELEFRRSVAHEKDPEKKKQLIKSHKQKIGHAKQLNEVWEKQDGMQSKFNPKVFFSLHDINGDDFWDHKELEAIFHRSAVKLHTIDGETDTDAVEEEMLDMRAWVMKEVDTDKDQLVSRDEFITFANSPEFQTNRDWKAVFPDFNGTELKEFKDRARKAGDPRQEHVQARVAEASKRFKDKFAKTMKLGTNALKEAIKPFAKVGGNKKPAAQGDEGAAANEQQQQQQQQEQK

>XP_004994683.1 [Salpingoeca rosetta] PTSG_04588

MRTKQAALLVLALVGVLAVVGCMQGMTGAHAAPVRTAVEDTDTDNAAAADAADAAESTAPPADDAPVDADSAQDTTDAGTDNGAAKPVPVSKDEPYKAYDPYDPYDYDDYDYGEYEELYNEQYDLYADEQLD--[2]--LKDEESYVGGDEDDDADPDVMSVRKEAAEKRRK--[1]--ADEEEYQRYLDEMMEEMARDSDLREDLIEDIQGVQEAERAPGDKDILKIALRLKNEERVRNIIEEK--[0]--ERQLLEAQRHEARRRAEARMHVPRPHDAMPEGGE--[0]--EEDEDLFLDPEAAFDKLTEIVKRRAAVLGEIDTKRRAAFLQKE--[0]--MMRELEFRRSVAHEKDPEKKKQLIKSHKQKI--[0]--GHAKQLNEVWEKQDGMQSKFNPK--[0]--VFFSLHDINGDDFWDHKELEAIFHRSAVKLHT--[2]--IDGETDTDAVEEEMLDMRAWVMKEVDTDK--[0]--DQLVSRDEFITFANSPEFQTNRDWKAVF--[0]--PDFNGTELKEFKDRARKAGDPRQEHVQAR--[0]--VAEASKRFKDKFAKTMKLGTNALKEAIKPFAKVGGNKKPAAQGDEGAAANEQQQQQQQQEQK

>XM_004994626.1 [Salpingoeca rosetta] (PTSG_04588), mRNA

ATGAGGACGAAGCAAGCGGCGCTGCTTGTGCTGGCGCTTGTTGGCGTGCTGGCTGTCGTGGGCTGCATGCAGGGGATGACTGGTGCACATGCTGCACCCGTCCGCACCGCTGTCGAGGACACGGACACAGACAACGCTGCTGCTGCCGATGCTGCCGATGCTGCCGAGAGCACCGCCCCACCTGCAGATGATGCGCCAGTGGACGCAGACAGCGCGCAAGACACGACCGACGCTGGCACAGACAACGGCGCAGCCAAGCCTGTTCCCGTGAGCAAGGATGAGCCCTACAAGGCCTACGACCCGTACGACCCTTACGACTACGACGACTATGACTACGGAGAGTATGAGGAGCTGTACAACGAGCAATATGACCTCTACGCCGACGAGCAGCTCGACCTGAAGGATGAAGAAAGCTATGTTGGCGGTGATGAGGATGACGACGCAGACCCAGATGTCATGTCCGTGCGCAAGGAAGCTGCCGAGAAGCGCAGAAAGGCCGATGAGGAAGAGTATCAGCGGTATCTTGACGAGATGATGGAGGAGATGGCGCGTGACTCAGACCTCCGTGAAGACCTCATTGAAGACATTCAGGGGGTGCAGGAGGCCGAGCGGGCGCCCGGCGATAAGGACATTCTCAAGATTGCCCTTCGCCTCAAGAACGAGGAGCGCGTGCGGAACATCATCGAGGAGAAGGAGCGGCAGTTGCTTGAGGCGCAGCGCCACGAAGCGCGACGGCGGGCAGAGGCGCGGATGCATGTTCCGCGGCCGCATGATGCAATGCCAGAAGGCGGAGAGGAGGAAGACGAGGATCTGTTTCTTGACCCCGAGGCCGCGTTTGACAAACTTACCGAGATTGTCAAACGCAGGGCCGCTGTTCTCGGCGAAATCGACACGAAGCGCCGTGCAGCATTCTTGCAGAAGGAGATGATGCGCGAGTTGGAGTTTCGGCGCTCCGTTGCGCACGAGAAAGATCCCGAGAAGAAGAAGCAGCTCATCAAGAGCCACAAGCAAAAGATCGGTCACGCCAAGCAGCTGAATGAGGTGTGGGAGAAGCAGGACGGCATGCAGTCGAAGTTCAACCCAAAGGTGTTCTTTTCGCTGCACGACATCAACGGCGACGATTTCTGGGACCACAAGGAGCTGGAGGCCATCTTCCACCGCTCAGCCGTCAAGCTGCACACCATCGACGGGGAGACGGATACGGATGCCGTTGAAGAGGAGATGCTCGACATGCGCGCTTGGGTCATGAAGGAGGTCGACACAGACAAGGATCAGCTTGTGTCACGTGACGAGTTCATCACATTTGCAAACTCGCCCGAGTTCCAGACCAACCGTGACTGGAAGGCCGTCTTCCCGGACTTCAACGGCACGGAGCTGAAGGAGTTCAAGGACCGCGCGAGAAAGGCAGGCGACCCGCGCCAGGAACACGTCCAAGCACGCGTTGCTGAGGCGAGCAAGCGGTTCAAGGACAAGTTTGCGAAGACGATGAAGCTCGGCACCAACGCGCTCAAGGAGGCCATCAAGCCGTTTGCGAAGGTCGGCGGCAATAAGAAGCCGGCTGCTCAGGGCGATGAGGGCGCCGCTGCCAACGAACAACAACAGCAGCAGCAGCAACAGGAGCAGAAGTGA

Genomic: ACSY01001018.1
