## Supplementary figures and images for "Pre-metazoan origin of neuropeptide signalling"

### Supplementary Figures 2 and 3

Color Align Conservation results

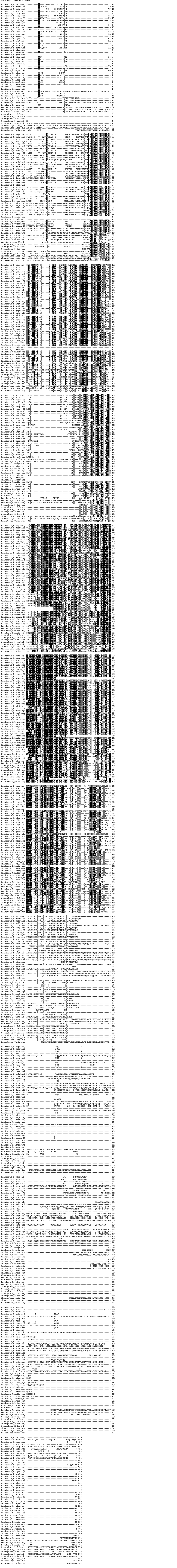
